# Towards a Neurobiologically-driven Ontology of Mental Functions: A Data-driven Summary of the Twenty Years of Neuroimaging Meta-Analyses

**DOI:** 10.1101/2023.03.29.534795

**Authors:** Jules R. Dugré, Stéphane Potvin

**Author notes:** **Corresponding authors** Jules Roger Dugré, PhD Candidate & Stéphane Potvin, PhD; Research Center of the Institut Universitaire en Santé Mentale de Montréal; 7331 Hochelaga, Montreal, Quebec, Canada; H1N 3V2; /.

## Abstract

A persistent effort in neuroscience has been to pinpoint the neurobiological substrates that support mental processes. The Research Domain Criteria (RDoC) aims to develop a new framework based on fundamental neurobiological dimensions. However, results from several meta-analysis of task-based fMRI showed substantial spatial overlap between several mental processes including emotion and anticipatory processes, irrespectively of the valence. Consequently, there is a crucial need to better characterize the core neurobiological processes using a data-driven techniques, given that these analytic approaches can capture the core neurobiological processes across neuroimaging literature that may not be identifiable through expert-driven categories. Therefore, we sought to examine the main data-driven co-activation networks across the past 20 years of published meta-analyses on task-based fMRI studies. We manually extracted 19,822 coordinates from 1,347 identified meta-analytic experiments. A Correlation-Matrix-Based Hierarchical Clustering was conducted on spatial similarity between these meta-analytic experiments, to identify the main co-activation networks. Activation likelihood estimation was then used to identify spatially convergent brain regions across experiments in each network. Across 1,347 meta-analyses, we found 13 co-activation networks which were further characterized by various psychological terms and distinct association with receptor density maps and intrinsic functional connectivity networks. At a fMRI activation resolution, neurobiological processes seem more similar than different across various mental functions. We discussed the potential limitation of linking brain activation to psychological labels and investigated potential avenues to tackle this long-lasting research question.

## INTRODUCTION

Modern biological theories posit that mental functions could be organized by distinct neurobiological systems which interact with each other (Cloninger, 1987; Eysenck, 1981; Gray, 1970; Jaak Panksepp, 1998). Indeed, Panksepp described seven core neurobiological systems which mirrored *emotional feelings*, all characterized by subcortical structures developed in the mammalian brain. Examples of such systems include **rage** (e.g., medial amygdala, hypothalamus & dorsal periaqueductal grey [PAG]), **fear** (e.g., basolateral and central amygdala, bed nucleus of the stria terminalis, anterior and medial hypothalamus and PAG), **seeking** (e.g., ventral midbrain, nucleus accumbens and medial frontal cortex) (Panksepp, 2010; Panksepp, Fuchs, & Iacobucci, 2011). In the past decades, research has described the neurobiological underpinnings of various mental processes such as executive functions (Corbetta & Shulman, 2002; Norman & Shallice, 1986; Petersen & Posner, 2012; Smith & Jonides, 1999), language (Price, 2000) and memory (Squire, 2004). Since the development of functional neuroimaging, which allows us to dive right into the activity of the human brain, there is an exponential demand for *validating* and *characterizing* neuroimaging results through their most likely psychological labels. These attempts largely reflect the human need to categorize and discriminate mental processes (at a psychological level) through their neurobiological substrates. Yet, the identification of reliable brain circuits remains one key challenge in neuroscience (Rust et LeDoux, 2022).

In 2010, the National Institute of Mental Health has developed an exhaustive classification of neurocognitive systems which are thought to vary in terms of units of analysis (e.g., genes, molecules, cells, brain circuits) (Cuthbert, 2015; Insel et al., 2010; Kozak & Cuthbert, 2016). This expert-driven classification is of utmost importance to better understand specific and shared dimensions across psychiatric disorders. The six neurocognitive domains include Negative Valence Systems, Positive Valence Systems, Cognitive Systems, Social Processes, Arousal and Regulatory Systems (Kozak & Cuthbert, 2016), and more recently the Sensorimotor Systems (National Advisory Mental Health Council Workgroup on Changes to the Research Domain Criteria Matrix, 2018). However, this classification still lack validation across neuroimaging literature. Indeed, several neurocognitive processes exhibit substantial overlaps in brain their respective brain circuits, raising question about the validity of such classification system. For example, past meta-analyses of fMRI studies on reward anticipation (Diekhof, Kaps, Falkai, & Gruber, 2012; Knutson & Greer, 2008; Liu, Hairston, Schrier, & Fan, 2011; Oldham et al., 2018; Wilson et al., 2018) and loss/punishment processing (Dugré, Dumais, Bitar, & Potvin, 2018; Wilson et al., 2018) showed that both reliably involve the ventral striatum, amygdala, ventral tegmental area, thalamus, insula and anterior midcingulate cortex/pre-supplementary motor area. Indeed, the most recent meta-analysis found no significant differences in the recruitment of such regions between reward and loss anticipation (Chen, Chaudhary, & Li, 2022). Rather than what was previously theorized, findings from meta-analyses on positive emotional stimuli (Lindquist, Satpute, Wager, Weber, & Barrett, 2016; Mende-Siedlecki, Said, & Todorov, 2013; Stevens & Hamann, 2012) and negative emotional stimuli (Lindquist et al., 2016; Pozzi, Vijayakumar, Rakesh, & Whittle, 2021; Ran, Cao, & Chen, 2018; Stevens & Hamann, 2012; Tao, He, Lin, Liu, & Tao, 2021; Vytal & Hamann, 2010) suggest that most of these regions appear to be valence-general (Lindquist et al., 2016). Spatial commonalities have also been observed across cognitive systems despite strong neuropsychological distinction. Despite that early evidence supported that shifting, updating, and inhibition were neuropsychologically distinct (Miyake et al., 2000, see also Karr et al., 2018) for a recent discussion about the latent structure), results from neuroimaging literature suggest that they show activation of common brain regions including frontal (Duncan & Owen, 2000; Niendam et al., 2012), and parietal regions (Niendam et al., 2012). While these tasks are delineated at a (neuro)psychological level, their respective brain circuits may be more similar than different at a BOLD fMRI resolution. Yet, the question remains largely unanswered: what are the main homogeneous brain circuits across fMRI tasks?

In the past decade, several authors aimed to tackle limitations surrounding expert-driven ontologies of mental processes using data-driven methods (Beam, Potts, Poldrack, & Etkin, 2021; Bolt et al., 2020; Laird et al., 2011; Nakai & Nishimoto, 2020; Poldrack & Yarkoni, 2016; Ray et al., 2013; Smith et al., 2009). Indeed, data-driven analytic techniques allows to capture neurobiological processes that may not be identifiable through expert-driven categories. Recently, Beam and colleagues (2021) have conducted a data-driven meta-analysis to better link brain structures and psychological constructs using approximately 20,000 neuroimaging studies. They observed that mental functions, at the lowest level of neurobiological specificity, may be mainly organized and characterized by **vision** (e.g., lingual gyrus, lateral occipital cortex, fusiform gyrus), **memory** (e.g., amygdala, hippocampus, parahippocampal gyrus), **reward** (e.g., striatum, orbitofrontal cortex, thalamus, brainstem), **cognition** (e.g., frontal pole, insula, cingulate cortex), **manipulation** (e.g., middle frontal gyrus, premotor, supplementary motor area, precentral/postcentral gyri, cerebellum) and **language** (e.g., superior and middle temporal gyri and heschl’s gyrus) systems. In contrast to Beam et al., (2021)’s structure-based approach (i.e., circuits are identifiable via non-overlapping sets of brain structures), Bolt and colleagues (2020) used a graph clustering approach on 8919 parcellated activation maps (i.e., 70 parcels) which allowed overlapping communities. They observed that BOLD activation maps could be optimally mapped onto 4 clusters (followed by the 7-cluster solution): **Object Viewing**, **Higher-Order Cognition**, **Affect/Introspection/Social**, and **Auditory-Motor**. While these studies provide new insight about the similarity and differences between brain circuits, they challenge current expert-driven classification of mental functions.

One difficulty faced when identifying neurobiological processes in neuroimaging is that studies usually contain small sample size which significantly increase the spatial uncertainties about the *presumed true* location of the effect (Eickhoff et al., 2009) and consequently alters the reliability of whole-brain activation maps (Turner, Paul, Miller, & Barbey, 2018). In addition, differences in methodological approaches may also partially explain discrepancies between results of fMRI studies (Botvinik-Nezer et al., 2020). While task-based neuroimaging suffers a reliability crisis, meta-analytic approaches may be a promising avenue to identify reliable neurobiological processes (Kim, Knodt, & Hariri, 2022; Wager et al., 2013). In the last decade, there has been an increasing number of published meta-analyses of fMRI studies which comprised distinct behavioral tasks and stimuli, bibliography, imaging modality, methodological approaches (e.g., statistical thresholding) and stereotaxic space (Tahmasian et al., 2019; Yeung, Robertson, Uecker, Fox, & Eickhoff, 2023). Recent advancements in data-driven methods have facilitated the analysis of large-scale neuroimaging datasets, allowing valuable insights of the brain activity patterns underpinning mental functions. Such data-driven approaches have recently been applied to meta-analytic data to uncover main co-activation patterns across studies on reward processing (Flannery et al., 2020), emotional processing (Riedel et al., 2018), face processing (Laird et al., 2015), naturalistic paradigms (Bottenhorn et al., 2019) as well as aberrant co-activation patterns across pediatric psychiatric disorders (Dugré, Eickhoff, & Potvin, 2022). While the classical approach aims to meta-analyze studies sharing a similar construct at a psychological level, it remains largely unknown whether these constructs are characterized by common or distinct brain circuit. As such, applying data-driven methods to meta-analytic data may be critical to clarify the similarity and differences in brain circuits characterizing mental functions.

Consequently, we aimed to summarize the last 20 years of meta-analyses of task-based neuroimaging studies by using a data-driven approach to identify the main coactivation patterns across mental processes. Here, we conducted a correlation-based hierarchical clustering on 1,347 meta-analytic experiments and extracted spatially convergent brain activation maps. To identify reliable psychological labels characterizing the resulting co-activation networks, we used different approaches: the labels attached by the authors of the original meta-analysis and spatial correlation with Neurosynth (Yarkoni, Poldrack, Nichols, Van Essen, & Wager, 2011), and NeuroQuery (Dockès et al., 2020) databases. In addition to the psychological labelling, we aimed to validate the data-driven co-activation maps by assessing their relationships with 19 receptor density maps (Hansen et al., 2022) and well-defined intrinsic functional connectivity networks (7- & 17-networks, (Schaefer et al., 2018). “As a form of convergent validity, we compared our co-activation networks to the data-driven voxelwise maps reported by Bolt and colleagues (Bolt et al., 2020).

## 2. METHODS

### 2.1. Identification of included meta-analyses

Our search focused primarily on meta-analyses published up to December 2021^st^ which was mainly retrieved on the publication list of the BrainMap (Yeung et al., 2023) (see https://brainmap.org/pubs/). Given that several other meta-analytic methods exist such as SDM and MKDA, we also conducted a non-systematic literature search using Pubmed and Google Scholar. Relevant meta-analyses were found by keyword search: “meta-analysis”, “fMRI”, “neuroimaging”, “cognition”, “emotion”, “reward”, “punishment”, “social cognition”, “perception”, “visual”, “motor”, “memory”, “audition”. Further meta-analyses were identified through cross-referencing. Inclusion criteria were: (1) original manuscript from a peer-reviewed journal, (2) task-based functional MRI meta-analyses, (3) use of a whole-brain methodology (i.e., meta-analyses using ROIs were excluded) irrespectively of the task contrasts, (4) Healthy Subjects with no reported psychiatric disorders. Only main contrasts (i.e., Single dataset analysis such as Condition > Control or Fixation) were included. Comparisons between two meta-analytic maps (i.e., contrasts analyses such as Meta-analysis 1 *versus* Meta-analysis 2) were excluded. Coordinates of meta-analytic experiments were manually extracted. Coordinates that were reported originally in Talairach stereotaxic space were converted into MNI (Montreal Neurologic Institute) space.

### 2.2. Data-Driven Meta-Analysis

#### 2.2.1. Correlation-Matrix-Based Hierarchical Clustering

Modeled activation (MA) map was created for each meta-analytic experiment (2mm^3^ resolution), converted into a 1D feature vector of voxel values (i.e., 2mm^3^ grey matter mask in MNI space), and then concatenated together to form an experiment (*e*) by voxel (v) matrix (i.e., 1,347 meta-analytic experiments by 226,654 voxels). Then, pairwise Spearman’s rank correlation was performed between the 1D vectors to obtain spatial similarity between maps (Kriegeskorte, Mur, & Bandettini, 2008). We thus identified meta-analytic experiments that showed similar brain topographic map, through a Correlation-Matrix-Based Hierarchical Clustering (Jules R. Dugré, Simon B. Eickhoff, & Stéphane Potvin, 2022; Laird et al., 2015; Michael C Riedel et al., 2018). The hierarchical clustering was carried out using rank-correlation distance (1–*r*) and complete linkage method. More importantly, the most optimal number of clusters was found by assessing the silhouette, calinski-harabasz indices, and adjusted rand indices for a *k* range of 2 to 20 clusters (Eickhoff, Thirion, Varoquaux, & Bzdok, 2015) (See Supplementary Material). More precisely, at each *k*, metrics were compared against a null distribution of random spatial arrangement. To do so, 2,500 datasets were created artificially by shuffling foci locations across meta-analyses but preserving original meta-analyses’ properties (e.g., number of foci, sample size). The average of each metric (i.e., Silhouette and Calinski-Harabasz indices) derived from true dataset, was compared against the artificially created null distribution and then plotted for *k* range of 2 to 20 clusters. This improves our ability to select the most optimal *k* by considering the probabilities of getting a certain metric value in a random spatial arrangement. Given that the ground truth class labels are unknown, we compared, for each *k*, the consistency (adjusted rand index) between Label_TRUE_ with the Label_NULL_ then averaged across the 2500 iterations. A local minimum in the plots suggests a decrease in overlap between both sets of labels. After having found the optimal number of clusters, we randomly removed 10% of the meta-analyses and re-ran the clustering algorithm 10,000 times. we calculated the hamming distance between each experiment’s label ids (1,347 meta-analyses by 10,000 iterations ids) to calculate the proportion of disagreement between two meta-analyses’ set of labels. This was done to select the most stable labeling solution for a final hierarchical clustering. All these analyses were performed using Scikit-learn (version 0.21.3) in Python (version 3.7.4) (Pedregosa et al., 2011).

#### 2.2.3. Meta-Analytical Groupings

An activation likelihood estimaton (ALE) algorithm was then used to identify spatially convergent brain regions in each of the resulting groups of experiments (GingerALE version 3.0.2) (Eickhoff, Bzdok, Laird, Kurth, & Fox, 2012; Simon B Eickhoff et al., 2009). Voxel-wise ALE scores were computed as the union of MA maps, which provide a quantitative assessment of spatial convergence across experiments. These voxel-wise maps were cut off by a cluster-forming threshold. In fact, the size of the supra-threshold clusters was compared against a null distribution of cluster sizes derived from artificially created datasets in which foci were shuffled across experiments, but the other properties of original experiments (e.g., number of foci, uncertainty) were kept (Eickhoff et al., 2012). Considering the large sample size and the presence of meta-analysis, we used the following stringent statistical threshold: a voxel-level threshold of p<0.0001 and a cluster-level family-wise correction (cFWE) based on the cluster mass, with 10,000 permutations (Eickhoff et al., 2016) using the Neuroimaging Meta-Analysis Research Environment package for python (NiMARE) ((Salo et al., 2022).

### 2.3. Functional Characterization

#### Psychological Labels

We carried out additional analyses to examine spatial similarities between the resulting co-activation networks and psychological terms. First, we identified the 10 most contributing meta-analytic experiments for each co-activation networks. Terms attached to the meta-analytic experiments were extracted from the original author’s label of the contrast of interest. Second, we characterized our meta-analytic networks using two distinct automated meta-analytic decoding based on spatial similarity: Neurosynth term-based decoding (Yarkoni et al., 2011) and NeuroQuery Image Search (Dockès et al., 2020). Although these two methods focus on spatial similarity, they both differ in their method to extract terms from documents (i.e., abstracts [NeuroSynth] versus full article [NeuroQuery]) as well as in their analytic approaches. Briefly, Neurosynth method calculates the pearson’s correlation coefficient across voxels of two maps (input and automated meta-analytic images). The NeuroQuery method rather utilizes normalized (divided by L2 norm) sets of reduced brain maps (DiFuMo atlas rather than voxelwise maps), calculate the scalar product of two vectors (*input* and *term*), and weights the similarity by the log of proportion of articles in the corpus in which the term appears. The top 10 most strongly associated maps were extracted and used to generate a Wordcloud.

#### PET Density Maps & Intrinsic Functional Connectivity Networks

We sought to examine how our Co-activation Networks (CN) may be related with each other based on neurochemical substrates (i.e., receptor and transporter density). First, we compared our meta-analytic maps with 19 whole-brain receptor/transporter density maps described in (Hansen et al., 2022) which are distributed across 9 neurotransmitter systems including serotonin (i.e., 5-HT_1A_ , 5-HT_1B_ , 5-HT_2A_ , 5-HT_4_ , 5-HT_6_ , 5-HTT), dopamine (i.e., D_1_ , D_2_ , DAT), norepinephrine (i.e., NET), Histamine (i.e., H_3_), acetylcholine (i.e., α4β2, M_1_ , VAChT), cannabinoid (i.e., CB_1_), opioid (i.e., MOR), glutamate (i.e., NMDA, mGluR_5_) and GABA (i.e., GABA_A/BZ_). Spatial associations between the CNs and the PET density maps were conducted using JuSpace (version 1.4) (Dukart et al., 2021). Briefly, the mean values of 728 brain regions which included 690 parcels of the Yale Brain Atlas (McGrath et al., 2022) and additional 38 subcortical and cerebellar regions from the AAL3 atlas (Rolls, Huang, Lin, Feng, & Joliot, 2020), were extracted for the two images. Partial correlation (Spearman’s rank correlation) adjusting for spatial autocorrelation (i.e., local grey matter probabilities) was then performed between the two sets. Exact permutation-based p-values (with 10,000 permutations) were computed and corrected using false discovery rate (FDR).

In addition, we aimed to compare how our data-driven coactivation maps were related to well-defined resting-state intrinsic networks using the 7- & 17-networks from Shaefer’s 400 cortical parcels (Schaefer et al., 2018). We computed the percentage of voxels from the thresholded (p<0.0001, cFWE<0.05) binarized coactivation maps that overlapped with those from the intrinsic networks.

#### Spatial Similarity and Overlaps across the Co-activation Networks

After having identified and described the CNs, we conducted pairwise spearman’s rank correlation across voxels of the unthresholded maps to examine their spatial similarity. Doing so allowed us to validate our clustering solutions and describe the similarity in activation patterns between the co-activation networks. Furthermore, we examine which voxels were the most activated across the networks (spatial overlap). This was done by spatially overlapping the thresholded and binarized maps (p<0.0001, cFWE<0.05). Voxels with larger values (up to 13) suggest activation in a greater variety of brain circuits (i.e., non-specific). Their corresponding co-activation networks were extracted.

#### Comparisons with previous Data-Driven Ontology of Brain Circuits

We sought to compare the resulting CNs with those from previous data-driven framework (Bolt et al., 2020). For instance, Bolt et al. (2020)’s used a Symmetric non-negative matrix factorization (Sym-NMF) on BrainMap data to generate their data-driven maps. Spatial similarity was examined using the same method as presented above to compare the identified CNs (unthresholded z-maps) and Bolt et al., (2020)’s voxelwise activation probability maps. We chose their 7-cluster solution, *a posteriori*, to approximate the level of neurobiological specificity (i.e., number of data-driven maps) found in our study.

## 3. RESULTS

### 3.1. Data-Driven Synthesis of fMRI Meta-analyses

A total of 1,347 meta-analytic experiments met the inclusion criteria which comprised 19,822 foci (see Supplementary Material for the list of included studies). Clustering solutions were investigated for a range of *k*=2-20 cluster solutions (see Supplementary Material). Silhouette coefficient showed that k=4, k=8, k=13, k=17 were the best solutions. Calinski-Harabasz index rather exhibited a monotonic increase. However, k=13 & k=9 both showed greatest increase from previous (k-1) and following (k+1) solutions). Adjusted Rand Indices showed that the clustering solutions were the most highly discordant from random spatial arrangement at k=4 and k=14. Also, aRI values were negative at k=2, k=3, and k=11 to k=20, suggesting strong dissimilarity between these solutions and random arrangements. Overall, these metrics indicated that the 13-cluster solution was the most optimal solution. The 13 voxelwise brain maps (i.e., z-score unthresholded & thresholded/binarized maps, are available on NeuroVault: https://neurovault.org/collections/13769/).

#### CN1 – Multi-Demand (219 meta-analytic experiments)

The Multi-Demand CN included meta-analytic experiments on Timing (Teghil et al., 2019), Subtraction (Arsalidou & Taylor, 2011), Stroop Task (Cieslik, Mueller, Eickhoff, Langner, & Eickhoff, 2015), Incongruent (Stroop) (Huang, Su, & Ma, 2020), Stimulus-Stimulus Conflict (Q. Li et al., 2017), Resolution Interference (Nee, Wager, & Jonides, 2007), Verbal Stroop (Laird et al., 2005), Inhibition (Wu et al., 2020), Grammar Learning (Tagarelli, Shattuck, Turkeltaub, & Ullman, 2019), Task Switching (Worringer et al., 2019). Main activation comprised bilateral insula, anterior midcingulate/pre-supplementary motor area, bilateral basal ganglia, dorsolateral profrontal cortex, middle frontal gyrus, supramarginal/angular gyri, posterior fusiform and superior parietal lobule (see Figure 2 & Supplementary Material for complete lists of regions and their corresponding MNI coordinates).

**Figure 1.**
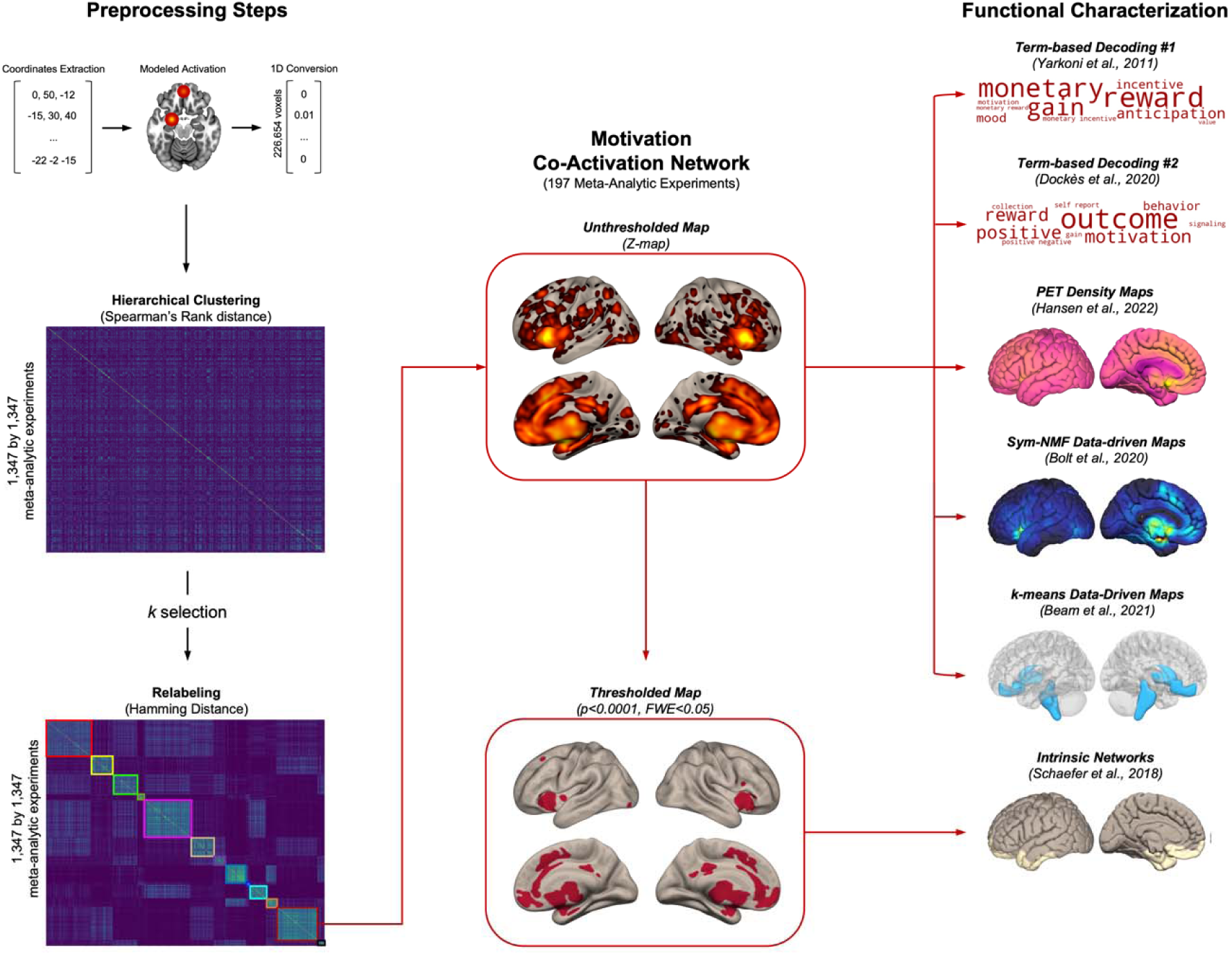
Summary of the Data-Driven Meta-analytic Groupings & Functional Characterization Steps followed in this study included (1) extracting coordinates from original meta-analytic’ studies, (2) creating a modeled activation map (i.e., spherical Gaussian distribution around peak coordinates), (3) converting these maps to a 1D feature vector, (4) concatenating the modeled activation maps yielding a 1,347 meta-analytic experiments by 226,654 voxels matrix, (5) applying a hierarchical clustering on distance measure (1 – Spearman’s rank correlation) and comparing cluster solution to random spatial arrangement for a range of k=2 to 20 clusters, (6) finding the most stable solution with a subsampling method and computing Hamming distanc between cluster’s labels, as used previously (Dugré, Eickhoff et Potvin, 2022). Experiments within each groups of experiments were meta-analyzed with ALE method to extract their spatially convergent brain activation maps. Functional characterization included spatial similarity with psychological terms according to NeuroSynth (Yarkoni et al., 2011) and NeuroQuery (Dockès et al., 2020); with 19 PET density maps (Hansen et al., 2022), the data-driven 7-cluster solution provided by Bolt and colleagues (2020) and 7- & 17-Intrinsic Functional Connectivity Network (400 parcels, Schaefer et al., 2018)

**Figure 2.**
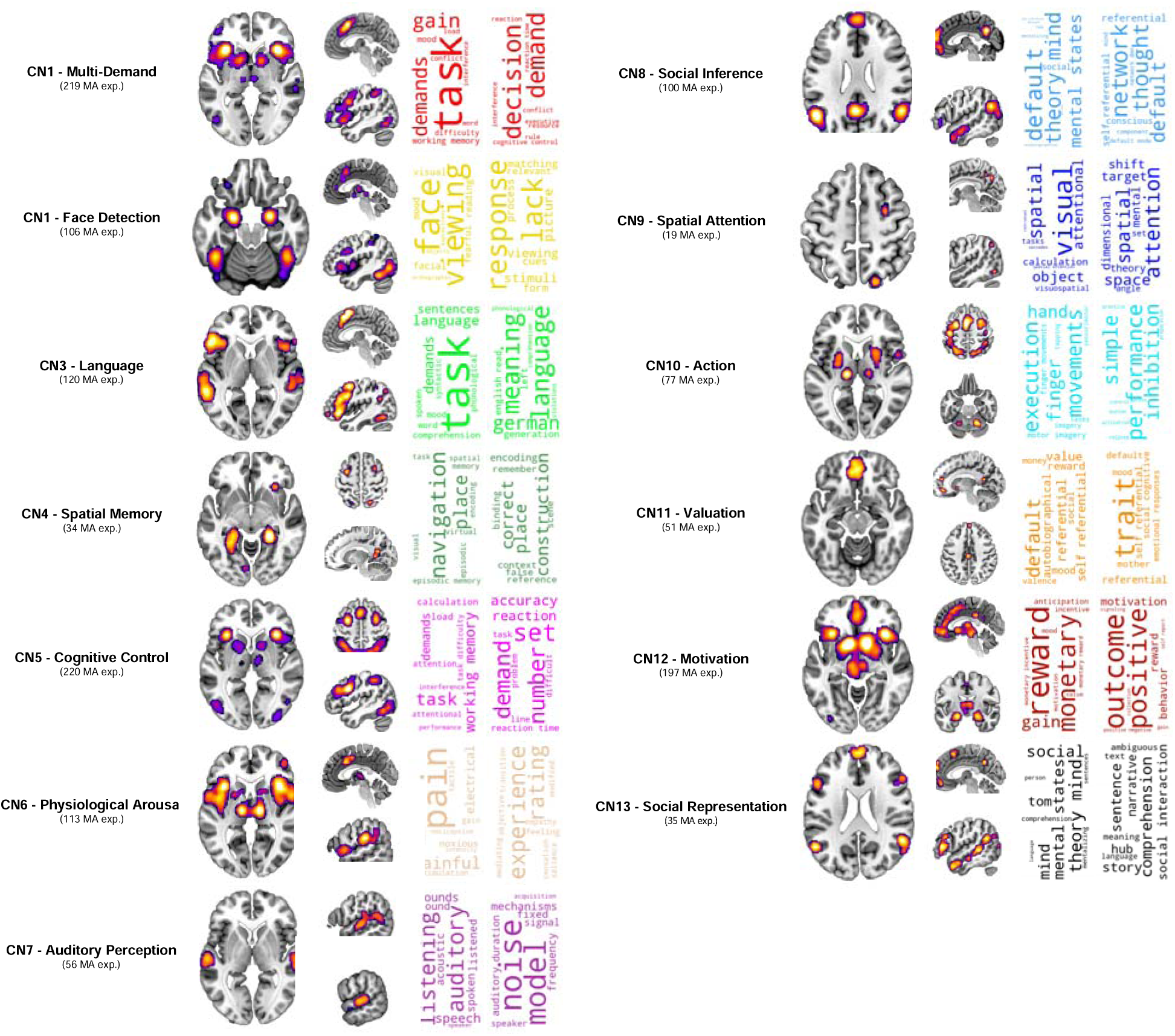
Spatially convergent brain regions for each of the identified co-activation networks (p=0.0001 at a voxel-level, cFWE<0.05). Wordclouds represent the top 10 psychological terms most strongly associated with the unthresholded co-activation networks (z-maps). Wordclouds were generated with the Neurosynth (Yarkoni et al., 2011) and NeuroQuery (Dockès et al., 2020) methods, respectively. MA = Meta-analytic experiments.

As shown in Figure 2, functional characterization indicated that this coactivation network is associated with a widespread of cognitive terms including demand, load, cognitive control, conflict monitoring, cognitive interference (see Figure 2). Spatial overlap with the intrinsic functional connectivity networks suggested relatively strong overlap in voxels with the FP (32.8%) & the VentAttn (32%) networks (see Figure 3). More precisely, overlap was stronger in FP-A (21.25%), VentAttn-B (20.89%), followed by VentAttn-A (16.57%) networks. Finally, this network was mainly associated with CB_1_ (*r*=.57), 5-HT_1B_ (*r*=.48), mGluR5 (*r*=.47), H3 (*r*=.46) density maps after FDR correction.

**Figure 3.**
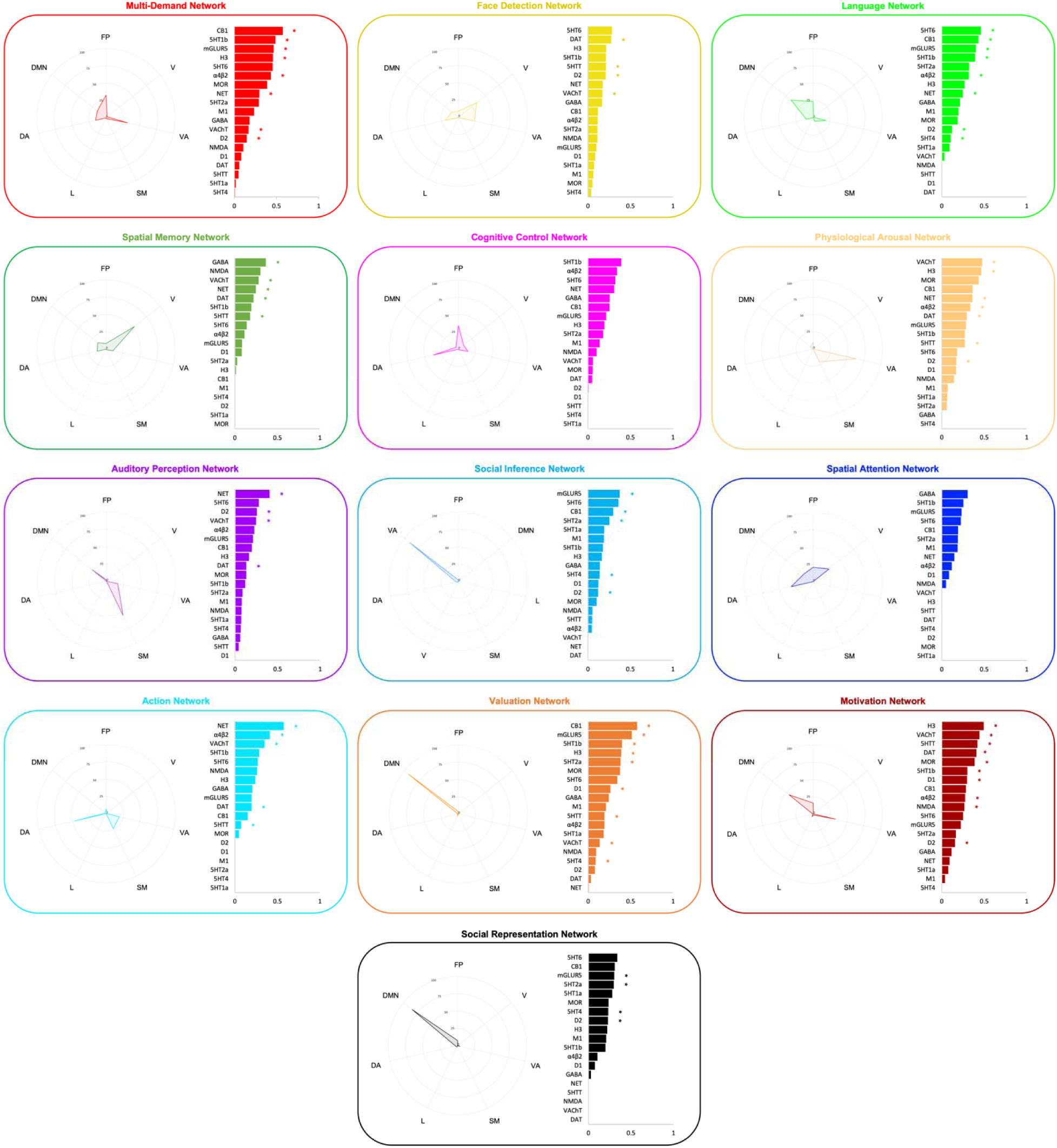
Spatial similarity between the data-driven co-activation networks and intrinsic functional connectivity networks (Schaefer et al., 2018) and PET receptor/transporter density maps (Hansen et al., 2022). Spatial similarity was computed using percentage of voxels’overlap for the intrinsic functional connectivity networks and Fisher’s z (Spearman’s Rank Correlation) for the PET density maps. FP = Frontoparietal Network; VA = Ventral Attention Network; DA = Dorsal Attention Network; V = Visual Network; SM = Somatomotor Network; L = Limbic, and DMN = Default Mode Network. * = pFDR<0.05

#### CN2 – Face Detection (106 meta-analytic experiments)

The Face Detection CN included meta-analyses on Visual Faces (Satpute et al., 2015), Negative Faces (García-García et al., 2016), Face Processing (Müller, Höhner, & Eickhoff, 2018), Emotion Structure (Kober et al., 2008), Emotion (Adolfi et al., 2017), Empathy (Diveica, Koldewyn, & Binney, 2021), Vision (Aesthetic) (Brown, Gao, Tisdelle, Eickhoff, & Liotti, 2011), Empathy (Van’t Hooft et al., 2021), Pain Empathy (Jauniaux, Khatibi, Rainville, & Jackson, 2019), and Empathy (Ding et al., 2020). Main activation comprised the bilateral amygdala, bilateral posterior fusiform, precentral gyrus, aMCC/pre-SMA, anterior Supramarginal gyrus, Superior Parietal Lobule, Middle Temporal Gyrus, Superior Colliculus, perigenual ACC and the left thalamus.

Functional characterization using Neurosynth suggests strong correspondence with visual (i.e., viewing, visual), facial (i.e., face, facial, expressions) and emotional (i.e., fearful, mood) terms, whereas NeuroQuery yield more visual (i.e., viewing, stimuli, picture, cues) decoding terms (i.e., matching, form, relevant, process). Voxels of the Face Detection network moderately overlap with those in the Visual (34.2%), VentAttn (23.4%) and the DorsAttn (19.9%). However, relationship with the 17 networks rather revealed stronger overlap with DorsAttn-A (24.7%), Visual-A (21.7%) and VentAttn-A (14.5%) networks. Finally, this network was significantly associated with DAT (*r*=.27), 5-HTT (*r*=.21), D_2_ (*r*=.21), & VAChT (*r*=.17) density maps after FDR correction.

#### CN3 – Language (120 meta-analytic experiments)

Meta-analyses forming this co-activation network mainly included Explicit Evaluation of Voices Expressions (Dricu & Frühholz, 2016), Difficult Speech (Adank, 2012), Passive Listening to Speech (LaCroix, Diaz, & Rogalsky, 2015), Language Processing (Ferstl, Neumann, Bogler, & von Cramon, 2008), Sign Language Comprehension (Trettenbrein, Papitto, Friederici, & Zaccarella, 2021), Cognitive Emotion Regulation (Langner, Leiberg, Hoffstaedter, & Eickhoff, 2018), Action Withholding (Zhang, Geng, & Lee, 2017), Theory of Mind in Adults (Lee, Walker, Hale, & Chen, 2017), Stop Signal (Puiu et al., 2020), Action Stopping (Rae, Hughes, Weaver, Anderson, & Rowe, 2014). The Language CN included the aMCC/pre-SMA, posterior part of the superior and middle temporal gyrus (i.e., Wernicke’s area), fronto-insular cortex (including Broca’s area), middle frontal gyrus, angular gyrus, and superior parietal lobule. This map was left lateralized, although activation in several regions were also observed in the right hemisphere (e.g., inferior and middle frontal gyri and angular gyri).

Functional characterization using Neurosynth and NeuroQuery both revealed strong associations with language and semantic terms (i.e., language, sentences, comprehension, word, generation, phonological). Main overlaps between voxels of this coactivation network and those from the intrinsic functional connectivity networks were mainly observed with the DMN (40.8%), FP (23.2%) and VentAttn (18.9%). This was mainly due to stronger overlaps with DMN-B (29.5%), FP-A (16.4%), and TempPar (13.12%) networks. Finally, this map was spatially associated with 5-HT_6_ (*r*=.46), CB_1_ (*r*=.43), mGluR_5_ (*r*=.40), and 5-HT_1B_ (*r*=.39) receptor density maps.

#### CN4 – Spatial Memory (34 meta-analytic experiments)

Meta-analytic experiments forming this co-activation network mainly included Spatial Navigation (Encoding) (Kühn & Gallinat, 2014), Spatial Navigation (Li et al., 2021), Landmark (Spatial Information) (Qiu et al., 2019), Environmental Navigational Space (Li et al., 2021), Spatial Navigation (Retrieval) (Kühn & Gallinat, 2014), Visuospatial Problem Solving (Bartley et al., 2018), Route-Based Navigational Strategies (Qiu et al., 2019), Large-Scale Spatial Abilities (Men) (Yuan et al., 2019), Scenes Repetition Suppression (Kim, 2017), and Spatial Navigation (Spreng, Mar, & Kim, 2009). Spatial memory was mainly represented by activity in the posterior hippocampus/parahippocampal gyrus, the retrosplenial cortex, supplementary motor area, frontal eye fields, anterior insula, intraparietal sulcus, superior parietal lobule, MT+, and the lingual gyrus.

Functional characterization with both NeuroSynth and NeuroQuery methods revealed association with navigation (i.e., place, context, scene) and episodic memory (i.e., encoding, remember). Voxels from this co-activation largely overlaps with those from the visual network (52.6%), followed by those from the DMN (15.1%). Associations with the 17 networks revealed stronger overlap with Visual-B (27.7%), DMN-C (27.7%), followed by the DorsAttn-A (14.5%) networks. Spatial Memory network was significantly associated with GABA_A/BZ_ (*r*=.37) and VAChT (*r*=.28) receptor as well as NET (*r*=.25), DAT (*r*=.22), and 5-HTT (*r*=.18) transporter density maps, after applying FDR correction.

#### CN5 – Cognitive Control (220 meta-analytic experiments)

The Cognitive Control network was mainly formed by meta-analysis on Working Memory (Rottschy et al., 2012), Uncertainty (Cognition) (Wu et al., 2020), N-Back (Mencarelli et al., 2019), Executive Processing (Niendam et al., 2012), Cognitive Control (Cieslik et al., 2015), Calculation Tasks (Arsalidou & Taylor, 2011), N-Back in Young Adults (Yaple, Stevens, & Arsalidou, 2019), Working Memory (Lee & Xue, 2018), Updating in Children (McKenna, Rushe, & Woodcock, 2017), N-Back (Wang, He, et al., 2019). It comprised activation of the anterior insula, ventrolateral prefrontal cortex, middle frontal gyrus, paracingulate sulcus/pre-SMA, supramarginal gyrus, caudate, thalamus, fusiform gyrus.

Functional characterization revealed association with various cognitive terms including task demand, task difficulty as well as working memory, calculation, accuracy and reaction. Examining voxels overlaps with those from the 7-networks revealed that voxels of this co-activation network moderately overlapped with those from the DorsAttn (36.9%), FP (34.1%), followed by those from the VentAttn (14.1%). More precisely, this was mainly due to stronger overlap with FP-A (28.9%), DorsAttn-A (22.4%), and VentAttn-B (11.2%) networks. No significant association with receptor or transporter density maps were observed after applying FDR correction.

#### CN6 – Physiological Arousal (113 meta-analytic experiments)

The Physiological Arousal co-activation network mainly comprised meta-analytic experiments on Physiosexual Arousal (Poeppl, Langguth, Laird, & Eickhoff, 2014), Mechanical Pain (Xu et al., 2020), Non-Visceral Pain (Xu et al., 2020), Thermal Pain (Friebel, Eickhoff, & Lotze, 2011), Sexual Arousal (Penile Turgidity) (Kühn & Gallinat, 2011), Physiological Stress (Kogler et al., 2015), Noxious Cold Exposure (King & Carnahan, 2019), Experimentally-Induced Pain (Friebel et al., 2011), Self-Oriented Pain Empathy (Jauniaux et al., 2019), Affective Taste (Yeung, Goto, & Leung, 2018). This coactivation network was formed by activation in the insula (anterior to posterior), thalamus, periaqueductal grey, dorsal anterior mid cingulate cortex, lateral prefrontal cortex, amygdala, ventral putamen/pallidum, Crus 1 and Lobule VI.

Functional decoding using NeuroSynth indicated main association with terms underlying pain processing (i.e., pain, painful, noxious, nociceptive) sensory stimulation (i.e., electrical, stimulation, tactile, touch, intensity). Analyses using NeuroQuery rather yielded similarity with experiencing a sensation in general (i.e., experience, rating, feeling, sensation, salience, empathy). Voxels of the Physiological Arousal map showed important overlaps with those of the VentAttn (63.6%) followed by those from the SomMot (20.7 %) networks. More precisely, stronger overlaps were observed with the VentAttn-A (45.3%), VentAttn-B (22.9%), followed by the SomMot-B (20.1%). This network was mainly associated with VAChT (*r*=.48), H_3_ (*r*=.46), NET (*r*=.36), and α4β2 (*r*=.34) receptor/transporter density maps.

**Table 1.**
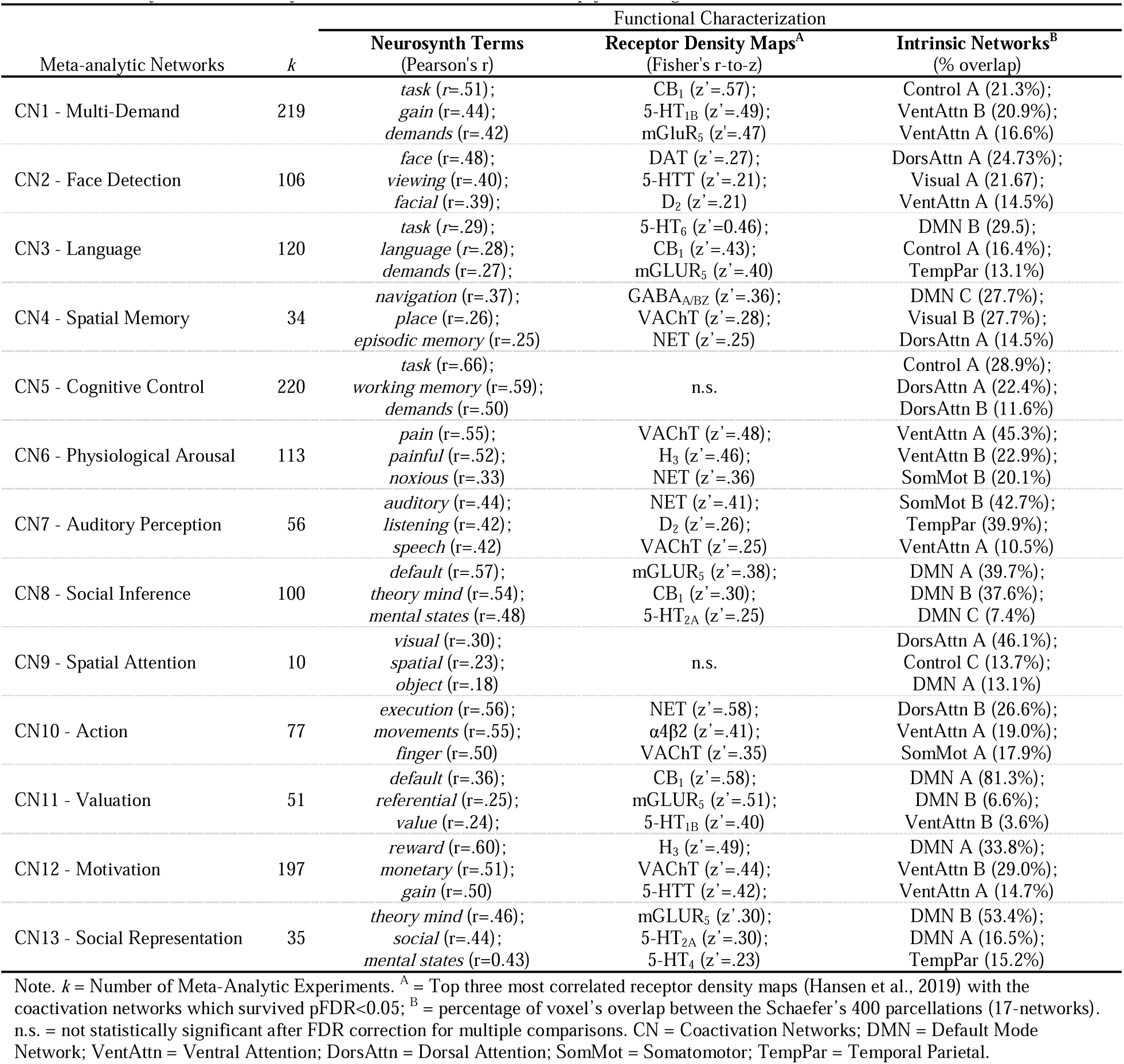
Summary of the Meta-analytic Coactivation Networks and their psychobiological correlates

#### CN7 – Auditory Perception (56 meta-analytic experiments)

The Auditory network mainly included meta-analyses on Audiovisual Stimuli (both Living and Static, (Csonka et al., 2021), Lexical Non-Tonal Tone (Murphy, Jogia, & Talcott, 2019), Tone (Samson, Zeffiro, Toussaint, & Belin, 2010), Speech (Turkeltaub & Coslett, 2010), Emotional Voices (Schirmer, 2018), Phoneme (Murphy et al., 2019), Music (Schirmer, 2018), Words Naming (McNorgan, Chabal, O’Young, Lukic, & Booth, 2015), Auditory-Laughter Humor (Farkas et al., 2021). This network was characterized by significant activity in the superior temporal gyrus/planum temporal (including auditory cortex A1-A2), dorsal aMCC, Crus 1 but also the precentral gyrus and right parietal operculum.

Functional decoding using NeuroSynth and NeuroQuery suggested both strong association with auditory terms (e.g., sounds, listened, acoustic, spoken, noise, frequency, signal, speaker). Voxels of the Auditory co-activation network revealed main overlap with voxels of the SomMot (55.1%) followed by those of the DMN (26.2%) and VentAttn (16.7%) intrinsic networks. More precisely, these overlaps were mainly due to the overlap with the SomMot-B (42.7%), TempPar (39.9%), followed by the VentAttn-A (10.5%) networks. This network spatially correlated with NAT (*r*=.41), D_2_ (*r*=.26), VAChT (*r*=.25), and DAT (*r*=.13) receptor/transporter density maps.

#### CN8 – Social Inference (100 meta-analytic experiments)

The Social Inference co-activation network was mainly formed meta-analyses Social Concepts (Arioli, Gianelli, & Canessa, 2021), Cognitive Morality (Eres, Louis, & Molenberghs, 2018), ToM Stories (Molenberghs, Johnson, Henry, & Mattingley, 2016), False Beliefs (van Veluw & Chance, 2014), False Beliefs (Diveica et al., 2021), False Beliefs (Krall et al., 2015), Trait Judgments (Theory of Mind) (Schurz, Radua, Aichhorn, Richlan, & Perner, 2014), Other’s morality (Eres et al., 2018), False Beliefs Reasoning (Schurz, Aichhorn, Martin, & Perner, 2013), and Cognitive Theory of Mind (Molenberghs et al., 2016). This co-activation network involved the activation of temporo-parietal junction, posterior cingulate cortex, medial prefrontal cortex, anterior and posterior middle temporal gyrus (but not mid subregion), lateral orbitofrontal cortex, amygdala, hippocampus, parahippocampal gyrus and pre-SMA.

Functional characterization using Neurosynth and NeuroQuery both revealed strong association with default mode network, theory of mind, mentalizing, self-referential processes (i.e., autobiographical, beliefs). Voxels of this co-activation network showed substantial overlap with those of the DMN (89.7%). This was mainly due to stronger overlap with DMN-A (39.7%) and DMN-B (37.6%), compared to DMN-C (7.3%) networks. Significant spatial correlations were observed with mGluR_5_ (*r*=.38), CB_1_ (*r*=.30), and 5-HT_2A_ (*r*=.25) receptor density maps.

#### CN9 – Spatial Attention (19 meta-analytic experiments)

Spatial Attention co-activation network included a wide range of meta-analytic experiments such as Spatial Navigation (Egocentric) (Derbie et al., 2021), Precue Spatial Attention (Wallis, Stokes, Cousijn, Woolrich, & Nobre, 2015), Task Deactivation (Schilbach, Eickhoff, Rotarska-Jagiela, Fink, & Vogeley, 2008), Familiarity Processing (Qin & Northoff, 2011), Words (McNorgan et al., 2015), Semantic Retrieval (Words) (Kim, 2016), Orthography (Wu, Ho, & Chen, 2012), Working Memory in Adolescence (Andre, Picchioni, Zhang, & Toulopoulou, 2016), Egocentric Spatial Coding (Derbie et al., 2021), and Focused Attention Mediation (Deactivation) (Fox et al., 2016). This network comprised the lateral occipital cortex, precuneus, Brodmann area 6, fusiform gyrus, angular gyrus, temporo-occipital fusiform gyrus, and the lateral prefrontal cortex.

Functional characterization revealed that the Spatial Attention CN was associated with visuospatial processes (i.e., space, spatial, visual, visuospatial) and attention (i.e., attention, attentional, spatial attention). Voxels of the spatial attention coactivation network were well distributed across those of the DorsAttn (32.6%), Visual (29.3%), FP (20.7%), and the DMN (17.4%) networks. Examining overlap with the 17 networks rather suggested stronger overlaps with the DorsAttn-B (46.1%), followed by the FP-C (13.7%) and the DMN-A (13.1%) networks. No significant association with receptor or transporter density maps were observed after applying FDR correction.

#### CN10 – Action (77 meta-analytic experiments)

The Action co-activation network comprised meta-analytic experiments such as Motor Task (Yang, 2015), Motor Imagery (Hardwick, Caspers, Eickhoff, & Swinnen, 2018), Action Execution (Papitto, Friederici, & Zaccarella, 2020), Reaching (Blangero et al., 2009), Motor Planning (Papitto et al., 2020), Motor Imagery (McNorgan, 2012), Motor Imagery (Hétu et al., 2013), Motor Learning (Hardwick, Rottschy, Miall, & Eickhoff, 2013), Motor Imagery (Papitto et al., 2020), Prosaccades (Cieslik, Seidler, Laird, Fox, & Eickhoff, 2016). This CN include activation of the SMA, thalamus, Lobule VI, premotor, precentral gyrus, putamen/pallidum, lateral occipital cortex and central opercular.

Functional characterization using Neurosynth revealed strong association with execution, movements, and imagery, whereas NeuroQuery rather include closer similarity with performance, simple, inhibition, button. Overlaps in voxels with the 7 networks suggested main overlaps with the DorsAttn (47.1%) and the SomMot (23.7%) networks. More precisely stronger overlaps were observed with the DorsAttn-B (26.6%), VentAttn-A (19.0%), SomMot-A (17.9%) and DorsAttn-A (14.4%) networks. Significant spatial correlations were observed with NET (*r*=.58), α4β2 (*r*=.41), and vAChT (*r*=.35) receptor/transporter density maps.

#### CN11 – Valuation (51 meta-analytic experiments)

The Valuation network was formed by meta-analyses on Food Value (Clithero & Rangel, 2014), Beautiful Faces (Chuan-Peng, Huang, Eickhoff, Peng, & Sui, 2020), State Anger (Puiu et al., 2020), Model-Based Decision-Making (Y. Huang, Yaple, & Yu, 2020), Social Norm Representation (Zinchenko & Arsalidou, 2018), Intention (Gilead, Liberman, & Maril, 2013), Positive Face Evaluation (Mende-Siedlecki et al., 2013), Subjective Value (Levy & Glimcher, 2012), Beautiful Visual Art (Chuan-Peng et al., 2020), and Upward Comparison (Yaple & Yu, 2020). Activated brain regions included the (ventro)medial prefrontal cortex/perigenual ACC, posterior MCC, ventral PCC, angular gyrus, dlPFC, Nucleus Accumbens and the left anterior insula/ventrolateral prefrontal cortex.

Functional Characterization suggested that this network was associated with self-referential processes (default, referential, autobiographical) but also in valuation (i.e., value, reward, money, valence). Substantial overlaps were observed between voxels of the valuation co-activation network and those of the DMN (92.0%). This overlap was mainly due to the DMN-A (81.3%), compared to the DMN-B (6.6%) and the DMN-C (2.8%). Finally, strong spatial association was observed in relation with CB_1_ (*r*=.58), mGluR_5_ (*r*=.51), 5-HT_1B_ (*r*=.40), H_3_ (*r*=.39), and 5-HT_2A_ (*r*=.38).

#### CN12 – Motivation (197 meta-analytic experiments)

the Motivation network was formed by meta-analytic experiments on Subjective Value Reward (Bartra, McGuire, & Kable, 2013), Risk Processing (Wu, Sun, Camilleri, Eickhoff, & Yu, 2021), Reward Outcome (Bartra et al., 2013), Reward (Wang, Zhang, & Jia, 2019), Anticipation of Aversive Stimuli (Andrzejewski, Greenberg, & Carlson, 2019), Subjective Value Negative (Bartra et al., 2013), Prediction Error Outcome (RPE) (Chase, Kumar, Eickhoff, & Dombrovski, 2015), Anticipation of Reward (Liu et al., 2011), Uncertainty during Associative Learning (Morriss, Gell, & van Reekum, 2019), Decision Risk (Wu et al., 2021). This CN was characterized by activity in the mesolimbic system as well as the ventromedial prefrontal cortex, posterior cingulate cortex, lateral occipital cortex, Brodmann area 44 and mid insula.

Functional characterization using Neurosynth and NeuroQuery both indicated strong associations with reward processing terms (i.e., reward, gain, monetary, monetary incentive, monetary reward) as well as motivation. Voxels of this network mainly overlapped with those of the DMN (44.6%) and the VentAttn (32.9%) networks. These spatial overlaps were principally due to the DMN-A (33.8%) and the VentAttn-B (28.9%). Finally, main spatial correlations with receptor/transport density maps included H_3_ (*r*=.49), vAChT (*r*=.44), 5-HTT (*r*=.42), DAT (*r*=.41), and MOR (*r*=.39).

#### CN13 – Social Representation (35 meta-analytic experiments)

The Social Representation network was formed by meta-analytic experiments underpinning theory of mind such as Theory of Mind (Diveica et al., 2021), Theory of Mind (Molenberghs et al., 2016), Theory of Mind (Bzdok et al., 2012), Theory of Mind (Schurz et al., 2014), Social Cognition (Adolfi et al., 2017), Social Cognition and Theory of Mind (Van’t Hooft et al., 2021), Mentalizing (Bellucci, Camilleri, Eickhoff, & Krueger, 2020), Semantic Autobiographical Memory (Martinelli, Sperduti, & Piolino, 2013), Narrative Comprehension (Mar, 2011). This co-activation network was characterized by activity in the lateral orbitofrontal cortex, middle temporal gyrus, inferior frontal gyrus, temporo-parietal junction, precuneus, dorsomedial prefrontal cortex, amygdala, pre-SMA, precentral, ventromedial prefrontal cortex, thalamus, fusiform gyrus and ventral caudate. In contrast to the CN8 – Social Inference, this co-activation additionally includes the activity of the mid-MTG as well as the thalamus.

Functional characterization using Neurosynth and NeuroQuery both revealed main association with theory of mind (i.e., mental states, mentalizing, social interaction) as well as semantic processing (comprehension, sentences, narrative, meaning). Overlaps between voxels were mainly observed in relation with the DMN (83.7%) followed by the FP (7.6%) networks. More precisely, overlaps were stronger for the DMN-B (53.4%), DMN-A (16.5%) and the TempPar (15.2%) networks. This network was significantly associated with mGluR_5_ (*r*=.30), 5-HT_2A_ (*r*=.30), 5-HT_4_ (*r*=.23), and D_2_ (*r*=.23) receptor/transporter density maps.

### 3.2. Voxelwise Similarity & Spatial Overlaps between Co-activation Networks

Given that our approach does not restrict voxels to one particular network, we sought to examine the locations of the strongest spatial overlaps between maps. The meta-analytic maps showed small-to-moderate correlation between network’ voxels (average correlation between networks, *r*=.216), confirming their spatially distinct features. Furthermore, we observed that strongest pairwise correlations were found between the CN1 -Multi-Demand Network & the CN5 -Cognitive Control Network (*r*=.48), CN8 - Social Inference and CN12 - Social Representation (*r*=.41), CN3.- Language and CN1 - Multi-Demand Networks (*r*=.41), CN10 - Action and CN5 - Cognitive Control (*r*=.399), and between the CN2 - Face Detection and CN12 - Motivation networks (*r*=.396). The weakest correlation was found between the Action Network and the Social Inference Network (*r*=-0.045).

To clarify these correlations, we sought to examine which voxels (and corresponding brain regions) showed the greatest overlaps in reported activations across networks. Main spatial overlaps between voxels of the meta-analytic co-activation networks were observed in the anterior midcingulate cortex (aMCC) (x=-2, y=14, z=48, overlap=9/13 networks), left (x=-36, y=24, z=-2, overlap=7/13 networks), right ventrolateral prefrontal cortex (x=34, y=26, z=0, overlap=7/13 networks), left thalamus (x=-12, y=-14, z=10, overlap=6/13 networks), left dorsolateral prefrontal cortex (dlPFC) (x=-48, y=20, z=22, overlap=6/13 networks), the left middle frontal gyrus (MFG) (x=-48, y=6, z=28, overlap=6/13 networks) as well as the left amygdala (x=-22, y=-4, z=-15, overlap=5/13 networks) (see Figure 4).

**Figure 4.**
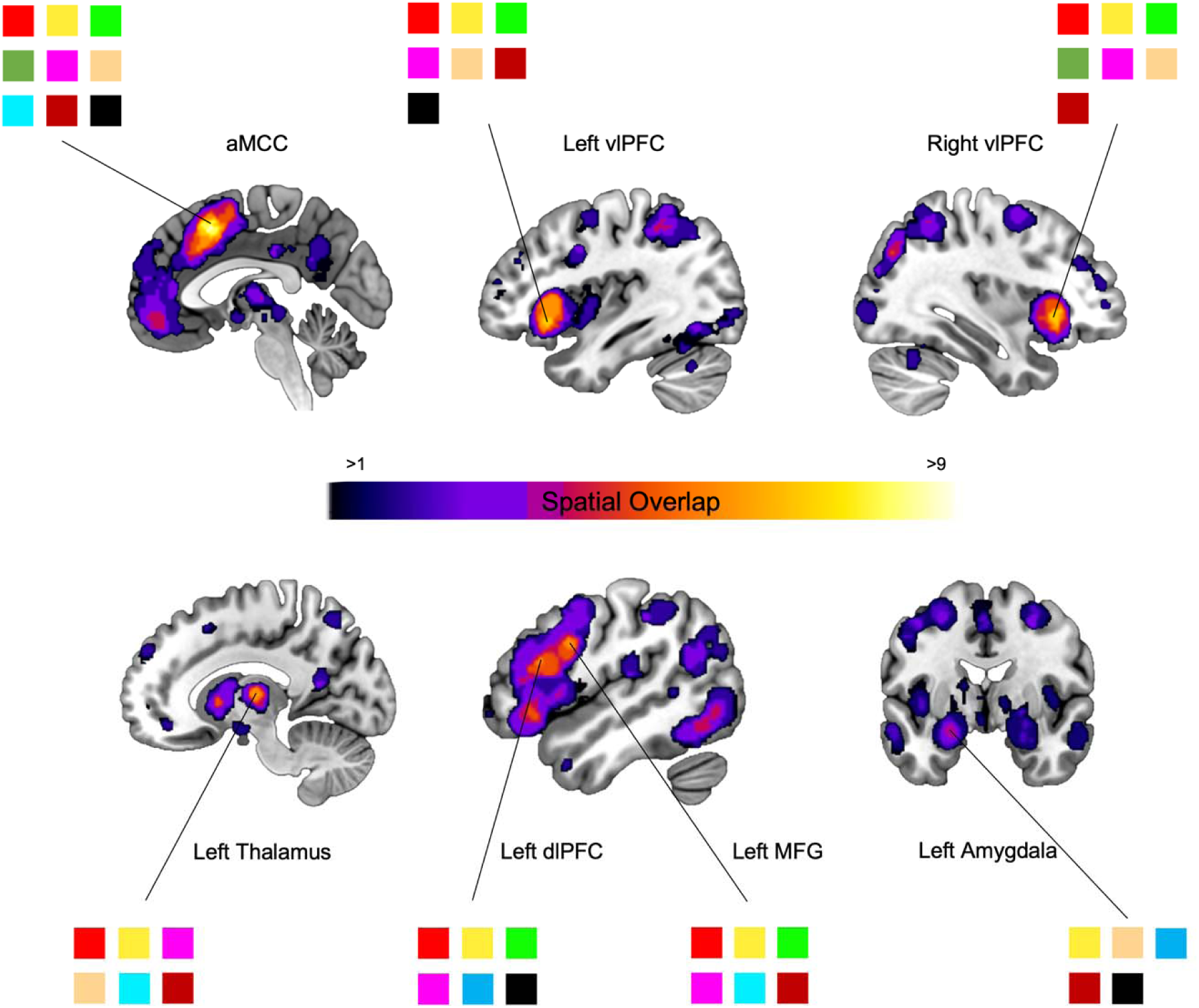
Main spatial overlap across voxels of the meta-analytic co-activation networks. Spatial overlaps between networks were mainly observed in the aMCC (9 networks), left & right vlPFC (both 7 networks), thalamus (6 networks), left dlPFC (6 networks), left MFG (6 networks), and left amygdala (5 networks). Red = Multi-Demand; Yellow = Face Detection; Green = Language; Dark Green = Spatial Memory; Magenta = Cognitive Control; Beige = Physiological Arousal; Purple = Auditory Perception; Sky Blue = Social Inference; Blue = Spatial Attention; Cyan = Action; Orange = Valuation; Maroon = Motivation; Black = Social Representation. aMCC = anterior MidCingulate Cortex; vlPFC = ventrolateral Prefrontal Cortex; dlPFC = dorsolateral Prefrontal Cortex; MFG = Middle Frontal Gyrus

### 3.3. Comparison with previous Data-Driven Meta-analytic Networks

The co-activation networks were then compared to Bolt and colleagues (2020)’ maps (Figure 5). We observed that their Passive Object Viewing map (e.g., Naming: Overt; Passive Viewing; Perception: Vision Shape) was mainly associated with the CN5 – Cognitive control (z_r_=.89), followed by the CN2 – Face Detection (z_r_=.82). Both maps (Passive Object Viewing and Cognitive Control) showed strong activation in bilateral fusiform gyrus. Their Motor Action map (e.g., Finger Tapping, Clicks, Electromyography) spatially matched with the CN10 – Action (z_r_=1.01). Common neurobiological features across both maps include the SMA and the MCC. Their Active Object Viewing map (e.g., Mental Rotation, Spatial Cognition, Saccades) showed greatest similarity with the CN5 – Cognitive Control (z_r_=1.26). Also, their Social/Self-Memory map (e.g., Theory of Mind, De-activation, Social Cognition) was spatially associated with both CN11 – Valuation (z_r_=.628) and CN8 – Social Inference (z_r_=.627). The Social/Self-Memory and Valuation maps both reported greater activation in the mPFC. We also observed that their Auditory Perception map (e.g., Passive Listening, Audition Perception, Pitch Monitor/Discrimination) was more strongly related to the CN1 – Multi-Demand Network (z_r_=.75) followed by the CN7 – Auditory Perception (z_r_=.65). This may be partially explained by strong activation in lateral frontal regions such as the IFG, in both maps. Their Affect/Reward map (e.g., Emotion: Fear; Memory; Affective Pictures) was spatially correlated with the CN2 – Face Detection (z_r_=.89) as well as the CN12 – Motivation (z_r_=.82) networks. Both Affect/Reward and CN2 networks showed strong activation in bilateral amygdala. Finally, their Higher-Order Cognition map (i.e., Word Generation [Overt/Covert], Stroop Color Word) spatially corresponded to the CN1 – Multi Demand Network (z_r_=1.11) with greatest activation in bilateral anterior insula.

**Figure 5.**
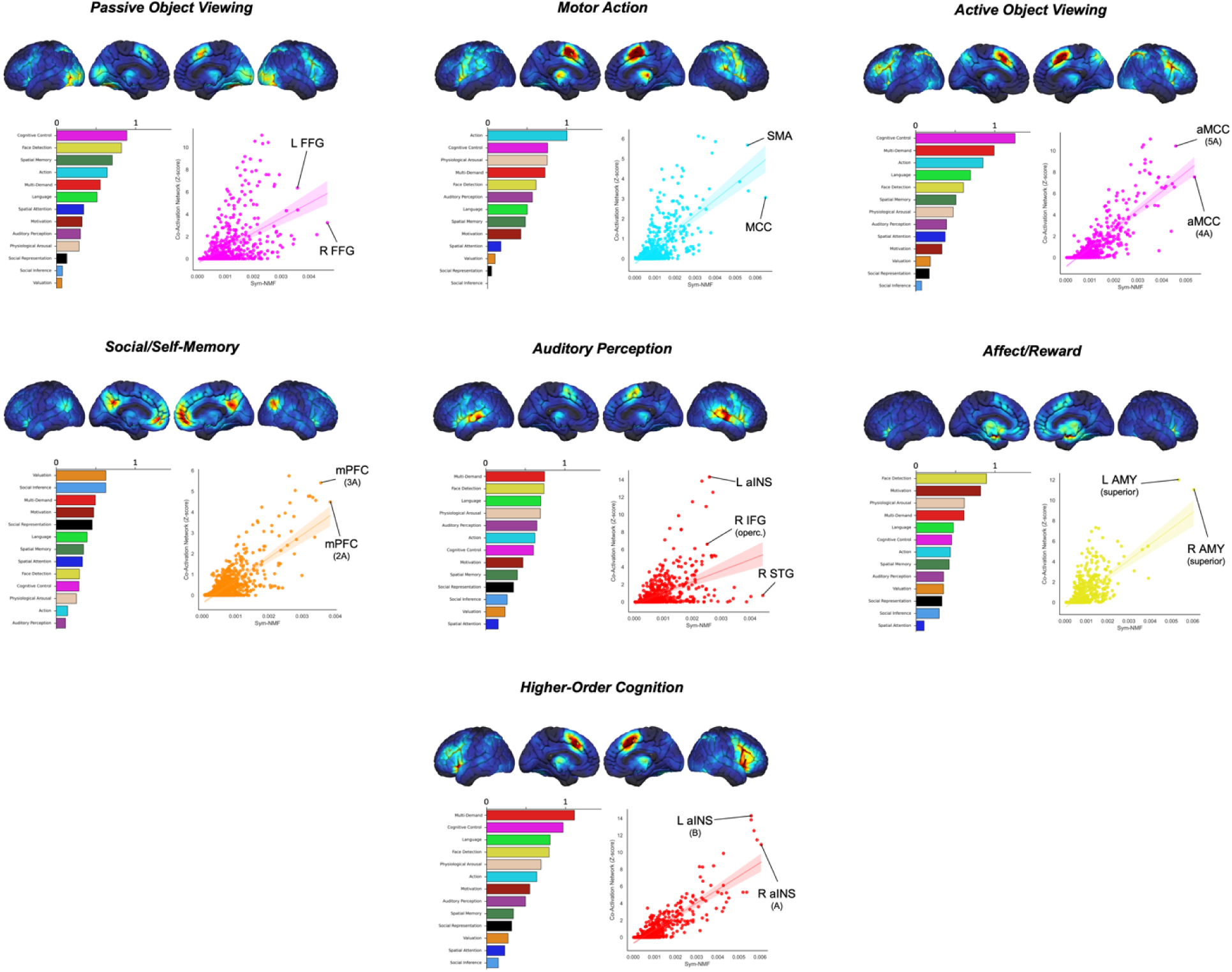
Comparison with previous Data-driven Meta-analytic Networks from Bolt and colleagues (2020). Bar graphs represent relationship between Bolt et al. (2020)’s networks identified through Symmetric non-negative matrix factorization and those identified in the current study. Spearman’s correlation coefficients were transformed using Fisher z-transformation. Scatter plots represent the original association between Bolt et al. (2020)’s network and the top corresponding Meta-analytic Network at a ROI-level (728 regions, see Method Section). L = Left; R = Right; FFG = Fusiform Gyrus; SMA = Supplementary Motor Area; MCC = Midcingulate Cortex; aMCC = anterior Midcingulate Cortex; mPFC = medial Prefrontal Cortex; aINS = anterior Insula; IFG = Inferior Frontal Gyrus; STG = Superior Temporal Gyrus; AMY = Amygdala; L = Left; R = Right

## 4. DISCUSSION

To address several limitations surrounding expert-driven categorization of mental functions, we aimed to identify the core data-driven co-activation networks across task-based fMRI meta-analyses (i.e., 1347 meta-analytic experiments, including 19,822 meta-analytic peaks) conducted in the past 20 years. Using a hierarchical clustering framework, based on co-activation similarity, we found that the neurobiological processes can be reduced to 13 homogeneous co-activation networks, namely Multiple-Demand, Face Detection, Language, Spatial Memory, Cognitive Control, Physiological Arousal, Auditory Perception, Social Inference, Spatial Attention, Action, Valuation, Motivation, and Social Representation networks. These co-activation networks were characterized by distinct associations with 19 PET receptor/transporter density maps (Hansen et al., 2022) as well as intrinsic functional connectivity networks (Schaefer et al., 2018). More importantly, our results were fairly consistent with other data-driven approaches using the BrainMap database (Bolt et al., 2020), or with a corpus of 20,000 fMRI studies (Beam et al., 2021). These results provide new insights about the main co-activation patterns across the task-based fMRI literature.

Over the past few years, several meta-analyses showed that two psychologically distinct constructs may rely on similar (or identical) neural maps (Chen et al., 2022; Lindquist et al., 2016). Using a data-driven method on a large number of published meta-analyses, we sought to identify common brain circuits across fMRI tasks. One main limitation of psychologically driven ontologies of mental faculties is that two psychological constructs may rely on activation of common brain regions. For example, the RDoC classification system clearly delineate between positive (e.g., reward anticipation/outcome, valuation, habit) and negative valence systems (e.g., anxiety, fear, loss, frustrative non-reward) (Kozak et Cuthbert, 2016). In our study, we did not observe a specific co-activation network underpinning *emotion* (i.e., discrete emotional categories). This can be attributable to the fact that some meta-analyses on emotions were classified in Face Perception co-activation networks due to methodological reasons. Indeed, across neuroimaging meta-analyses on emotions (in general or in discrete categories), many authors included “*viewing emotional facial stimuli*” which may have diluted the *true* effect of emotional response (Kober et al., 2008; Lindquist et al., 2016; Lindquist, Wager, Kober, Bliss-Moreau, & Barrett, 2012; Tao et al., 2021; Vytal & Hamann, 2010). Another possible explanation may be that previous meta-analyses on emotion (Lindquist et al., 2016), reward & punishment anticipation (Chen et al., 2022) as well as positive reward prediction error (Corlett, Mollick, & Kober, 2022) show similar activation in brain circuits involving the vmPFC, aMCC, aINS, dlPFC, basal ganglia, the amygdala as well as the midbrain. This could explain why the Affect/Reward map found by Bolt and colleagues (2020) mostly correlated with both our CN2 – Face Detection & our CN12 – Motivation Network. However, contrasting with our results, Beam et al. (2021)’s data-driven framework did find that Emotion was significantly distinguished from other processes based on activation of the amygdala. It is noteworthy to mention that, contrarily to ours and Bolt et al’s (2020), their framework assumed that mental functions can be identifiable through non-overlapping sets of brain structures; that, each structure is forced to belong to only one mental function. Intriguingly, it is relatively well accepted that the amygdala is significantly activated across various mental functions that are not specific to *emotion*, such as detection of visual stimuli in general as well as empathy and theory of mind (Adolphs, 2008, 2010; Pessoa & Adolphs, 2010). Likewise, we found that the amygdala was significantly activated in 5 co-activation networks including Face Detection, Physiological Arousal, Motivation, Social Inference and Social Representation (see Figure 4). Therefore, we argue that the co-activation pattern during which the activity of the amygdala occurs is much more informative than the structure alone.

The RDoC Social Processes include four distinct yet related subconstructs, such as affiliation and attachment (e.g., positive social interaction/social bonding), social communication (e.g., facial expression), self-perception and understanding (e.g., agency) and other-perception and understanding (e.g., understanding other’s mental states) (Kozak et Cuthbert, 2016). Recently, Pintos Lobo and colleagues (Pintos Lobo et al., 2023) conducted a meta-analysis of fMRI studies using the Social Cognition systems as described by the RDoC. They showed that the social cognition subsystems spatially overlapped across most brain regions (e.g., vmPFC, dmPFC, aMCC, FFG, IPL, pSTS, aINS), albeit these maps also showed some specific effects. More interestingly, the authors found poor spatial correspondence between the RDoC-driven and their own data-driven co-activation networks, therefore challenging the expert-driven categorization of social cognition at a BOLD fMRI resolution. In our study, we showed that social cognitions could be neurobiologically distinguished by Face Detection, Social Inference and Social Representation networks. In contrast with results from Pintos Lobo and colleagues (2023), our findings fit relatively well the RDoC subconstructs of Social Processes. More importantly, we replicated the meta-analytic findings of Schurz et al. (Schurz et al., 2021) demonstrating the heterogeneity in brain circuits underpinning theory of mind. Indeed, they found 3 distinct neurobiological processes: Cognitive, Intermediate and Affective, which correspond to our CN8 (Social Inference), CN13 (Social Representation) and CN2 (Face detection) networks, respectively (Schurz et al., 2021). The most notable differences in brain activation between these three circuits are observed in the fronto-insular cortex (vlPFC and aINS) as well as in the mMTG. For example, only the intermediate (or Social Representation) significantly activated the mMTG, a region which was also found in our CN3 - Language and CN7 - Auditory perception (see also Xu et al., 2019). Surprisingly, Beam et al. (2021)’s study did not identify any data-driven domain characterized by social cognition, but rather found that face perception was blurred within a vision domain. Contrasting with this, Bolt and colleagues (2020) found a broad social/self-memory network (e.g., theory of mind, deactivation, social cognition, autobiographical recall). This map correlated significantly with our CN11 – Valuation, and CN8 – Social Inference, due to strong spatial overlaps in regions of the DMN (i.e., mPFC, vPCC, TPJ), which corroborates the role of this network in self-generated thoughts and social processes (Andrews-Hanna, Smallwood, & Spreng, 2014; Li, Mai, & Liu, 2014). As Pintos Lobo and colleagues (2023) suggested, much work is needed to evaluate how well the RDoC-based categorization of social processes fits with current fMRI task results and conversely, what phenotypes adequately distinguish between data-driven maps of social processes.

In our study, we found that the co-activation networks could be reduced to 13 homogeneous brain co-activation circuits. Despite the fact that we used largely different methods, we found similar findings to the recent Beam and colleagues (2022)’s study. Indeed, we observed a vision (i.e., face detection), memory (i.e., spatial memory), reward (i.e., motivation and valuation), cognition (i.e., cognitive control), manipulation (i.e., action), language (i.e., language) and hearing (i.e., auditory perception) networks. Moreover, we were able to characterize the neurochemical substrates of these task-based networks. Among the most robust correlations, we found that the mGluR5 was associated with the Multi-demand and Language networks, which is consistent with the vast preclinical and clinical data showing that glutamate plays a key role in cognition (Esterlis et al., 2022; Mecca, 2019; Régio Brambilla et al., 2020). Likewise, the strong correlations between the vAChT, the cannabinoid CB_1_ receptor, the DAT and the opioid-mu receptor and the Valuation and Motivation networks are consistent with the well-known roles of these receptors and transporters in the brain reward system (Parsons & Hurd, 2015; Skirzewski et al., 2022; Le Merrer, Becker, Befort, & Kieffer, 2009; Nummenmaa et al., 2018). That being said, distinguishing mental processes (as neuropsychological constructs) on a neurobiological basis remains one key challenge in neuroimaging. If two distinct mental processes show similar brain co-activation, how do we interpret their distinction at a psychological level. While this is yet to be elucidated, mental processes are inherently complex as it relies on numerous additional information to adequately label to a mental function such as the context (e.g., valence) in which it is observed. For example, a variety of neuropsychological tasks have been developed to assess executive functions such as planning, cognitive flexibility, verbal fluency, inhibitory control, working memory, processing speed (de Assis Faria, Veiga Dias Alves, & Charchat-Fichman, H., 2015). Although the latent structure that characterize the vast number of neuropsychological tests remains elusive, research nonetheless support that executive functions are best described by unity and diversity (Karr et al., 2018; Miyake et al., 2000). In our study, we found 2 main brain circuits underpinning executive functions, which were characterized by highly similar (e.g., aMCC/preSMA Caudate, Thalamus, FEF, IPL) but also distinct (e.g., Angular Gyrus, ventrolateral prefrontal cortex) brain structures (i.e., CN1 – Multi-Demand & CN5 – Cognitive Control). At a task-based fMRI (tbfMRI) BOLD resolution, executive functions appear to be hardly distinguishable given their substantial spatial overlaps. Such difficulty was also observed in prior data-driven studies (Beam et al., 2021; Bolt et al., 2020). One explanation is that increasing the sample size in our study would result in more homogeneous maps and therefore unveil such distinctions. It is also possible that tbfMRI BOLD activation may not be at an adequate resolution to identify significant spatial distinction between similar psychological labels. However, the functional connectivity patterns during tbfMRI may be a promising avenue to adequately distinguish between spatially similar behaviors. While there is a growing interest for brain-behavior prediction using functional connectivity (FC) patterns during resting-state fMRI (rsfMRI), recent findings indicate that FC patterns during tbfMRI outperforms the former modality when predicting interindividual differences in performance during various cognitive tasks (Greene, Gao, Scheinost, & Constable, 2018; Rosenberg et al., 2016; Zhao et al., 2023). Although the well-defined intrinsic connectivity networks may capture some behaviorally relevant patterns at rest, it remains unknown to what extent they reflect neural processes during an explicit task. In our study, we found that the intrinsic connectivity networks only partially overlapped with task coactivation networks. For example, voxels of the DMN, sometimes inadequately referred to as a task-negative network (see discussion in Sonuga-Barke & Castellanos, 2007 and Spreng, 2012), significantly overlapped with our CN8 – Social Inference, CN11 – Valuation, and CN13 – Social Representation networks. Besides this, the other networks (when examined using the 7- or 17-network parcellations) overlapped poorly with our task co-activation networks. More precisely, while some research suggests that the DMN may be related to episodic memory, this intrinsic network only partially overlaps with the brain activation seen during episodic memory tasks. This is mainly due to additional activation of non-DMN regions such as those from the Visual B (e.g., Retrosplenial cortex and Lingual Gyrus) and DorsAttn (e.g., Frontal Eye Fields, Intraparietal sulcus, Superior Parietal Lobule, MT+) (see CN4 – Spatial Memory). These spatial differences highlight some of the advantages of tbfMRI as well as the difficulty of rsfMRI to identify task-specific patterns. Nevertheless, it is also noteworthy to mention that differences in functional connectivity patterns between regions-of-interest (ROIs) of a given co-activation network may give us new insights about the complexity of distinct yet similar psychological terms, and aid the brain-behavior prediction, as shown by prior work on working memory capacity (Pläschke et al., 2020); emotion categories (Huang et al., 2018), personality traits (Nostro et al., 2018) and psychopathologies (e.g., depression, Schilbach et al., 2014). We thus encourage researchers to test whether FC patterns between meta-analytically derived ROIs may outperform parcels from canonical networks or brain structures in brain-behavior prediction.

## LIMITATIONS

First, the number of mental functions, as studied by task-based fMRI, is somewhat limited given the finite number of fMRI tasks. Therefore, our study (and data-driven framework in general) is biased by current literature. As Beam and colleagues (2022, p.10-11) noted: “*Researchers may be more inclined to seek and report evidence for mental functions which have been previously well described in psychology, a form of confirmation bias, and to treat these mental functions as stable constructs, a form of reification bias”.* Hence, we cannot rule out the possibility that other co-activation networks may arise when considering all mental functions. Developing novel fMRI tasks may thus allow us to identify other neurobiological processes which are crucial to characterize the whole spectrum of mental functions. Second, coordinate-based meta-analysis usually contains main activated peaks across studies, leaving aside the role of *deactivated* voxels. Prior work indicated that deactivation data may be highly informative when describing neurobiological processes (Shulman et al., 1997). However, the deactivation pattern is mainly task-dependent (Spreng, 2012). For example, when one does not receive an expected reward (frustrative non-reward), the nucleus accumbens and vmPFC seems to deactivate (Dugré & Potvin, 2021), which mirrors a diminished release in dopamine during reward omission (Schultz, 2016; Sosa, Mata-Luévanos, & Buenrostro-Jáuregui, 2021; Tian & Uchida, 2015). Further investigation is needed to determine whether two similar brain maps (based on activation), such as the multiple-demand and cognitive control networks, could be better distinguished by their pattern of deactivation. Studies using real-world data is necessary to validate our results and to test this more precisely. Lastly, the associations between the co-activation networks and PET receptor density maps (i.e., group-level) were assessed via spatial similarity. These should nonetheless be interpreted with cautious as specific studies are necessary to test whether interindividual variability in neurotransmitters density are indeed related to differences in tbfMRI activation.

## 5. CONCLUSIONS

Describing the brain circuits that underpin mental faculties has been a longstanding pursuit in neuroscience. A growing body of task-based activation research highlights that many processes may rely on common brain circuits, which raises questions about the neurobiological validity of expert-driven categories. Accordingly, we have shown that the results of neuroimaging meta-analyses from the last 20 years can be organized into 13 distinct brain circuits. These co-activation patterns showed different association with neurochemical substrates as well as intrinsic connectivity networks, stressing their distinctions. More importantly, we validated some co-activation patterns found in previous study, namely the Action, the Cognitive Control and the Multi-Demand Networks. This work, in conjunction with other data-driven framework (Bolt et al., 2020; Beam et al., 2021; Nakai et Nishimoto, 2020), should serve as a starting point to develop a neurobiologically-driven ontology to successfully map mental functions and their related psychopathologies. We encourage researcher to use these meta-analytic co-activation patterns as priors to aid the reliability in task-based fMRI, that is by using peak coordinates of meta-analytic in functional connectivity analyses and/or by investigating the relationships between deviation from *normative activation* and psychopathologies.

