## Supplementary Material for "Towards a Neurobiologically-driven Ontology of Mental Functions: A Data-driven Summary of the Twenty Years of Neuroimaging Meta-Analyses"

**Corresponding authors**

| **Supplementary Table 1.** Included Meta-analytic Experiments | | | |
| --- | --- | --- | --- |
| **First Author, Date** | **PMID** | **Contrast** | **Number of foci** |
| Adank et al., 2012 | 22633697 | Core Comprehension Network | 18 |
|  |  | Difficult Speech | 8 |
|  |  | Speech Production | 25 |
| Addis et al., 2016 | 27235570 | Autobiographical Memory Retrieval | 17 |
| Adolfi et al., 2017 | 28088652 | Emotion | 49 |
|  |  | Interoception | 30 |
|  |  | Social Cognition | 43 |
| Alain et al., 2018 | 29536592 | Listening Linguistic Complexity | 5 |
|  |  | Listening Spectrally Degraded Speech | 3 |
|  |  | Listening Speech Noise | 3 |
| Albrecht et al., 2010 | 19913573 | Intranasal Trigeminal Stimulation (CO2) | 42 |
| Amanzio et al., 2013 | 22125184 | Noxious Administration | 15 |
|  |  | Placebo Analgesia | 14 |
|  |  | Prestimulation Pain Expectation | 5 |
| Andre et al., 2016 | 27995059 | Working Memory (Adolescence) | 5 |
| Andrzejewski et al., 2019 | 31502205 | Anticipation of Aversive Stimuli | 13 |
| Araujo et al., 2013 | 24027520 | Judging Others Traits | 8 |
|  |  | Judging Others Traits (Close) | 10 |
|  |  | Judging Others Traits (Distant) | 19 |
|  |  | Judging Self-Traits | 9 |
| Arioli et al., 2020 | 32897484 | Emotional Concepts | 7 |
|  |  | Social Concepts | 13 |
| Arioli et al., 2019 | 31077492 | Processing Others Actions | 20 |
|  |  | Processing Others Mental States | 7 |
|  |  | Processing Social Interactions | 12 |
| Arsalidou et al., 2011 | 20946958 | Addition | 33 |
|  |  | Calculation Tasks | 30 |
|  |  | Multiplication | 31 |
|  |  | Number Tasks | 35 |
|  |  | Subtraction | 20 |
| Arsalidou et al., 2011 | 21350872 | Dynamic Faces | 20 |
| Arsalidou et al., 2017 | 28844728 | Calculations (Children) | 10 |
|  |  | Number Tasks (Children) | 5 |
| Arsalidou et al., 2020 | 31927758 | Erotic Processing | 24 |
|  |  | Food Processing | 21 |
|  |  | Money Processing | 19 |
| Arya et al., 2016 | 27367364 | Bladder Filling | 7 |
| Bartley et al., 2018 | 29944961 | Mathematical Problem Solving | 9 |
|  |  | Verbal Problem Solving | 14 |
|  |  | Visuospatial Problem Solving | 13 |
| Bartra et al., 2013 | 23507394 | Decision Reward | 3 |
|  |  | Outcome Stage | 6 |
|  |  | Punishment Outcome | 5 |
|  |  | Reward Outcome | 8 |
|  |  | Subjective Valuation (Money) | 4 |
|  |  | Subjective Valuation (Primary Incentives) | 4 |
|  |  | Subjective Value Negative | 7 |
|  |  | Subjective Value Reward | 9 |
| Basten et al., 2015 | 10.1016j.intell.2015.04.009 | Intelligence | 8 |
| Beissner et al., 2013 | 23785162 | Parasympathetic Regulation | 9 |
|  |  | Sympathetic Regulation | 10 |
| Bello et al., 2020 | 32848663 | Mirror Visual Illusion | 12 |
| Bellucci et al., 2017 | 27859899 | Trust Conditional | 2 |
|  |  | Trust Decision Reciprocity | 4 |
|  |  | Trust Feedback of Decision | 3 |
| Bellucci et al., 2018 | 29958872 | Reciprocity Decisions | 4 |
|  |  | Rejection Unfair Offers | 5 |
|  |  | Trust Decisions | 2 |
| Bellucci et al., 2020 | 32707344 | Empathy | 11 |
|  |  | Mentalizing | 9 |
|  |  | Prosocial Behaviors | 4 |
| Bellucci et al., 2020 | 32302599 | Social Punishment | 3 |
|  |  | Social Punishment (Second-Party) | 2 |
|  |  | Social Punishment (Third-Party) | 2 |
| Belyk et al., 2013 | 23934416 | Affective Prosody | 30 |
|  |  | Linguistic Prosody | 22 |
| Benoit et al., 2015 | 26142352 | Episodic Stimulation & Memory | 32 |
| Berretz et al., 2021 | 33497786 | Stress Induction | 3 |
|  |  | Stress Induction | 5 |
| Biggs et al., 2020 | 33011229 | Pain Fear (CS+) | 9 |
|  |  | Pain Fear (CS-) | 14 |
| Bisenius et al., 2015 | 26241685 | Bistable Perception | 7 |
|  |  | Consciousness | 11 |
|  |  | Masking Paradigm | 9 |
| Blangero et al., 2009 | 19109986 | Reaching | 17 |
| Boccia et al., 2015 | 26322002 | Creativity | 28 |
|  |  | Musical Creativity | 11 |
|  |  | Verbal Creativity | 20 |
|  |  | Visuospatial Creativity | 6 |
| Boccia et al., 2015 | 26619805 | Visual Aesthetic Experience | 27 |
| Boccia et al., 2016 | 26809288 | Moral Reasoning | 45 |
|  |  | Moral Reasoning (1PP) | 34 |
|  |  | Moral Reasoning (3PP) | 36 |
| Boccia et al., 2019 | 31491472 | Episodic Autobiographical Memory | 19 |
|  |  | Recent Autobiographical Memories | 8 |
|  |  | Recent Autobiographical Memories | 2 |
| Bohrn et al., 2012 | 22824234 | Figurative Language | 6 |
|  |  | Literal Language | 3 |
| Bottenhorn et al., 2019 | 30793072 | Naturalistic | 51 |
| Bourguignon et al., 2014 | 25305636 | Discourse Generation | 4 |
|  |  | Picture Naming | 5 |
|  |  | Sentence Generation | 6 |
|  |  | Sentence Reading | 6 |
|  |  | Word Generation | 2 |
|  |  | Word Reading | 1 |
| Bourguignon et al., 2019 | 30831311 | Lexico-Semantic | 5 |
|  |  | Working Memory | 5 |
| Brooks et al., 2012 | 22001789 | Subliminal Arousal | 14 |
|  |  | Subliminal Emotional Faces | 7 |
|  |  | Subliminal Facial Arousal | 10 |
|  |  | Subliminal Physiological Arousal | 9 |
| Brooks et al., 2017 | 27539864 | Emotional Words | 14 |
|  |  | Emotions | 22 |
| Brown et al., 2005 | 15846815 | Fluent Production | 17 |
| Brown et al., 2009 | 19162389 | Syllable Singing | 24 |
| Brown et al., 2011 | 21699987 | Audition (Aesthetic) | 8 |
|  |  | Gustation (Aesthetic) | 14 |
|  |  | Olfaction (Aesthetic) | 6 |
|  |  | Vision (Aesthetic) | 33 |
| Bryant et al., 2016 | 10.190702572-7389-160005 | Moral Dilemmas | 14 |
|  |  | Moral Judgments of Right or Wrong | 16 |
| Buchsbaum et al., 2005 | 15846821 | Go No-Go | 14 |
|  |  | Task Switching | 15 |
|  |  | Wisconsin Card | 15 |
| Buhle et al., 2014 | 23765157 | Emotion Reappraisal | 32 |
|  |  | Emotional Baseline | 3 |
| Bzdok et al., 2011 | 20978908 | Facial Positive Assessment | 4 |
| Bzdok et al., 2012 | 22270812 | Empathy | 22 |
|  |  | Morality | 12 |
|  |  | Theory of Mind | 17 |
| Cacioppo et al., 2013 | 24002359 | Rejection (Romantic Breakup) | 8 |
|  |  | Rejection (Stranger Cyberball) | 8 |
| Cai et al., 2014 | 25355218 | Go No-Go & Stop-Signal | 60 |
| Cai et al., 2019 | 31641118 | Inhibition (Adult) | 48 |
|  |  | Inhibition (Children) | 47 |
| Calder et al., 2011 | 21536554 | Anxiety when viewing Negative Faces | 8 |
| Carter et al., 2010 | 10.1037a0018046 | Delay Discounting | 24 |
| Caspers et al., 2010 | 20056149 | Action Imitation | 14 |
|  |  | Action Observation | 19 |
| Cespón et al., 2020 | 33025513 | Simon Task | 7 |
| Chae et al., 2013 | 23395475 | Acupuncture Stimulation (Activation) | 16 |
|  |  | Acupuncture Stimulation (Deactivation) | 12 |
|  |  | Tactile Stimulation (Activation) | 8 |
|  |  | Tactile Stimulation (Deactivation) | 3 |
| Chase et al., 2015 | 25665667 | Fixed Learning Rate (RPE) | 3 |
|  |  | Individual Learning Rate (RPE) | 13 |
|  |  | Instrumental (RPE) | 10 |
|  |  | Liquid Stimuli (RPE) | 5 |
|  |  | Money Stimuli (RPE) | 7 |
|  |  | Pavlovian (RPE) | 5 |
|  |  | Prediction Error Outcome (RPE) | 9 |
|  |  | Reward Prediction Error | 15 |
|  |  | Social Stimuli (RPE) | 3 |
|  |  | Temporal Difference Models (RPE) | 4 |
| Chauvigne et al., 2014 | 25324765 | Audiomotor Externally-Paced | 25 |
|  |  | Audiomotor Memory-Paced | 12 |
|  |  | Audiomotor Self-Paced | 26 |
| Chein et Schneider, 2005 | 16242923 | Practice (Decreases) | 14 |
|  |  | Practice (Increases) | 4 |
| Chen et al., 2017 | 28480333 | Animal Concepts | 6 |
|  |  | Artifact Concepts | 6 |
| Chen et al., 2018 | 30083997 | Cognitive Interference | 8 |
|  |  | Emotional Interference | 6 |
| Chen et al., 2020 | 32555497 | Body Mass Index (Sweet Stimuli) | 11 |
|  |  | Fasting (Sweet Stimuli) | 13 |
|  |  | Hunger (Sweet Stimuli) | 13 |
|  |  | Sweet Stimuli | 35 |
| Chen et al., 2020 | 32472741 | Drawing Creativity | 8 |
|  |  | Literary Creativity | 8 |
|  |  | Music Creativity | 4 |
| Chouinard et al., 2010 | 19800353 | Naming Animals | 17 |
|  |  | Naming Tools | 21 |
| ChuanPeng et al., 2020 | 33089442 | Beautiful Faces | 7 |
|  |  | Beautiful Visual Art | 3 |
| Cieslik et al., 2015 | 25446951 | Action Cancellation (Stop Signal) | 19 |
|  |  | Cognitive Control | 30 |
|  |  | Spatial Interference | 17 |
|  |  | Stroop Task | 25 |
|  |  | Withhold Prepotent Response | 14 |
| Cieslik et al., 2016 | 27211526 | Antisaccades | 12 |
|  |  | Prosaccades | 11 |
| Clark et al., 2020 | 32189178 | Proactive Control (Withholding) | 11 |
|  |  | Reactive Control (Inhibition) | 8 |
| Clithero et al., 2014 | 23887811 | Decision Values (CBMA) | 7 |
|  |  | Decision Values (IBMA) | 4 |
|  |  | Food | 3 |
|  |  | Money | 8 |
|  |  | Other Reward | 7 |
|  |  | Outcome Values | 6 |
|  |  | Subjective Value | 9 |
| Cocquyt et al., 2019 | 10.1016j.jneuroling.2019.04.001 | Compositional Association | 2 |
|  |  | Semantic Categorization | 2 |
| Cogdell-Brooke et al., 2020 | 32845058 | Divergent Thinking | 3 |
| Cona et al., 2015 | 25704073 | Encoding Intention | 10 |
|  |  | Maintenance of Intention | 21 |
|  |  | Maintenance of Intentions (Activations) | 20 |
|  |  | Maintenance of Intentions (Deactivations) | 16 |
|  |  | Retrieval of Intention | 48 |
| Cona et al., 2016 | 27185531 | Maintenance (F Prospective Memory) | 11 |
|  |  | Maintenance (NF Prospective Memory) | 13 |
|  |  | Retrieval (F Prospective Memory) | 20 |
|  |  | Retrieval (NF Prospective Memory) | 30 |
| Cona et al., 2019 | 30600568 | Long Term Navigation | 9 |
|  |  | Mental Rotation | 22 |
|  |  | Spatial Processing | 26 |
|  |  | Spatial Working Memory | 18 |
| Crepaldi et al., 2013 | 23825451 | Lexicon | 32 |
| Criaud et al., 2013 | 23164813 | Complex Stimulus Identification (No-Go) | 5 |
|  |  | No-Go Trials | 11 |
|  |  | Working Memory Loads (No-Go) | 5 |
| Cromheeke et Mueller, 2014 | 23563751 | Emotion-Cognition Interaction | 18 |
| Csonka et al., 2021 | 33718874 | Congruent Audiovisual Paradigms | 4 |
|  |  | Dynamic Audiovisual Stimuli | 4 |
|  |  | Incongruent Audiovisual Paradigms | 2 |
|  |  | Living Audiovisual Stimuli | 4 |
|  |  | NonVocal Audiovisual Stimuli | 1 |
|  |  | Nonliving Audiovisual Stimuli | 1 |
|  |  | Static Audiovisual Stimuli | 3 |
|  |  | Vocal Audiovisual Stimuli | 4 |
| Cutler et al., 2018 | 30195947 | Altruistic Decision to Give | 13 |
|  |  | Selfish Decision | 27 |
|  |  | Strategic Decision to Give | 20 |
| Dahlgren et al., 2020 | 32416099 | Emotional Encoding | 58 |
|  |  | Retrieval Emotional | 29 |
| DalBò et al., 2020 | 33179726 | Emotional Body Odor | 4 |
| Daniel et al., 2016 et al., 2 | 27378028 | Electrical Pain | 20 |
|  |  | Mechanical Pain | 17 |
|  |  | Thermal Pain | 10 |
| Daniel et al., 2017 | 27481545 | Delay-To-Match-Sample | 19 |
|  |  | Delay-To-Match-Sample (Non-Verbal) | 23 |
|  |  | Delay-To-Match-Sample (Verbal) | 6 |
| Darda et al., 2019 | 31028924 | Inhibition Imitation | 27 |
| Dastoffo et al., 2017 | 28804467 | Behavioral Prediction Error | 24 |
|  |  | Passive Prediction Error | 5 |
|  |  | Prediction Error | 32 |
| Davis et al., 2009 | 19933145 | Spoken Pseudowords | 50 |
|  |  | Spoken Words | 50 |
| Del Casale et al., 2016 | 26777155 | Hypnotic Analgesic Suggestions (Activation) | 3 |
|  |  | Hypnotic Analgesic Suggestions (Deactivation) | 1 |
| Del Casale et al., 2017 | 28673832 | Empathic Emotional Face Processing | 12 |
| Deng et al., 2018 | 29636070 | Multi-Source Interference | 3 |
| Denny et al., 2012 | 22452556 | Others Processing | 27 |
|  |  | Self Processing | 49 |
| Derbie et al., 2021 | 33880818 | Allocentric Spatial Coding | 6 |
|  |  | Egocentric Spatial Coding | 8 |
|  |  | Spatial Judgment (Allocentric) | 5 |
|  |  | Spatial Judgment (Egocentric) | 4 |
|  |  | Spatial Navigation (Allocentric) | 3 |
|  |  | Spatial Navigation (Egocentric) | 6 |
| Derfuss et al., 2005 | 15846824 | Stroop Color Words | 5 |
|  |  | Task Switching | 8 |
| Devoto et al., 2018 | 30071209 | Hungry | 13 |
| DiVita et al., 2016 | 27177829 | Body Representation | 25 |
| Diekhof et al., 2011 | 21669291 | Extinction (Fear) | 4 |
|  |  | Fear Reappraisal | 31 |
|  |  | Fear Regulation (Placebo) | 13 |
|  |  | Fear Regulation Deactivation (Placebo) | 6 |
| Diekhof et al., 2012 | 22366111 | Reward Anticipation | 12 |
|  |  | Reward Outcome | 17 |
| Dijkstra et al., 2020 | 32407735 | Postural Control | 5 |
| Ding et al., 2020 | 31786222 | Empathy | 36 |
|  |  | Empathy towards Emotional Faces | 8 |
|  |  | Empathy towards Emotional Situations | 13 |
|  |  | Empathy towards Physical Pain | 25 |
| Diveica et al., 2021 | 34742940 | Difficult Theory of Mind | 24 |
|  |  | Easy Theory of Mind | 10 |
|  |  | Empathy | 55 |
|  |  | Empathy for Emotions | 34 |
|  |  | Empathy for Pain | 44 |
|  |  | Explicit Empathy for Emotions | 23 |
|  |  | Explicit Empathy for Pain | 25 |
|  |  | Explicit Moral Reasoning | 12 |
|  |  | False Beliefs | 15 |
|  |  | Implicit Empathy for Emotions | 7 |
|  |  | Implicit Empathy for Pain | 30 |
|  |  | Implicit Moral Reasoning | 4 |
|  |  | Moral Reasoning | 24 |
|  |  | Semantic Control | 23 |
|  |  | Theory of Mind | 29 |
|  |  | Theory of Mind (Normal Difficulty) | 21 |
|  |  | Trait Inference | 22 |
|  |  | Theory of Mind | 26 |
| Dricu et al., 2016 | 27836460 | Explicit Evaluation of Emotional Expressions | 20 |
|  |  | Explicit Evaluation of Facial Expressions | 19 |
|  |  | Explicit Evaluation of Voices Expressions | 8 |
|  |  | Incidental Perception of Emotional Expressions | 3 |
|  |  | Incidental Perception of Facial Expressions | 6 |
|  |  | Passive Perception of Emotional Expressions | 15 |
|  |  | Passive Perception of Facial Expressions | 10 |
|  |  | Emotion Discrimination | 4 |
|  |  | Emotion Labelling | 11 |
|  |  | Emotion Matching | 17 |
|  |  | Emotional Ratings | 2 |
|  |  | Off-Screen Emotion Labelling | 6 |
|  |  | On-Screen Emotion Labelling | 5 |
| Duerden et al., 2011 | 22131304 | Noxious Cold Pain | 64 |
|  |  | Noxious Heat Pain | 61 |
|  |  | Noxious Pain | 48 |
|  |  | Noxious Pain (Cutaneous) | 17 |
|  |  | Noxious Pain (Left Side) | 48 |
|  |  | Noxious Pain (Muscle) | 21 |
|  |  | Noxious Pain (Right Side) | 48 |
| Dugré et al., 2018 | 29761060 | Anticipation Punishment | 68 |
|  |  | Outcome Punishment | 12 |
| Dugré et al., 2021 | 10.1101/2021.05.12.21257119 | Frustration Non-Reward (Activation) | 7 |
|  |  | Frustration Non-Reward (Deactivation) | 4 |
|  |  | Retaliatory Behaviors | 7 |
| Enge et al., 2020 | 32304886 | Language Comprehension (ALE) | 19 |
|  |  | Language Comprehension (SDM) | 6 |
| Eres et al., 2017 | 28724332 | Affective Morality | 8 |
|  |  | Cognitive Morality | 10 |
|  |  | Distal Morality | 4 |
|  |  | Explicit Morality | 5 |
|  |  | Implicit Morality | 3 |
|  |  | Morality | 6 |
|  |  | Other's Morality | 4 |
|  |  | Proximal Morality | 4 |
|  |  | Self Morality | 5 |
| Erickson et al., 2014 | 24996043 | Conflicting Audiovisual Speech | 17 |
|  |  | Validating Audiovisual Speech | 7 |
| Eun Han et al., 2018 | 29496487 | Food Craving DecisionMaking | 6 |
|  |  | Food Decision Making (Craving Regulation) | 7 |
|  |  | Food Regulation Task | 5 |
| Falcone et al., 2018 | 10.1007s12671-018-0884-5 | Mindful Meditation | 8 |
|  |  | Mindful Meditation (Experts) | 6 |
|  |  | Mindful Meditation (Novice) | 3 |
| Fallon et al., 2020 | 32608498 | Empathy for Pain | 25 |
| Fan et al., 2011 | 20974173 | Empathy | 12 |
| Farkas et al., 2021 | 33836303 | Auditory-Laughter Humor | 8 |
|  |  | Humor | 15 |
|  |  | Picture Humor | 16 |
|  |  | Text Humor | 15 |
| Faroqi-Shah et al., 2018 | 30120847 | Nouns | 1 |
|  |  | Verbs | 6 |
| Farrell et al., 2005 | 15846813 | Left Upper Limb Heat | 18 |
|  |  | Right Upper Limb Heat | 16 |
| Fede et al., 2020 | 30706370 | Moral Brain | 20 |
| Feng et al., 2014 | 25327760 | Fairness (Ultimatum Game) | 9 |
|  |  | Unfairness (Ultimatum Game) | 29 |
| Feng et al., 2018 | 29496613 | Emotional Stroop | 3 |
| Feng et al., 2021 | 33781834 | Social Interactions | 9 |
|  |  | Social Prediction Error | 4 |
| Ferstl et al., 2008 | 17557297 | Coherence Building | 10 |
|  |  | Contextual Language Integration | 8 |
|  |  | Language Processing | 10 |
| Filgueiras et al., 2018 | 29260381 | Kinesthetic (Sport Imagery) | 7 |
|  |  | Visual (Sport Imagery) | 6 |
| Filkowski et al., 2016 | 27988321 | Men Emotional Processing | 21 |
|  |  | Women Emotional Processing | 14 |
| Fischer et al., 2020 | 33099860 | Memory-Guided Attention | 4 |
| Flannery et al., 2020 | 31872334 | Reward Processing | 87 |
| Fouragnan et al., 2018 | 29575249 | Reward Prediction Error (Negative) | 22 |
|  |  | Reward Prediction Error (Positive) | 8 |
|  |  | Reward Prediction Error (Signed) | 5 |
|  |  | Reward Prediction Error (Surprise) | 18 |
| Fox et al., 2013 | 23908622 | REM Sleep (Activation) | 10 |
|  |  | REM Sleep (Deactivation) | 7 |
| Fox et al., 2015 | 25725466 | Spontaneous Thought | 13 |
| Fox et al., 2016 | 27032724 | Focused Attention Meditation (Activation) | 2 |
|  |  | Focused Attention Meditation (Deactivation) | 2 |
|  |  | Loving & Compassion-Meditation (Activation) | 3 |
|  |  | Mantra-Meditation (Activation) | 7 |
|  |  | Mantra-Meditation (Deactivation) | 1 |
|  |  | Open Monitoring-Meditation (Activation) | 5 |
|  |  | Open Monitoring-Meditation (Deactivation) | 1 |
| Frank et al., 2014 | 24984244 | Emotional Downregulation | 12 |
|  |  | Emotional Response | 3 |
|  |  | Emotional Upregulation | 8 |
| Freitas et al., 2018 | 30344470 | Familiar Music | 37 |
|  |  | Unfamiliar Music | 15 |
| Friebel et al., 2011 | 21798355 | Experimentally-induced Pain | 17 |
|  |  | Non-Thermal Pain | 11 |
|  |  | Thermal Pain | 20 |
| Fu et al., 2021 | 33549749 | Nocebo Hyperalgesia | 3 |
|  |  | Placebo Analgesia | 3 |
| Fullana et al., 2016 | 26122585 | CS+ minus CS- | 41 |
|  |  | CS+ minus CS- (Deactivation) | 42 |
| Fullana et al., 2018 | 29530516 | Extinction Learning (Fear) | 21 |
|  |  | Extinction Recall (Fear) | 39 |
|  |  | Extinction Recall (Same Fear Context) | 15 |
|  |  | Late Extinction (Fear) | 3 |
| FusarPoli et al., 2009b | 19348735 | Emotional Face Processing | 27 |
| Fusar-Poli et al., 2009 | 19949718 | Facial Angry | 11 |
|  |  | Facial Disgusted | 11 |
|  |  | Facial Emotions | 14 |
|  |  | Facial Fear | 11 |
|  |  | Facial Happy | 10 |
|  |  | Facial Neutral | 15 |
|  |  | Facial Sad | 3 |
| Gabay et al., 2014 | 25454357 | Fairness (Ultimatum Game) | 6 |
|  |  | Response (Ultimatum Game) | 12 |
| Gao et al., 2019 | 31255920 | Audiovisual Affective Processing | 6 |
| García-García et al., 2016 | 27168344 | Disgusting Stimuli | 8 |
|  |  | Explicit Negative Stimuli | 5 |
|  |  | Implicit Negative Stimuli | 10 |
|  |  | Negative Faces | 8 |
|  |  | Negative Images | 7 |
|  |  | Negative Stimuli | 10 |
|  |  | Negative Stimuli (Men) | 5 |
|  |  | Negative Stimuli (Older Adults) | 3 |
|  |  | Negative Stimuli (Women) | 6 |
|  |  | Negative Stimuli (Young Adults) | 6 |
|  |  | Negative Words | 3 |
|  |  | Passive Viewing Negative Stimuli | 9 |
| Garrigan et al., 2017 | 27842284 | Moral Evaluations | 4 |
|  |  | Moral Response Decisions | 4 |
| Garrison et al., 2013 | 23567522 | Instrumental (PE) | 21 |
|  |  | Pavlovian (PE) | 20 |
|  |  | Prediction Error | 33 |
|  |  | Punishment (PE) | 26 |
|  |  | Reward (PE) | 21 |
| Gavazzi et al., 2020 | 32748318 | Proactive Inhibitory Process | 10 |
|  |  | Reactive Inhibitory Process | 5 |
|  |  | Reactive Inhibitory Process | 15 |
| Gianaros et al., 2009 | 19410652 | Blood Pressure | 29 |
| Gifuni et al., 2017 | 27704409 | GuiltFeelings | 17 |
| Gilead et al., 2013 | 23063843 | Intention | 5 |
| GonenYaacovi et al., 2013 | 23966927 | Creative Combination Generation | 38 |
|  |  | Creativity | 52 |
|  |  | Unusual Creative Generation | 29 |
| Gordon et al., 2018 | 30452442 | Passive Listening | 6 |
| Groenendijk et al., 2021 | 33534812 | Micturition Control | 11 |
|  |  | Pelvic Muscle Contraction | 18 |
| Grosbras et al., 2011 | 21391275 | Body Movement | 19 |
|  |  | Face Movement | 19 |
|  |  | Hand Movement | 13 |
|  |  | Object Hand Move | 25 |
|  |  | Static Body Perception | 9 |
|  |  | Static Face Perception | 22 |
| Gu et al., 2012 | 22961548 | Empathetic Pain Perception | 28 |
| Gu et al., 2014 | 23749500 | Empathy | 41 |
|  |  | Empathy for Negative Emotions | 18 |
|  |  | Empathy for Positive Emotions | 14 |
|  |  | Pain Empathy | 28 |
| Gu et al., 2019 | 30807783 | Reward Anticipation (Money) | 7 |
|  |  | Reward Anticipation (Social) | 7 |
| Halani et al., 2019 | 31816125 | Bladder Filling | 5 |
|  |  | Rectal Distention | 8 |
| Han et al., 2017 | 10.108003057240.2016.1262834 | Moral Judgment | 13 |
|  |  | Moral Sensibility | 35 |
|  |  | Morality | 26 |
| Hardwick et al., 2013 | 23194819 | Motor Learning | 13 |
|  |  | Sensori-Motor Tasks | 11 |
|  |  | Serial Response Time Tasks | 8 |
| Hardwick et al., 2018 | 30098990 | Action Observation | 39 |
|  |  | Motor Imagery | 30 |
|  |  | Movement Execution | 37 |
| Harvie et al., 2019 | 30069958 | Bladder Voiding | 5 |
| Heard et al., 2020 | 31783081 | Merge (Syntax) | 10 |
|  |  | Movement (Syntax) | 10 |
|  |  | Reanalysis (Syntax) | 9 |
|  |  | Rhythm | 28 |
|  |  | Syntax | 19 |
| Hetu et al., 2013 | 23583615 | Motor Imagery | 34 |
| Hill et al., 2014 | 25042764 | DMTS+N-Back (Female) | 103 |
|  |  | DMTS+N-Back (Male) | 80 |
|  |  | Working Memory (Female) | 42 |
|  |  | Working Memory (Male) | 39 |
|  |  | Working Memory (Men) | 27 |
|  |  | Working Memory (Women) | 31 |
| Hobeika et al., 2016 | 27012301 | Analogical Reasoning | 13 |
|  |  | Semantic Analogy | 7 |
|  |  | Visuospatial Analogy | 4 |
|  |  | Visuospatial Matrix Problems | 14 |
| Hoffman et al., Morcom et al., 2018 | 29183684 | Semantic (Older Adults) | 9 |
|  |  | Semantic (Young Adults) | 19 |
| Hu et al., 2016 | 26695384 | Self Face Processing | 11 |
|  |  | Self Reference Processing | 7 |
| Huang et al., 2020 | 32283275 | Model Free Decision-Making | 3 |
|  |  | Model-Based Decision-Making | 9 |
| Huang et al., 2020 | 31830505 | Incongruent (Stroop) | 21 |
| Huerta et al., 2014 | 24174404 | Food Odor Cues | 12 |
|  |  | Food Taste Cues | 15 |
|  |  | Food Visual Cues | 12 |
| Hung et al., 2018 | 29923271 | Cognitive Inhibition | 14 |
|  |  | Emotional Inhibition | 4 |
|  |  | Response Inhibition | 6 |
| Ishibashi et al., 2016 | 27362967 | Tool Cognition | 16 |
| Jamadar et al., 2013 | 24137150 | Antisaccades | 17 |
|  |  | Prosaccades | 8 |
| Janacszek et al., 2020 | 31765803 | Sequence Learning | 14 |
| Jauniaux et al., 2019 | 31393982 | 1st Person Pain Empathy | 25 |
|  |  | 3rd Person Pain Empathy | 7 |
|  |  | Emotional-Communicative Pain Empathy | 12 |
|  |  | Other-Oriented Pain Empathy | 20 |
|  |  | Pain Empathy | 62 |
|  |  | Self-Oriented Pain Empathy | 13 |
|  |  | Somatosensory Pain Empathy | 33 |
|  |  | Stimuli-Oriented Pain Empathy | 12 |
| Jensen et al., 2016 | 26871535 | Noxious Pain | 28 |
| Jirak et al., 2010 | 20739194 | Embodiment of language | 21 |
| Johnston et al., 2019 | 31155815 | Auditory Feedback Error | 6 |
|  |  | Sensory Feedback Error | 5 |
|  |  | Visual Feedback Error | 6 |
| Kalisch et al., 2009 | 19539645 | Reappraisal of Emotion | 18 |
| Kaufmann et al., 2011 | 21761997 | Non-Symbolic Numbers (Children) | 15 |
|  |  | Symbolic Numbers (Children) | 16 |
| Kenzie et al., 2017 | 28801769 | Illusion of Movement | 16 |
|  |  | Imposed Movement | 18 |
|  |  | Proprioceptive Stimuli | 22 |
| Keuken et al., 2014 | 24994979 | Perceptual Decision (Hard Trials) | 22 |
|  |  | Perceptual Decision Task | 12 |
|  |  | Reward Anticipation | 17 |
| Kim et al., 2010 | 20097295 | Familiarity | 21 |
|  |  | Familiarity Strength | 9 |
|  |  | Recollection | 17 |
| Kim et al., 2011 | 21391260 | Context Switching | 19 |
|  |  | Perceptual Switching | 17 |
|  |  | Response Switching | 13 |
| Kim et al., 2011 | 20869446 | Subsequent Forgetting | 10 |
|  |  | Subsequent Memory | 11 |
|  |  | Subsequent Pictoral Associative Memory | 12 |
|  |  | Subsequent Pictoral Memory | 15 |
|  |  | Subsequent Verbal Associative Memory | 11 |
|  |  | Subsequent Verbal Forgetting | 10 |
|  |  | Subsequent Verbal Memory | 10 |
| Kim et al., 2012 | 22446489 | Autobiographical Memory Retrieval | 15 |
|  |  | Autobiographical Memory Retrieval | 9 |
| Kim et al., 2014 | 23900833 | Auditory Oddball | 19 |
|  |  | Autobiographical Retrieval | 16 |
|  |  | Laboratory-based Recollection | 7 |
|  |  | Subsequent Memory | 12 |
|  |  | Visual Oddball | 17 |
| Kim et al., 2016 | 26562053 | Episodic Retrieval (Old) | 13 |
|  |  | Episodic Retrieval (Recall) | 12 |
|  |  | Semantic Retrieval | 11 |
|  |  | Semantic Retrieval (Words) | 6 |
| Kim et al., 2017 | 28009076 | Faces Repetition Suppression | 6 |
|  |  | Objects Repetition Suppression | 14 |
|  |  | Repetition Enhancement (Memory) | 12 |
|  |  | Repetition Suppression (Memory) | 10 |
|  |  | Scenes Repetition Suppression | 11 |
|  |  | Words Repetition Suppression | 8 |
| Kim et al., 2018 | 29456134 | Repetition Enhancement (Memory) | 5 |
|  |  | Retrieval Success (Memory) | 8 |
|  |  | Subsequent Forgetting | 7 |
| Kim et al., 2019 | 31373730 | Encoding | 8 |
|  |  | Maintenance | 8 |
|  |  | Repetition Suppression (Memory) | 6 |
| Kim et al., 2019 | 31034858 | Retrieval | 6 |
|  |  | Retrieval Success | 10 |
|  |  | Subsequent Memory | 8 |
| Kim et al., 2020 | 31697955 | Brand Love | 9 |
|  |  | Compassion | 25 |
|  |  | Maternal Love | 18 |
| King et al., 2014 | 24927986 | Dynamic Grip | 5 |
|  |  | Power Grip | 10 |
|  |  | Precision Grip | 9 |
|  |  | Static Grip | 13 |
| King et al., 2019 | 31283953 | Innocuous Cold Exposure | 3 |
|  |  | Noxious Cold Exposure | 11 |
| Knutson et al., 2008 | 18829428 | Gain Anticipation | 10 |
|  |  | Loss Anticipation | 7 |
| Kober et al., 2008 | 18579414 | Emotion Structure | 67 |
| Koch et al., 2018 | 30412701 | Alternative Emotional Reappraisal | 22 |
|  |  | Emotional Action Control | 2 |
|  |  | Single Emotional Reappraisal Strategy | 24 |
| Koelsch et al., 2014 | 24552785 | Music-evoked Emotions | 13 |
| Koelsch et al., 2020 | 32898679 | Music-Evoked Emotions | 23 |
| Kogler et al., 2015 | 26123376 | Physiological Stress | 24 |
|  |  | Psychosocial Stress | 4 |
| Kogler et al., 2020 | 32562973 | Affective Empathy | 5 |
|  |  | Cognitive Empathy | 8 |
|  |  | Empathy | 7 |
|  |  | Empathy For Emotions | 9 |
|  |  | Empathy For Pain | 9 |
| Kohn et al., 2014 | 24220041 | Emotion Regulation | 8 |
| Krain et al., 2006 | 16632383 | Ambiguous Decision-Making | 18 |
|  |  | Decision-Making | 24 |
|  |  | Risky Decision-Making | 16 |
| Krall et al., 2015 | 24915964 | False Beliefs | 7 |
|  |  | Reorienting Attention | 7 |
| Kraynak et al., 2018 | 30067939 | Peripheral Inflammation Physiology | 10 |
| Kubit et al., 2013 | 23847497 | Reorienting | 19 |
|  |  | Target Detection | 19 |
|  |  | Theory Of Mind | 12 |
| Kuhn et al., 2011 | 21599838 | Sexual Arousal | 16 |
|  |  | Sexual Arousal (Penile Turgidity) | 8 |
| Kuhn et al., 2012 | 22406357 | Subjective Liking | 9 |
|  |  | Subjective Pleasantness | 9 |
|  |  | Subjective Pleasantness (Inside Scanner) | 14 |
| Kuhn et al., 2013 | 23362184 | Non Spatial (Retrieval) | 17 |
|  |  | Non-Spatial (Encoding) | 16 |
|  |  | Spatial Navigation (Encoding) | 11 |
|  |  | Spatial Navigation (Retrieval) | 10 |
| Kurkela et al., 2016 | 26683385 | Encoding False Memory | 2 |
|  |  | False Retrieval | 15 |
|  |  | Perceptual False Memory Retrieval | 4 |
|  |  | Pictoral False Memory Retrieval | 3 |
|  |  | Retrieval of False Memory | 6 |
|  |  | Semantic False Memory Retrieval | 6 |
|  |  | Verbal False Memory Retrieval | 6 |
| Kwok et al., 2015 | 25196948 | Processing Lexical Tone | 2 |
| Lacroix et al., 2015 | 26321976 | Music Discrimination | 11 |
|  |  | Music Error Detection | 14 |
|  |  | Music Memory | 16 |
|  |  | Passive Listening to Music | 10 |
|  |  | Passive Listening to Speech | 10 |
|  |  | Speech Detection | 16 |
|  |  | Speech Discrimination | 10 |
|  |  | Speech Memory | 14 |
| Laird et al., 2005 | 15846823 | Manual Stroop | 5 |
|  |  | Stroop Color Words | 13 |
|  |  | Verbal Stroop | 8 |
|  |  | Task Deactivation | 9 |
| Laird et al., 2010 | 20197097 | Encoding | 22 |
|  |  | Recall | 16 |
| Lamm et al., 2011 | 20946964 | Empathy for Pain | 19 |
|  |  | Empathy for Pain (Cues) | 6 |
|  |  | Empathy for Pain (Pictures) | 11 |
| Lamp et al., 2019 | 30687211 | Left Hand Stimulation | 7 |
|  |  | Right Hand Stimulation | 11 |
| Langner et al., 2018 | 29730485 | Cognitive Action Regulation | 21 |
|  |  | Cognitive Emotion Regulation | 17 |
| Langner et al., 2014 | 23163491 | Auditory (Vigilant Attention) | 5 |
|  |  | Duration (Vigilant Attention) | 9 |
|  |  | Go No-Go (Vigilant Attention) | 8 |
|  |  | No Overt Response (Vigilant Attention) | 7 |
|  |  | Overt Response (Vigilant Attention) | 10 |
|  |  | Predictable Occurrence (Vigilant Attention) | 12 |
|  |  | Simple Reaction (Vigilant Attention) | 4 |
|  |  | Unpredictable Occurrence (Vigilant Attention) | 4 |
|  |  | Vigilant Attention | 37 |
|  |  | Visual (Vigilant Attention) | 9 |
| Lanz et al., 2011 | 21373762 | Allodynia Pain | 34 |
|  |  | Hyperalgesia | 29 |
|  |  | Mechanical & Electrical Pain | 27 |
|  |  | Mechanical Allodynia Hyperalgesia | 42 |
|  |  | Pain | 33 |
|  |  | Thermal Allodynia Hyperalgesia | 25 |
|  |  | Thermal Pain | 32 |
| Lee et al., 2012 | 19270039 | Evaluation of Other's Emotions | 12 |
|  |  | Evaluation of Own Emotions | 11 |
|  |  | Stimulus-focused Evaluation | 21 |
| Lee et al., 2017 | 28237296 | Cool Executive Functions (Adults) | 41 |
|  |  | Cool Executive Functions (Children) | 12 |
|  |  | Reward Decision-Making (Adults) | 34 |
|  |  | Reward Decision-Making (Children) | 20 |
|  |  | Theory of Mind (Adults) | 37 |
|  |  | Theory of Mind (Children) | 10 |
| Lee et al., 2018 | 30212509 | Emotional Downregulation | 15 |
|  |  | Working Memory | 16 |
| Lee et al., 2020 | 33192395 | Repetition Enhancement (Memory) | 5 |
|  |  | Repetition Suppression (Conceptual Priming) | 25 |
|  |  | Repetition Suppression (Memory) | 26 |
|  |  | Repetition Suppression (Perceptual Priming) | 6 |
| Levy et al., 2011 | 21486295 | Motor Inhibition | 21 |
|  |  | Reflexive Reorienting | 24 |
| Levy et al., 2012 | 22766486 | Subjective Value | 2 |
| Li et al., 2015 | 26318367 | Encoding (Older) | 12 |
|  |  | Encoding (Young) | 11 |
|  |  | Executive (Older) | 11 |
|  |  | Executive (Young) | 4 |
|  |  | Retrieval (Older) | 8 |
|  |  | Retrieval (Young) | 15 |
| Li et al., 2017 | 29017916 | Conflict | 17 |
|  |  | Stimulus-Response Conflict | 6 |
|  |  | Stimulus-Stimulus Conflict | 12 |
| Li et al., 2019 | 30686987 | Large-scale Spatial Abilities | 31 |
|  |  | Small-scale Spatial Abilities | 25 |
| Li et al., 2021 | 34129948 | Allocentric Spatial Reference | 13 |
|  |  | Egocentric Spatial Reference | 6 |
|  |  | Environmental Navigational Space | 20 |
| Li et al., 2021 | 10.1017S136672892000070X | L1 | 10 |
|  |  | L2 | 10 |
| Li et al., 2021 | 34129948 | Spatial Navigation | 17 |
|  |  | Vista Navigational Space | 3 |
| Lindquist et al., 2012 | 22617651 | Anger Experience | 1 |
|  |  | Anger Perception | 13 |
|  |  | Disgust Experience | 7 |
|  |  | Disgust Perception | 3 |
|  |  | Fear Experience | 2 |
|  |  | Fear Perception | 6 |
|  |  | Happy Experience | 1 |
|  |  | Sad Experience | 7 |
|  |  | Sad Perception | 3 |
| Linortner et al., 2014 | 24836898 | Left Ankle Movement | 5 |
|  |  | Right Ankle Movement | 3 |
| Lisofsky et al., 2014 | 24929201 | Deception | 6 |
| Liu et al., 2011 | 21185861 | Anticipation Salient Stimuli | 21 |
|  |  | Anticipation of Reward (Parametric) | 11 |
|  |  | Negative Reward | 13 |
|  |  | Negative Reward (Parametric) | 14 |
|  |  | Outcome Salient Stimuli | 17 |
|  |  | Positive Reward (Parametric) | 18 |
|  |  | Positive Reward | 18 |
|  |  | Reward Evaluation (Parametric) | 8 |
|  |  | Reward Outcome (Parametric) | 14 |
|  |  | Salient Stimuli Processing | 34 |
| Liu et al., 2016 | 27295606 | Language | 8 |
| Lohse et al., 2014 | 24831923 | Motor acquisition (Long Time - Decreases) | 17 |
|  |  | Motor acquisition (Long Time - Increases) | 9 |
|  |  | Motor acquisition (Medium Time - Decreases) | 18 |
|  |  | Motor acquisition (Medium Time - Increases) | 12 |
|  |  | Motor acquisition (Short Time - Decreases) | 14 |
|  |  | Motor acquisition (Short Time - Increases) | 15 |
| Long et al., 2020 | 32312660 | Sexual Non-Preference | 5 |
|  |  | Sexual Non-Preference (Men) | 4 |
|  |  | Sexual Preference | 3 |
|  |  | Sexual Preference (Men) | 3 |
|  |  | Sexual Stimuli | 5 |
| Lopez et al., 2012 | 22516007 | Caloric Vestibular Stimulation | 20 |
|  |  | Galvanic Vestibular Stimulation | 14 |
|  |  | Left Ear Stimulation | 11 |
|  |  | Right Ear Stimulation | 16 |
|  |  | Saccular Vestibular Stimulation | 4 |
| Luk et al., 2011 | 24795491 | Language Switching (Biliingual) | 7 |
| Luo et al., 2017 | 29064617 | Downward Social Comparison | 3 |
|  |  | Upward Social Comparison | 3 |
| MacCormack et al., 2020 | 36043210 | Affect (Older) | 21 |
|  |  | Affect (Young) | 32 |
| Maillet et al., 2014 | 24973756 | Successful Encoding | 16 |
|  |  | Unsuccessful Encoding | 17 |
| Maki-Marttunen et al., 2019 | 30659515 | Working Memory (AX-CPT) | 11 |
| Makovac et al., 2020 | 31790922 | Perseverative Cognition | 36 |
| Mar et al., 2011 | 21126178 | Narrative Comprehension | 15 |
|  |  | Non-Story-based (ToM) | 23 |
|  |  | Story-based (ToM) | 13 |
| Marchesotti et al., 2017 | 28321973 | Hand Motor Imagery | 30 |
| Marstaller et al., 2014 | 10.1016j.jneuroling.2014.04.003 | Co-Speech Gesture | 8 |
| Martinelli et al., 2012 | 22359397 | Conceptual Self | 10 |
|  |  | Episodic Autobiographical Memory | 13 |
|  |  | Semantic Autobiographical Memory | 16 |
| Martins et al., 2020 | 33421544 | Anticipation Social Punishment | 19 |
|  |  | Outcome Social Reward | 16 |
|  |  | Receipt Social Punishment | 11 |
| MasHerrero et al., 2021 | 33440196 | Food Reward | 5 |
|  |  | Music Reward | 4 |
| McDermott et al., 2009 | 19159634 | Retrieval Memory (Autobiographical-Based) | 13 |
|  |  | Retrieval Memory (Laboratory-Based) | 18 |
| McKenna et al., 2017 | 28439231 | Inhibition (12 yrs Children) | 18 |
|  |  | Inhibition (Children) | 20 |
|  |  | Switching (Children) | 4 |
|  |  | Updating (Children) | 25 |
| McMillan et al., 2007 | 17280644 | N-Back | 6 |
|  |  | N-Back (Memory Load) | 9 |
| McNorgan et al., 2012 | 23087637 | Auditory Imagery | 10 |
|  |  | General Imagery | 9 |
|  |  | Gustatory Imagery | 10 |
|  |  | Motor Imagery | 5 |
|  |  | Tactile Imagery | 3 |
| McNorgan et al., 2015 | 25524364 | Lexical Decision | 8 |
|  |  | Naming | 2 |
|  |  | Pseudowords | 4 |
|  |  | Pseudowords Lexical Decision | 4 |
|  |  | Words | 4 |
|  |  | Words Lexical Decision | 7 |
|  |  | Words Naming | 1 |
| Mechias et al., 2010 | 19786103 | Instructed Fear | 15 |
|  |  | Uninstructed Fear | 29 |
| Mencarelli et al., 2019 | 31179585 | Auditory N-Back | 12 |
|  |  | Faces N-Back | 14 |
|  |  | Letter N-Back | 26 |
|  |  | N-Back | 32 |
|  |  | N-Back (Activations) | 13 |
|  |  | N-Back (Deactivations) | 5 |
|  |  | Non-Verbal N-Back | 17 |
|  |  | Number N-Back | 17 |
|  |  | Object Images N-Back | 11 |
|  |  | Spatial N-Back | 20 |
|  |  | Verbal N-Back | 26 |
| Mende-Siedlecki et al., 2013 | 22287188 | Negative Face Evaluation | 2 |
|  |  | Positive Face Evaluation | 7 |
| Meneguzzo et al., 2014 | 25593703 | Subliminal Stimulation | 3 |
|  |  | Supraliminal Stimulation | 2 |
| Merritt et al., 2021 | 33760100 | In-group Social Cognition | 9 |
|  |  | Out-group Social Cognition | 5 |
| Messina et al., 2015 | 26217277 | Emotion Baseline | 2 |
|  |  | Reappraisal | 6 |
| Messina et al., 2021 | 33475715 | Emotion Acceptance | 15 |
| Mitricheva et al., 2019 | 31308220 | Erotic Visual Stimulation (Bisexuals) | 27 |
|  |  | Erotic Visual Stimulation (Heterosexuals) | 34 |
|  |  | Erotic Visual Stimulation (Homosexuals) | 32 |
|  |  | Erotic Visual Stimulation (Transsexuals) | 14 |
|  |  | Sexual Arousal | 22 |
|  |  | Sexual Cues (Men) | 16 |
|  |  | Sexual Cues (Women) | 4 |
| Mohr et al., 2010 | 20463224 | Anticipation of Risk | 14 |
|  |  | Decision of Risk | 11 |
|  |  | Risk | 15 |
| Molenberghs et al., 2009 | 19580913 | Mirror Imitation | 10 |
| Molenberghs et al., 2012 | 21782846 | Auditory (Mirror System) | 14 |
|  |  | Classical Studies (Mirror System) | 25 |
|  |  | Emotional Expressions (Mirror System) | 13 |
|  |  | Mirror System Functions | 28 |
|  |  | Somatosensory (Mirror System) | 2 |
| Molenberghs et al., 2016 | 27073047 | Affective Theory of Mind | 17 |
|  |  | Animations (Theory of Mind) | 19 |
|  |  | Cartoons (Theory of Mind) | 9 |
|  |  | Cognitive Theory of Mind | 19 |
|  |  | Explicit (Theory of Mind) | 31 |
|  |  | Implicit (Theory of Mind) | 27 |
|  |  | Interactive Games (Theory of Mind) | 5 |
|  |  | Photographs (Theory of Mind) | 10 |
|  |  | RMET (Theory of Mind) | 21 |
|  |  | Stories (Theory of Mind) | 18 |
|  |  | Theory of Mind | 43 |
|  |  | Verbal (Theory of Mind) | 23 |
|  |  | Videos (Theory of Mind) | 7 |
|  |  | Visual (Theory of Mind) | 38 |
| Morandini et al., 2020 | 33035933 | Vigilant Attention (Children) | 5 |
| Morawetz et al., 2017 | 27894828 | Attention-focused Strategies (Emotion) | 3 |
|  |  | Emotion Reappraisal | 39 |
|  |  | Emotion Regulation (Image Viewing) | 37 |
|  |  | Emotion Regulation (Other Stimuli) | 14 |
|  |  | Goal Downregulation (Emotion) | 44 |
|  |  | Goal Upregulation (Emotion) | 11 |
|  |  | Response-focused Strategies (Emotion) | 5 |
| Morawetz et al., 2020 | 32659287 | Emotion Regulation | 36 |
| Morelli et al., 2015 | 25554428 | Personal Reward | 36 |
|  |  | Vicarious Reward | 41 |
| Morrison et al., 2016 | 26873519 | Affective Touch | 4 |
|  |  | Discriminative Touch | 11 |
| Morriss et al., 2019 | 30550858 | Basic Threat and Reward Uncertainty | 2 |
|  |  | Uncertainty during Associative Learning | 6 |
|  |  | Uncertainty under Decision-Making | 2 |
| Muller et al., 2018 | 29665467 | Face Processing | 25 |
| Murphy et al., 2003 | 14672157 | Anger Affect Program | 2 |
|  |  | Disgust Affect Program | 3 |
|  |  | Fear Affect Program | 2 |
|  |  | Happy Affect Program | 2 |
|  |  | Sad Affect Program | 2 |
| Murphy et al., 2018 | 10.1016j.jneuroling.2018.08.005 | Lexical Decision Task | 3 |
|  |  | Lexical Non-Tonal Tone | 1 |
|  |  | Lexical Tonal Tone | 6 |
|  |  | Phoneme | 3 |
|  |  | Sentence Prosody | 9 |
|  |  | Single Word Reading | 11 |
|  |  | Word Prosody | 4 |
| Murray et al., 2012 | 22230705 | Others Reflection | 12 |
|  |  | Self Reflection | 16 |
| Murty et al., 2010 | 20688087 | Successful Emotion Encoding | 12 |
| Mwilambwe-Tshilobo et al., 2021 | 33359341 | Social Exclusion (Cyberball) | 6 |
| Nani et al., 2019 | 31418337 | Subseconds (Motor) | 48 |
|  |  | Subseconds (Perceptual) | 26 |
|  |  | Supraseconds (Motor) | 17 |
|  |  | Supraseconds (Perceptual) | 9 |
| Navarro et al., 2018 | 30442593 | Car Driving | 23 |
|  |  | Operational Car Driving | 4 |
|  |  | Strategic Car Driving | 8 |
|  |  | Tactical Car Driving | 3 |
| Nee et al., 2007 | 17598730 | Flanker | 2 |
|  |  | Go No-Go | 9 |
|  |  | Resolution Interference | 12 |
|  |  | Stimulus-Response | 8 |
|  |  | Stroop | 10 |
| Nee et al., 2013 | 22314046 | Executive Process (Working Memory) | 49 |
|  |  | Shifting (Working Memory) | 6 |
|  |  | Spatial (Working Memory) | 8 |
|  |  | Verbal Spatial (Working Memory) | 6 |
| Neumann et al., 2008 | 17390315 | Stroop | 13 |
| Neumann et al., 2015 | 26467981 | Expertise Auditory | 7 |
|  |  | Expertise NoMusic | 2 |
|  |  | Expertise Visual | 1 |
| Niendam et al., 2012 | 22282036 | Executive Processing | 31 |
|  |  | Flexibility Processing | 37 |
|  |  | Inhibition Processing | 45 |
|  |  | Initiation Behavior | 20 |
|  |  | Working Memory | 31 |
| Nitschke et al., 2016 | 27627877 | Planning | 18 |
|  |  | Planning Complexity | 19 |
| Noori et al., 2016 | 27397863 | Food Cues | 24 |
|  |  | Sexual Cues | 50 |
| Oldham et al., 2018 | 29696725 | Punishment Anticipation | 15 |
|  |  | Reward Anticipation | 20 |
|  |  | Reward Outcome | 8 |
| Ortiz-Teran et al., 2021 | 33801075 | Investment Decision-Making | 4 |
| Ortuño et al., 2011 | 21041067 | Time Estimation | 24 |
| Owen et al., 2005 | 15846822 | N-Back Working Memory | 14 |
|  |  | Non-Verbal N-Back | 18 |
|  |  | Verbal N-Back | 16 |
| Palermo et al., 2015 | 25529840 | Anticipation of Pain | 52 |
| Pando-Naude et al., 2021 | 34675231 | Music Imagery | 26 |
|  |  | Music Perception | 75 |
|  |  | Music Production | 70 |
| Papitto et al., 2020 | 31678500 | Action Execution | 34 |
|  |  | Action Imitation | 26 |
|  |  | Action Observation | 30 |
|  |  | Motor Imagery | 39 |
|  |  | Motor Learning | 19 |
|  |  | Motor Planning | 19 |
| Parro et al., 2018 | 30120846 | Motivated Cognitive Control | 4 |
| Paul et al., 2018 | 29987638 | Maternal Responding (Own Children) | 5 |
| Penner et al., 2020 | Penner, M., Moes, M., & Cecala, A. (2020). Meta-Analysis of the Neural Correlates of Finger Gnosis using Activation Likelihood Estimation. In CogSci. | Finger Gnosis | 15 |
| Petacchi et al., 2005 | 15846816 | Auditory | 11 |
| Pidgeon et al., 2016 | 27781148 | Visual Creativity | 13 |
| Planton et al., 2013 | 23831432 | Handwritting Processes | 19 |
| Poeppl et al., 2014 | 23674246 | Deactivation after Sexual Arousal | 5 |
|  |  | Physiosexual Arousal | 10 |
|  |  | Psychosexual Arousal | 17 |
| Poeppl et al., 2016 | 27742561 | Sexual Stimulation (Women) | 19 |
| Poeppl et al., 2016 et al., 2 | 27339689 | Sexual Non-Preference (Women) | 1 |
|  |  | Sexual Preference (Women) | 4 |
| Pollack et al., 2015 | 25806009 | Reading & Rhyming | 6 |
| Pollack et al., 2018 | 28533112 | Arithmetic (Adults) | 12 |
|  |  | Arithmetic (Children) | 7 |
|  |  | Phonological (Adults) | 14 |
|  |  | Phonological (Children) | 8 |
| Poudel et al., 2020 | 32078973 | Ambiguous Decision-Making | 3 |
|  |  | Decision-Making | 8 |
|  |  | Perceptual Decision-Making | 8 |
|  |  | Risky Decision-Making | 4 |
| Powers et al., 2019 | 30502352 | Distancing from Emotions | 6 |
| Pozzi et al., 2021 | 33268030 | Emotion Reactivity | 8 |
|  |  | Explicit Emotion Regulation | 4 |
|  |  | Implicit Emotion Regulation | 5 |
|  |  | Implicit Emotion Regulation (Adolescents) | 3 |
|  |  | Implicit Regulation of Negative Emotions | 6 |
|  |  | Implicit Regulation of Negative Emotions (Adolescents) | 4 |
|  |  | Implicit Regulation of Negative Emotions (Early Adults) | 3 |
|  |  | Reactivity of Negative Emotions | 5 |
| Prado et al., 2011 | 21568632 | Deductive Reasoning | 26 |
| Puiu et al., 2020 | 32314475 | Go No-Go | 34 |
|  |  | Go No-Go (Adults) | 34 |
|  |  | Go No-Go (Children) | 8 |
|  |  | Response Inhibition | 37 |
|  |  | Response Inhibition (Adults) | 39 |
|  |  | Response Inhibition (Children) | 6 |
|  |  | State Anger | 8 |
|  |  | Stop Signal | 24 |
|  |  | Stop Signal (Adults) | 25 |
|  |  | Stop Signal (Children) | 2 |
| Purcell et al., 2011 | 22013427 | Central Cognition | 7 |
|  |  | Central Peripheral Cognition | 13 |
|  |  | Written Production | 22 |
| Qin et al., 2020 | 32492474 | Exteroceptive Processing | 14 |
|  |  | Familiarity | 7 |
|  |  | Interoceptive Processing | 11 |
|  |  | Mental Self-Processing | 12 |
| Qin et al., 2011 | 21609772 | Familiarity Processing | 4 |
|  |  | Other processing | 4 |
|  |  | Self Processing | 5 |
| Qiu et al., 2019 | 31201830 | Landmark (Spatial Information) | 13 |
|  |  | Map-based Navigational Strategies | 8 |
|  |  | Orientation Cues (Spatial Information) | 13 |
|  |  | Route-based Navigational Strategies | 12 |
|  |  | Spatial Navigation | 20 |
| Rae et al., 2014 | 24128740 | Action Selection | 29 |
|  |  | Action Stopping | 41 |
| Ran et al., 2017 | 29128283 | Predictable Emotions | 13 |
|  |  | Unpredictable Emotions | 2 |
|  |  | Unpredictable Negative Emotions | 3 |
| Ran et al., 2018 | 29733250 | Attachment | 4 |
| Rapp et al., 2012 | 22759997 | Metaphoric Stimuli | 17 |
|  |  | Non-Literal Language | 20 |
|  |  | Novel Metaphors | 10 |
|  |  | Salient Metaphors | 10 |
| Rhoads et al., 2021 | 34160604 | Altruistic Decisions | 17 |
|  |  | Cooperative Decisions | 16 |
|  |  | Equitable Decisions | 9 |
|  |  | Selfish Decisions | 42 |
| Rice et al., 2015 | 25771223 | Conceptual Knowledge Non-Verbal | 12 |
|  |  | Conceptual Knowledge Verbal | 11 |
| Riedel et al., 2018 | 29484767 | Emotion | 15 |
| Roberts et al., 2019 | 31054845 | Itch | 12 |
| Roberts et al., 2020 | 32271923 | Sucrose | 14 |
|  |  | Sweet Taste | 11 |
| Rodd et al., 2015 | 25576690 | Semantic & Syntax | 22 |
|  |  | Semantics | 8 |
|  |  | Syntax | 12 |
| Rotge et al., 2015 | 25140048 | Cyberball | 25 |
|  |  | Social Pain | 28 |
| Rottschy et al., 2012 | 22178808 | Memory Load (WM) | 10 |
|  |  | Non-Verbal (WM) | 3 |
|  |  | Object Identification (WM) | 3 |
|  |  | Object Location (WM) | 5 |
|  |  | Task-Effect (WM) | 5 |
|  |  | Verbal (WM) | 1 |
|  |  | Working Memory | 25 |
| Sabatinelli et al., 2011 | 20951215 | Emotional Faces | 15 |
|  |  | Emotional Scenes | 13 |
| Sadigh-Eteghad et al., 2014 | 24950901 | Cognition (Elderly) | 47 |
| Sakrekia et al., 2016 | 27484872 | Stable Affordance | 15 |
|  |  | Variable Affordance | 8 |
| Samson et al., 2011 | 21833294 | Music | 10 |
|  |  | Noise | 7 |
|  |  | Tone | 2 |
|  |  | Vocal Sounds | 4 |
| Santarnecchi et al., 2017 | 10.1016j.intell.2017.04.008 | Complexity (Intelligence) | 26 |
|  |  | Fluid Intelligence | 30 |
|  |  | Rule Application (Intelligence) | 31 |
|  |  | Rule Inference (Intelligence) | 29 |
|  |  | Verbal Intelligence | 23 |
|  |  | Vistospatial Intelligence | 10 |
| Santos et al., 2016 | 27898705 | Facial Trustworthiness | 2 |
|  |  | Facial Untrustworthiness | 6 |
| Satpute et al., 2015 | 26696928 | Auditory | 31 |
|  |  | Gustatory | 23 |
|  |  | Olfactory | 19 |
|  |  | Somato-sensory | 21 |
|  |  | Visual Faces | 52 |
|  |  | Visual Pictures | 37 |
| Schilbach et al., 2008 | 18434197 | Task Deactivation | 4 |
| Schilbach et al., 2012 | 22319593 | Emotion Processing | 7 |
|  |  | Social Cognition Tasks | 8 |
|  |  | Unconstrained Cognition (DMN) | 9 |
| Schirmer et al., 2012 | 22732561 | Environmental Sounds | 24 |
|  |  | Human Vocalizations | 15 |
|  |  | Music | 16 |
| Schirmer et al., 2017 | 29186621 | Emotional Faces | 9 |
|  |  | Emotional Voices | 9 |
| Schulz et al., 2016 | 28080975 | Heart Interoception | 12 |
| Schurz et al., 2013 | 24198773 | False Belief Reasoning | 27 |
|  |  | Visual Perspective Taking | 23 |
| Schurz et al., 2014 | 24486722 | False Beliefs (ToM) | 26 |
|  |  | RMET (ToM) | 27 |
|  |  | Rational Actions (ToM) | 22 |
|  |  | Social Animations (ToM) | 32 |
|  |  | Strategic Games (ToM) | 15 |
|  |  | Theory of Mind | 40 |
|  |  | Trait Judgments (ToM) | 22 |
| Sebastian et al., 2011 | 10.1017S0142716411000075 | L1 Bilingual | 14 |
|  |  | L1 Non-Bilingual | 11 |
|  |  | L2 Bilingual | 10 |
|  |  | L2 Non-Bilingual | 17 |
| Seghezzi et al., 2019 | 31629195 | Self-Consciousness | 94 |
| Seghezzi et al., 2019 | 31031676 | External Agency | 3 |
|  |  | Motor Intention | 5 |
|  |  | Sense of Agency | 4 |
| Servaas et al., 2013 | 23685122 | Neuroticism | 15 |
|  |  | Neuroticism (Decreases) | 48 |
| Sescousse et al., 2013 | 23415703 | Reward Outcome (Erotic) | 26 |
|  |  | Reward Outcome (Taste) | 28 |
|  |  | Reward Outcome (Money) | 24 |
| Shaw et al., 2012 | 21278194 | Action Hands & Faces | 20 |
| Shen et al., 2016 | 27720998 | Character Chunk Decomposition (Insight) | 12 |
|  |  | Compound Remote Associate (Insight) | 3 |
|  |  | Prototype Heuristic (Insight) | 6 |
| Shen et al., 2018 | 30165082 | Insight | 8 |
| Shkurko et al., 2013 | 22847948 | In-group (Social Categorization) | 7 |
|  |  | Out-group (Social Categorization) | 6 |
|  |  | Social Categorization | 12 |
| Silverman et al., 2015 | 26254587 | Reward Anticipation (Adolescence) | 5 |
|  |  | Reward Outcome (Adolescence) | 8 |
| SimanTov et al., 2019 | 31437478 | Prediction | 17 |
| Simmonds et al., 2009 | 17850833 | Complex Go No-Go | 10 |
|  |  | Simple Go No-Go | 4 |
| Skipper et al., 2014 | 25092665 | Non-Word Sounds | 10 |
|  |  | Speech Phonology | 10 |
| Sokolowski et al., 2016 | 27769786 | Non-Symbolic Numbers | 18 |
|  |  | Numbers | 13 |
|  |  | Passive Numbers | 9 |
|  |  | Symbolic Numbers | 12 |
| Sokolowski et al., 2017 | 28119003 | Non-Numerical | 12 |
|  |  | Non-Symbolic Numerical | 17 |
|  |  | Numerical | 22 |
|  |  | Symbolic Numerical | 15 |
| Song et al., 2017 | 28522823 | Stroop Task | 6 |
| Sorella et al., 2021 | 33503484 | Anger Experience | 5 |
|  |  | Anger Perception | 9 |
| Soros et al., 2008 | 19107749 | Saliva Swallowing | 11 |
|  |  | Water Swallowing | 13 |
| Spaniol et al., 2009 | 19428409 | Encoding Success | 21 |
|  |  | Objective Recollection | 17 |
|  |  | Retrieval Success | 19 |
|  |  | Subjective Recollection | 16 |
| Sperduti et al., 2011 | 21212978 | External Agency | 5 |
|  |  | Self-Agency | 11 |
| Sperduti et al., 2012 | 22005087 | Meditation | 3 |
| Spreng et al., 2009 | 18510452 | Autobiographical memory | 22 |
|  |  | Prospection Self | 12 |
|  |  | Spatial Navigation | 17 |
|  |  | Theory of Mind | 22 |
| Spreng et al., 2010 | 20109489 | Cognition | 32 |
| Stawarczyk et al., 2015 | 25931002 | Episodic Future Thinking | 16 |
|  |  | Mind Wandering | 7 |
|  |  | Personal Goals | 12 |
| Steele et al., 2004 | 15006653 | Cognitive Tasks | 2 |
|  |  | Emotion Induction | 2 |
| Stefaniak et al., 2021 | 33744459 | Comprehension Task | 15 |
|  |  | Language Network | 30 |
|  |  | Production Task | 23 |
| Stevens et al., 2012 | 22450197 | Negative Emotions (Men) | 12 |
|  |  | Negative Emotions (Women) | 7 |
|  |  | Positive Emotions (Men) | 10 |
|  |  | Positive Emotions (Women) | 1 |
| Stoléru et al., 2012 | 22465619 | Sexual Arousal (Heterosexuals) | 42 |
| Sulpizio et al., 2020 | 31838193 | Grammar (L1) | 9 |
|  |  | Grammar (L2) | 6 |
|  |  | Language Switching | 18 |
|  |  | Lexico-Semantics (L1) | 18 |
|  |  | Lexico-Semantics (L2) | 10 |
|  |  | Phonology (L1) | 3 |
|  |  | Phonology (L2) | 6 |
| Swick et al., 2011 | 21376819 | Go No-Go | 26 |
|  |  | Stop-Signal | 17 |
| Tagarelli et al., 2019 | 30826361 | Grammar Learning | 28 |
|  |  | Lexicon Learning | 26 |
| Tan et al., 2005 | 15846817 | Alphabetic Words | 11 |
|  |  | Chinese Characters | 10 |
| Tanasescu et al., 2016 | 27168346 | Cutaneous Experimental Pain | 48 |
| Tang et al., 2012 | 22450260 | Food | 16 |
| Tanzer et al., 2020 | 10.1007s10902-019-00195-7 | Engagement (Happiness) | 4 |
|  |  | Happiness | 11 |
|  |  | Meaning (Happiness) | 5 |
|  |  | Pleasure (Happiness) | 7 |
| Tao et al., 2021 | 33359340 | Explicit Fear Processing | 13 |
|  |  | Implicit Fear Processing | 13 |
|  |  | Language Switching | 29 |
|  |  | Task Switching | 7 |
| Taylor et al., 2013 | 23046391 | Pseudowords | 34 |
|  |  | Words | 23 |
| Teghil et al., 2019 | 30316722 | Externally-Cued Timing | 12 |
|  |  | Internally-based Timing | 11 |
|  |  | Timing | 19 |
| Thayer et al., 2012 | 22178086 | Heart Variability | 3 |
| Timmers et al., 2018 | 30542272 | Empathy | 37 |
|  |  | Empathy Pain | 27 |
|  |  | Empathy for Affective State | 30 |
| Tomasino et al., 2012 | 23316154 | Focused-Attention Meditation | 6 |
|  |  | Mantra-Induced Meditation | 4 |
|  |  | Meditation (Expert) | 7 |
|  |  | Meditation (Novice) | 5 |
|  |  | Meditation Network (Activation) | 11 |
|  |  | Meditation Network (Deactivation) | 9 |
| Tomasino et al., 2014 | 25446003 | Abstract Verbs | 11 |
| Tomasino et al., 2014 | 24975229 | Buddhism (Meditation) | 5 |
|  |  | Hinduism (Meditation) | 5 |
| Tomasino et al., 2016 | 26779003 | Mental Rotation | 24 |
|  |  | Mental Rotation (Bodily Stimuli) | 22 |
|  |  | Mental Rotation (Motor Strategy) | 15 |
|  |  | Mental Rotation (Non-Bodily Stimuli) | 13 |
|  |  | Mental Rotation (Visuospatial Strategy) | 16 |
| Trettenbrein et al., 2020 | 33118302 | Sign Language Comprehension | 13 |
|  |  | Sign-like Action Observation | 21 |
| Turesky et al., 2016 | 27799910 | Right Hand Motor (Old) | 14 |
|  |  | Right Hand Motor (Young) | 9 |
| Turkeltaub et al., 2010 | 20413149 | Phoneme Perception | 2 |
|  |  | Speech | 5 |
| Turkerltaub et al., 2002 | 12169260 | Single Word Reading | 12 |
| Turner et al., 2012 | 21791362 | Inhibition (Older Adults) | 11 |
|  |  | Inhibition (Younger Adults) | 14 |
|  |  | Working Memory (Older Adults) | 16 |
|  |  | Working Memory (Younger Adults) | 19 |
| Turner et al., 2015 | 26426534 | Conflict | 3 |
|  |  | Model Building | 7 |
|  |  | Rule Checking | 12 |
|  |  | Rule Finding | 13 |
|  |  | Set Shifting | 9 |
|  |  | Statement Verification | 9 |
| Vaccaro et al., 2018 | 30542659 | Meta-Cognition | 8 |
| VantHooft et al., 2021 | 33421942 | Empathy | 84 |
|  |  | Music Perception | 34 |
|  |  | Social Cognition | 59 |
|  |  | Theory of Mind | 50 |
| Vargas et al., 2016 | 30240034 | Heart Rate | 5 |
|  |  | Heart Rate (Negative corr) | 8 |
|  |  | Heart Rate Variability | 9 |
|  |  | Heart Rate Variability (Negative corr) | 3 |
| Vartanian et al., 2012 | 22804698 | Analogy | 2 |
|  |  | Analogy & Metaphor | 7 |
|  |  | Metaphor | 3 |
| Vartanian et al., 2014 | 24704947 | Painting Viewing | 17 |
| Veldhuizen et al., 2011 | 21305668 | Gustatory Stimulation | 15 |
| Verhagen et al., 2006 | 16457886 | Odor Taste | 23 |
| Vijayakumar et al., 2017 | 28235565 | Social Exclusion | 11 |
| Von Der Heide et al., 2013 | 23378834 | Familiar Faces Recognition | 15 |
|  |  | Famous Faces Recognition | 18 |
| Vytal-Hamann et al., 2010 | 19929758 | Anger | 13 |
|  |  | Disgust | 16 |
|  |  | Fear | 11 |
|  |  | Happiness | 8 |
|  |  | Sadness | 22 |
| Wager et al., 2003 | 12880784 | Approach Emotions | 4 |
|  |  | Withdraw Emotions | 14 |
| Wager et al., 2004 | 15275924 | Switching Density | 7 |
| Wager et al., 2003 | 15040547 | Executive Processing (Working Memory) | 30 |
|  |  | Passive Storage (Working Memory) | 22 |
| Wagner et al., 2014 | 24456150 | Phonematic Verbal Fluency | 22 |
|  |  | Semantic Verbal Fluency | 14 |
| Walenski et al., 2019 | 30689268 | Sentence Comprehension | 21 |
|  |  | Sentence Production | 9 |
| Wallis et al., 2015 | 26042457 | Precue Spatial Attention | 19 |
|  |  | Retrocue Spatial Attention | 21 |
| Wang et al., 2010 | 20108224 | Abstract Processing | 8 |
|  |  | Concrete Processing | 8 |
| Wang et al., 2019 | 30708115 | Effect of Memory Load (N-Back) | 5 |
|  |  | Effect of Object Identity (N-Back) | 3 |
|  |  | N-Back | 22 |
| Wang et al., 2019 | 31474844 | Prosocial Behaviors | 3 |
|  |  | Reward | 8 |
| Wang et al., 2020 | 31339998 | Reward Network | 14 |
| Watanuki et al., 2020 | 33100955 | Romantic Love | 33 |
| Watson et al., 2013 | 23574587 | Action Concepts | 4 |
|  |  | Action Images | 4 |
|  |  | Action Lexical-Semantic Concepts | 4 |
|  |  | Action Verbs | 4 |
| Wertheim et al., 2018 | 30024328 | Deductive Reasoning | 5 |
|  |  | Geometrical Form | 2 |
|  |  | Inductive Reasoning | 8 |
|  |  | Relational Reasoning | 8 |
|  |  | Semantic Reasoning | 2 |
|  |  | Symbolic Reasoning | 6 |
| Wesley et al., 2014 | 24041504 | Delay Discounting | 20 |
|  |  | Finger Tapping | 12 |
|  |  | Response Inhibition | 14 |
|  |  | Working Memory | 16 |
| Wiener et al., 2010 | 19800975 | Subseconds (Motor) | 24 |
|  |  | Subseconds (Perceptual) | 17 |
|  |  | Supraseconds (Motor) | 16 |
|  |  | Supraseconds (Perceptual) | 11 |
| Wiener et al., 2010 | 20863842 | Implicit Timing | 1 |
| Wilson et al., 2018 | 30255220 | Anticipation of Loss | 214 |
|  |  | Anticipation of Reward | 197 |
| Winlove et al., 2018 | 29502874 | Imagination of Common Concrete Object | 10 |
|  |  | Visual Imagery | 22 |
|  |  | Visual Imagery (Eyes Closed) | 14 |
|  |  | Visual Imagery (vs Active Baseline) | 15 |
| Witt et al., 2008 | 18511305 | Finger Tapping | 10 |
| Witteman et al., 2012 | 22841991 | Emotional Prosody (Stimulus Effect) | 24 |
|  |  | Emotional Prosody (Task Effect) | 6 |
| Worringer et al., 2019 | 31037397 | Dual-Tasking | 18 |
|  |  | Task Switching | 17 |
| Wu et al., 2012 | 22759996 | Orthography | 9 |
|  |  | Phonology | 24 |
|  |  | Semantic | 25 |
| Wu et al., 2015 | 25891081 | Divergent Thinking | 18 |
| Wu et al., 2016 | 27592151 | Agreement with Group | 3 |
|  |  | Conforming Behaviors | 4 |
|  |  | Disagreement with Group | 6 |
|  |  | Social Influence on Valuation | 3 |
|  |  | Unfairness (Ultimatum Game) | 15 |
| Wu et al., 2021 | 33940147 | Ambiguity Processing | 4 |
|  |  | Decision Risk | 6 |
|  |  | Gains Gambles (Risk) | 3 |
|  |  | Mix Gambles (Risk) | 3 |
|  |  | Risk (Binary Contrast) | 5 |
|  |  | Risk (Parametric) | 4 |
|  |  | Risk Processing | 5 |
| Wu et al., 2020 | 31674015 | Cognitive Flexibility | 10 |
|  |  | Conflict Processing | 6 |
|  |  | Inhibition | 19 |
|  |  | Planning | 7 |
|  |  | Uncertainty (Cognition) | 17 |
|  |  | Uncertainty (Decision-Making) | 6 |
|  |  | Uncertainty (Executive Control) | 19 |
|  |  | Uncertainty (Working Memory) | 14 |
|  |  | Verbal Working Memory | 14 |
|  |  | Visuospatial Working Memory | 12 |
| Xiong et al., 2019 | 30531091 | Facial Expression of Pain | 30 |
|  |  | Facial Pain Empathy | 30 |
| Xu et al., 2016 | 27895564 | Emotional Interference | 19 |
|  |  | Non-Emotional Spatial Interference | 22 |
|  |  | Non-Emotional Verbal Interference | 22 |
| Xu et al., 2020 | 31954149 | Chemical Pain | 10 |
|  |  | Distal Pain | 6 |
|  |  | Electrical Pain | 5 |
|  |  | Left Pain | 6 |
|  |  | Mechanical Pain | 7 |
|  |  | Nociceptive Pain | 4 |
|  |  | Non-Thermal Pain | 5 |
|  |  | Non-Visceral Pain | 10 |
|  |  | Pain (Female) | 5 |
|  |  | Pain (Male) | 8 |
|  |  | Proximal Pain | 6 |
|  |  | Right Pain | 9 |
|  |  | Thermal Pain | 5 |
|  |  | Visceral Pain | 5 |
| Yang et al., 2014 | 25450866 | Motor Task (Expert) | 22 |
|  |  | Motor Task (Novice) | 18 |
| Yang et al., 2015 | 25681159 | Anidiomatic Action | 3 |
|  |  | Fictive Motion | 7 |
|  |  | Metaphoric Action | 4 |
| Yang et al., 2017 | 28926795 | Nouns | 15 |
|  |  | Verbs | 19 |
| Yang et al., 2019 | 31593233 | Betrayal | 2 |
|  |  | Cooperation | 3 |
|  |  | Fairness | 4 |
|  |  | Non-Cooperation | 6 |
|  |  | Other Non-Cooperation | 7 |
|  |  | Other-Cooperation | 1 |
|  |  | Reciprocity | 3 |
|  |  | Self Non-Cooperation | 2 |
|  |  | Unfairness | 6 |
| Yaple et al., 2018 | 29732553 | N-Back (Children) | 13 |
| Yaple et al., 2019 | 31004627 | Prediction Errors | 6 |
|  |  | Reversal Errors | 9 |
|  |  | Reversal Switches | 6 |
| Yaple et al., 2019 | 30954708 | N-Back (Middle-aged Adults) | 21 |
|  |  | N-Back (Older Adults) | 20 |
|  |  | N-Back (Young Adults) | 28 |
| Yaple et al., 2020 | 33463879 | Monetary Downward Comparison | 4 |
|  |  | Monetary Upward Comparison | 5 |
|  |  | Social Downward Comparison | 1 |
|  |  | Social Upward Comparison | 3 |
| Yaple et al., 2020 | 32638450 | Reward Processing (Adolescents) | 15 |
|  |  | Reward Processing (Adults) | 17 |
|  |  | Reward Processing (Children) | 7 |
| Yeo et al., 2017 | 28467892 | Symbolic Numerical | 9 |
| Yeung et al., 2017 | 29250913 | Stimuli Mimicking Dental Treatment | 5 |
| Yeung et al., 2017 | 28413706 | Taste Stimulation | 13 |
| Yeung et al., 2018 | 29247808 | Affective Taste | 16 |
|  |  | Intensity Taste | 1 |
|  |  | Quality Taste | 1 |
| Yu et al., 2019 | 31251965 | Deceptive Responses | 24 |
|  |  | False Recognition | 9 |
| Yuan et al., 2015 | 26051526 | Drawing | 33 |
|  |  | Memory-based Production | 28 |
|  |  | Model-based Production | 36 |
|  |  | Writing | 24 |
| Yuan et al., 2019 | 31275121 | Large-scale Spatial Abilities (Men) | 28 |
|  |  | Large-scale Spatial Abilities (Women) | 8 |
|  |  | Small-scale Spatial Abilities (Men) | 31 |
|  |  | Small-scale Spatial Abilities (Women) | 17 |
| Yuki et al., 2019 | 31181421 | Metacognition about Memory | 6 |
| Zaccarella et al., 2017 | 28743620 | Content & Function of Word List | 10 |
|  |  | Licit Word List Structure | 11 |
| Zacks et al., 2008 | 17919082 | Mental Rotation | 10 |
| Zapparoli et al., 2017 | 28567010 | What? | 4 |
|  |  | When? | 8 |
|  |  | Whether | 6 |
| Zevin et al., 2010 | 20089919 | Sensitivity to Phonetic Change | 4 |
| Zhang et al., 2016 | 27833571 | Number Processing | 21 |
| Zhang et al., 2017 | 28551777 | Action Cancellation | 16 |
|  |  | Action Withholding | 21 |
|  |  | Interference Resolution | 8 |
| Zhang et al., 2021 | 33817920 | Executive Functions | 4 |
|  |  | Executive Functions (Adolescents) | 8 |
|  |  | Executive Functions (Adults) | 4 |
|  |  | Inhibition | 9 |
|  |  | Inhibition (Adolescents) | 5 |
|  |  | Inhibition (Adults) | 8 |
|  |  | NonVerbal Inhibition | 12 |
|  |  | NonVerbal Switching | 9 |
|  |  | NonVerbal Working Memory | 12 |
|  |  | Social Semantic Network | 6 |
|  |  | Switching | 9 |
|  |  | Switching (Adolescents) | 1 |
|  |  | Switching (Adults) | 9 |
|  |  | Verbal Inhibition | 7 |
|  |  | Verbal Switching | 5 |
|  |  | Verbal Working Memory | 13 |
|  |  | Working Memory | 12 |
|  |  | Working Memory (Adolescents) | 12 |
|  |  | Working Memory (Adults) | 12 |
| Zhao et al., 2017 | 10.1016j.jneuroling.2017.04.001 | Chinese Characters | 12 |
|  |  | Chinese Characters (Stimulus Effects) | 10 |
|  |  | Chinese Characters (Task Effects) | 10 |
| Zhou et al., 2017 | 28523225 | Prosaccades | 19 |
|  |  | Word Reading | 15 |
| Zinchenko et al., 2017 | 29160930 | Norm Violation | 11 |
|  |  | Social Norm Representation | 2 |
|  |  | Social Norms | 10 |
| Zinchenko et al., 2018 | 29922137 | Dynamic Faces | 15 |
| Zinchenko et al., 2019 | 31488865 | Social Punishment | 4 |
| Zito et al., 2020 | 32502189 | Negative Agency (Motor Control) | 3 |
| van Hoorn et al., 2019 | 31006540 | Social Decision Making Adolescent | 8 |
|  |  | Decision-Making (Adolescent) | 8 |
| van Meer et al., 2015 | 25285373 | Food (Adolescent) | 15 |
|  |  | Food (Adult) | 29 |
| van Veluw et al., 2013 | 24535033 | False Beliefs (ToM) | 10 |
|  |  | Self Face Recognition | 5 |
| van der Laan et al., 2011 | 21111829 | Food | 23 |
|  |  | High Energy Food | 6 |
|  |  | Hungry | 3 |
| vanderLaan et al., 2016 | 10.1016j.cofs.2014.11.001 | Eating Behaviors | 12 |
| vanderMeer et al., 2010 | 20015455 | Self Reflection | 9 |
| Note. | | | |

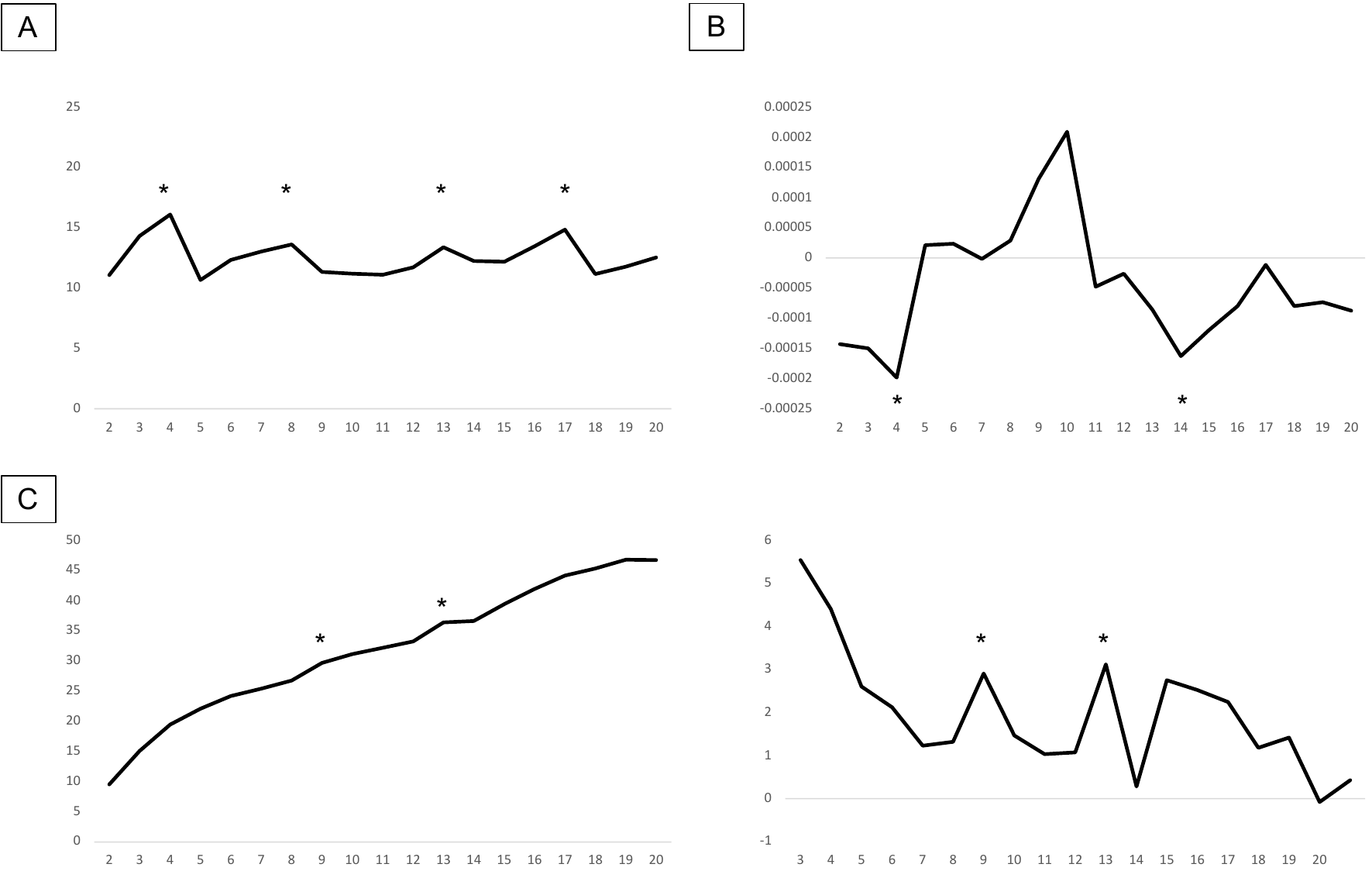

**Supplementary Figure 1.** Metrics for k=2 to k=20 solutions. **A.** The Silhouette coefficients showed that k=4, k=8, k=13, k=17 were the best solutions. **B.** Adjusted Rand Indices showed that the clustering solutions were the most highly discordant from random spatial arrangement at k=4 and k=14. Also, aRI values were negative at k=2, k=3, and k=11 to k=20, suggesting strong dissimilarity between these solutions and random arrangements. **C.** Calinski-Harabasz index exhibited a monotonic increase (Left) but k=13 & k=9 both showed greatest increase from previous (k-1) and following (k+1) solutions (Right).

| **Supplementary Table 2A.** Peak Coordinates of the **CN1 - Multi-Demand Network** (Cluster Peak) | | | | | |
| --- | --- | --- | --- | --- | --- |
| Regions | MNI Coordinates | | | Peak intensity  (Z-score) | Cluster size (mm^3^) |
|  | x | y | z |  |  |
| L_aINS | -32 | 22 | 0 | 22.43 | 29528 |
| R_aINS | 34 | 22 | -2 | 23.47 | 24096 |
| aMCC/pre-SMA | -2 | 16 | 48 | 18.60 | 18736 |
| R_pSMG | 40 | -44 | 44 | 9.71 | 10192 |
| L_pSMG | -42 | -44 | 46 | 8.73 | 6168 |
| R_Pallidum | 14 | 8 | 2 | 10.47 | 3000 |
| L_FFG | -42 | -64 | -12 | 9.64 | 2496 |
| L_THAL | -12 | -18 | 4 | 7.24 | 1824 |
| L_lPFC | -44 | 44 | 6 | 5.70 | 1400 |
| R_FEF | 30 | 0 | 56 | 5.69 | 1104 |
| L_PoCG | -60 | -22 | 30 | 6.89 | 1072 |
| L_IPL | -58 | -50 | 32 | 6.28 | 608 |
| R_FFG | 44 | -60 | -8 | 5.22 | 592 |
| L_dlPFC | -36 | 40 | 24 | 5.63 | 432 |
| Cerebellum VI | 32 | -60 | -30 | 5.36 | 432 |
| R_IPS | 34 | -70 | 36 | 5.56 | 360 |
| L_FEF | -26 | -2 | 52 | 5.55 | 336 |
| R_pMTG | 58 | -28 | -2 | 4.74 | 312 |
| R_aSMG | 62 | -22 | 36 | 5.55 | 272 |
| R_sLOC | 12 | -66 | 54 | 4.60 | 200 |
| L_MFG | -34 | 50 | 18 | 4.28 | 192 |
| R_pSTG | 62 | -18 | -2 | 4.71 | 104 |
| *Note.* | | | | | |

| **Supplementary Table 2B.** Peak Coordinates of the **CN1 - Multi-Demand Network** (with Subpeaks) | | | | |
| --- | --- | --- | --- | --- |
| Regions | MNI Coordinates | | | Peak intensity  (Z-score) |
|  | x | y | z |  |
| R_aINS | 34 | 22 | -2 | 23.47 |
| L_aINS | -32 | 22 | 0 | 22.43 |
| aMCC/pre-SMA | -2 | 16 | 48 | 18.60 |
| L_CAUD | -14 | 8 | 2 | 11.42 |
| L_MFG | -44 | 8 | 30 | 11.02 |
| R_Pallidum | 14 | 8 | 2 | 10.47 |
| R_dlPFC | 40 | 34 | 26 | 10.17 |
| R_pSMG | 40 | -44 | 44 | 9.71 |
| L_FFG | -42 | -64 | -12 | 9.64 |
| L_IFG | -50 | 10 | 6 | 9.44 |
| R_MFG | 46 | 10 | 30 | 9.22 |
| L_pSMG | -42 | -44 | 46 | 8.73 |
| R_IFG | 50 | 16 | 4 | 8.61 |
| R_pSMG | 50 | -44 | 40 | 8.56 |
| L_THAL | -12 | -18 | 4 | 7.24 |
| L_PoCG | -60 | -22 | 30 | 6.89 |
| L_IPL | -58 | -50 | 32 | 6.28 |
| R_THAL | 8 | -16 | 6 | 5.86 |
| R_pSMG | 60 | -40 | 14 | 5.80 |
| L_lPFC | -44 | 44 | 6 | 5.70 |
| R_FEF | 30 | 0 | 56 | 5.69 |
| L_dlPFC | -36 | 40 | 24 | 5.63 |
| R_IPS | 34 | -70 | 36 | 5.56 |
| L_FEF | -26 | -2 | 52 | 5.55 |
| R_aSMG | 62 | -22 | 36 | 5.55 |
| Cerebellum VI | 32 | -60 | -30 | 5.36 |
| R_FFG | 44 | -60 | -8 | 5.22 |
| R_pMTG | 58 | -28 | -2 | 4.74 |
| R_pSTG | 62 | -18 | -2 | 4.71 |
| R_sLOC | 12 | -66 | 54 | 4.60 |
| L_MFG | -34 | 50 | 18 | 4.28 |
| *Note.* | | | | |

| **Supplementary Table 3A.** Peak Coordinates of the **CN2 - Face Processing Network** | | | | | |
| --- | --- | --- | --- | --- | --- |
| Regions | MNI Coordinates | | | Peak intensity  (Z-score) | Cluster size (mm^3^) |
|  | x | y | z |  |  |
| R_FFG | 48 | -64 | -6 | 13.47 | 20752 |
| L_FFG | -46 | -70 | -4 | 14.70 | 15536 |
| R_PreCG | 48 | 8 | 28 | 11.57 | 13352 |
| L_PreCG | -50 | 6 | 28 | 9.62 | 12320 |
| L_AMY | -20 | -6 | -16 | 21.26 | 5880 |
| R_AMY | 22 | -4 | -16 | 18.41 | 4792 |
| aMCC/pre-SMA | -2 | 16 | 44 | 8.11 | 4296 |
| L_THAL | -12 | -12 | 8 | 8.94 | 2640 |
| L_IPS | -38 | -48 | 56 | 7.87 | 2560 |
| R_SPL | 32 | -54 | 56 | 7.29 | 1120 |
| R_aSMG | 62 | -20 | 34 | 7.88 | 1112 |
| Superior Colliculus | 0 | -30 | -6 | 6.29 | 824 |
| L_aSMG | -58 | -26 | 34 | 7.26 | 784 |
| R_PALL | 16 | 6 | 2 | 6.43 | 736 |
| L_MTG | -52 | -48 | 4 | 6.54 | 488 |
| HYP | -2 | -2 | -10 | 4.99 | 384 |
| L_CAUD | -12 | 10 | 0 | 5.15 | 368 |
| pgACC | 0 | 38 | 8 | 5.40 | 360 |
| L_lOFC | -26 | 32 | -16 | 5.17 | 304 |
| L_pHIPP | -20 | -30 | -2 | 5.06 | 264 |
| L_PALL | -18 | 2 | 2 | 4.56 | 208 |
| L_IFG | -48 | 10 | 4 | 4.62 | 184 |
| R_PUT | 20 | -4 | 14 | 4.48 | 160 |
| *Note.* | | | | | |

| **Supplementary Table 3B.** Peak Coordinates of the **CN2 - Face Processing Network** (with Subpeaks) | | | | |
| --- | --- | --- | --- | --- |
| Regions | MNI Coordinates | | | Peak intensity  (Z-score) |
|  | x | y | z |  |
| R_FFG | 48 | -64 | -6 | 13.47 |
| L_FFG | -46 | -70 | -4 | 14.70 |
| L_AMY | -20 | -6 | -16 | 21.26 |
| R_AMY | 22 | -4 | -16 | 18.41 |
| L_FFG | -42 | -52 | -18 | 14.08 |
| R_FFG | 42 | -50 | -18 | 12.29 |
| R_PreCG | 48 | 8 | 28 | 11.57 |
| L_PreCG | -50 | 6 | 28 | 9.62 |
| L_THAL | -12 | -12 | 8 | 8.94 |
| L_aINS | -32 | 22 | 2 | 8.73 |
| L_mINS | -40 | -2 | 0 | 8.48 |
| L_OccipitalPole | -30 | -92 | 0 | 8.21 |
| aMCC/pre-SMA | -2 | 16 | 44 | 8.11 |
| R_aSMG | 62 | -20 | 34 | 7.88 |
| L_IPS | -38 | -48 | 56 | 7.87 |
| R_OccipitalPole | 18 | -92 | -2 | 7.57 |
| R_IFG | 50 | 30 | -2 | 7.53 |
| R_SPL | 32 | -54 | 56 | 7.29 |
| L_aSMG | -58 | -26 | 34 | 7.26 |
| R_aINS | 38 | 20 | -2 | 7.17 |
| R_mINS | 42 | 4 | 0 | 6.69 |
| L_MTG | -52 | -48 | 4 | 6.54 |
| R_PALL | 16 | 6 | 2 | 6.43 |
| Superior Colliculus | 0 | -30 | -6 | 6.29 |
| L_V3 | -20 | -96 | -6 | 6.07 |
| L_PoCG | -40 | -38 | 44 | 5.69 |
| dACC | 0 | 20 | 26 | 5.58 |
| pSTG | 50 | -34 | 2 | 5.57 |
| pgACC | 0 | 38 | 8 | 5.40 |
| lOFC | -26 | 32 | -16 | 5.17 |
| L_CAUD | -12 | 10 | 0 | 5.15 |
| L_pHIPP | -20 | -30 | -2 | 5.06 |
| HYP | -2 | -2 | -10 | 4.99 |
| L_IFG | -48 | 10 | 4 | 4.62 |
| L_PALL | -18 | 2 | 2 | 4.56 |
| R_PUT | 20 | -4 | 14 | 4.48 |
| *Note.* | | | | |

| **Supplementary Table 4A.** Peak Coordinates of the **CN3 - Language Network** | | | | | |
| --- | --- | --- | --- | --- | --- |
| Regions | MNI Coordinates | | | Peak intensity  (Z-score) | Cluster size  (mm^3^) |
|  | x | y | z |  |  |
| L_IFG | -46 | 26 | -6 | 12.06 | 28208 |
| L_pMTG | -58 | -38 | 0 | 12.30 | 21152 |
| SMA | -4 | 12 | 60 | 12.41 | 12144 |
| R_IFG | 40 | 22 | -6 | 9.00 | 8120 |
| pSTG | 62 | -14 | -2 | 6.03 | 3104 |
| R_MFG | 40 | 22 | 42 | 7.80 | 744 |
| R_AG | 58 | -54 | 38 | 6.54 | 616 |
| L_TP | -52 | 10 | -20 | 5.95 | 488 |
| L_lPFC | -44 | 46 | -6 | 5.32 | 368 |
| L_SPL | -36 | -48 | 46 | 4.44 | 360 |
| R_lPFC | 42 | 48 | -4 | 5.15 | 352 |
| L_AG | -52 | -58 | 44 | 5.17 | 280 |
| FFG | -36 | -36 | -20 | 4.38 | 208 |
| R_IPL | 50 | -60 | 42 | 4.88 | 136 |
| L_MFG | -32 | 48 | 16 | 4.30 | 88 |
| *Note.* | | | | | |

| **Supplementary Table 4B.** Peak Coordinates of the **CN3 - Language Network** (with Subpeaks) | | | | |
| --- | --- | --- | --- | --- |
| Regions | MNI Coordinates | | | Peak intensity (Z-score) |
|  | x | y | z |  |
| SMA | -4 | 12 | 60 | 12.41 |
| L_pMTG | -58 | -38 | 0 | 12.30 |
| L_IFG | -46 | 26 | -6 | 12.06 |
| L_dlPFC | -48 | 22 | 24 | 11.37 |
| L_MFG | -42 | 4 | 48 | 10.55 |
| L_dlPFC | -44 | 10 | 28 | 9.99 |
| R_IFG | 40 | 22 | -6 | 9.00 |
| R_IFG | 50 | 30 | -8 | 8.91 |
| L_IPS | -28 | -64 | 46 | 8.43 |
| R_MFG | 40 | 22 | 42 | 7.80 |
| R_dlPFC | 50 | 20 | 24 | 7.41 |
| L_FFG | -46 | -60 | -12 | 7.04 |
| R_AG | 58 | -54 | 38 | 6.54 |
| L_STG | -60 | -12 | -4 | 6.43 |
| L_IPL | -46 | -54 | 24 | 6.17 |
| pSTG | 62 | -14 | -2 | 6.03 |
| L_TP | -52 | 10 | -20 | 5.95 |
| R_pSTG | 52 | -34 | 2 | 5.9 |
| L_sLOC | -46 | -66 | 36 | 5.56 |
| L_lPFC | -44 | 46 | -6 | 5.32 |
| L_AG | -52 | -58 | 44 | 5.17 |
| R_lPFC | 42 | 48 | -4 | 5.15 |
| R_IPL | 50 | -60 | 42 | 4.88 |
| L_SPL | -36 | -48 | 46 | 4.44 |
| L_FFG | -36 | -36 | -20 | 4.38 |
| L_MFG | -32 | 48 | 16 | 4.30 |
| *Note.* | | | | |

| **Supplementary Table 5A.** Peak Coordinates of the **CN4 - Spatial Memory Network** | | | | | |
| --- | --- | --- | --- | --- | --- |
| Regions | MNI Coordinates | | | Peak intensity  (Z-score) | Cluster size (mm^3^) |
|  | x | y | z |  |  |
| L_LG | -26 | -48 | -8 | 8.34 | 3904 |
| R_LG | 24 | -40 | -8 | 12.05 | 3568 |
| R_RSPL | 16 | -54 | 14 | 10.02 | 2648 |
| L_RSPL | -16 | -58 | 18 | 8.20 | 2064 |
| aMCC/pre-SMA | 4 | 16 | 48 | 7.01 | 1456 |
| L_FEF | -28 | -2 | 56 | 7.52 | 984 |
| R_sLOC | 34 | -76 | 26 | 4.77 | 864 |
| R_aINS | 32 | 24 | -4 | 7.36 | 840 |
| R_FEF | 26 | 4 | 56 | 6.82 | 792 |
| L_sLOC | -32 | -84 | 24 | 7.09 | 720 |
| R_SPL | 18 | -64 | 60 | 5.91 | 536 |
| L_V2 | -6 | -80 | -2 | 5.69 | 464 |
| MCC/SMA | -6 | 2 | 54 | 4.63 | 400 |
| PCUN | 14 | -74 | 40 | 4.72 | 384 |
| R_IPL | 44 | -76 | 32 | 4.95 | 272 |
| R_iLOC | 46 | -72 | -2 | 4.63 | 160 |
| L_V3 | -20 | -68 | -2 | 4.74 | 152 |
| L_SPL | -20 | -64 | 60 | 4.29 | 104 |
| L_iLOC | -44 | -76 | 0 | 4.32 | 104 |
| *Note.* | | | | | |

| **Supplementary Table 5B.** Peak Coordinates of the **CN4 - Spatial Memory Network** (with Subpeaks) | | | | |
| --- | --- | --- | --- | --- |
| Regions | MNI Coordinates | | | Peak intensity (Z-score) |
|  | x | y | z |  |
| L_LG | -26 | -48 | -8 | 8.34 |
| R_LG | 24 | -40 | -8 | 12.05 |
| R_vPCC | 16 | -54 | 14 | 10.02 |
| L_vPCC | -16 | -58 | 18 | 8.20 |
| R_RSPL | 10 | -46 | 2 | 8.14 |
| L_pHIPP | -22 | -38 | -6 | 7.57 |
| L_FEF | -28 | -2 | 56 | 7.52 |
| R_aINS | 32 | 24 | -4 | 7.36 |
| L_sLOC | -32 | -84 | 24 | 7.09 |
| aMCC/pre-SMA | 4 | 16 | 48 | 7.01 |
| R_FEF | 26 | 4 | 56 | 6.82 |
| L_RSPL | -10 | -52 | 6 | 6.32 |
| R_SPL | 18 | -64 | 60 | 5.91 |
| L_V2 | -6 | -80 | -2 | 5.69 |
| R_IPL | 44 | -76 | 32 | 4.95 |
| R_sLOC | 34 | -76 | 26 | 4.77 |
| L_V3 | -20 | -68 | -2 | 4.74 |
| PCUN | 14 | -74 | 40 | 4.72 |
| MCC/SMA | -6 | 2 | 54 | 4.63 |
| R_iLOC | 46 | -72 | -2 | 4.63 |
| L_iLOC | -44 | -76 | 0 | 4.32 |
| L_SPL | -20 | -64 | 60 | 4.29 |
| *Note.* | | | | |

| **Supplementary Table 6A.** Peak Coordinates of the **CN5 - Cognitive Control Network** | | | | | |
| --- | --- | --- | --- | --- | --- |
| Regions | MNI Coordinates | | | Peak intensity  (Z-score) | Cluster size (mm^3^) |
|  | x | y | z |  |  |
| R_pSMG | 42 | -44 | 46 | 19.49 | 40856 |
| L_dlPFC | -44 | 8 | 28 | 24.19 | 21072 |
| R_dlPFC | 48 | 10 | 28 | 21.44 | 19592 |
| aMCC/pre-SMA | 0 | 16 | 48 | 25.28 | 12432 |
| L_ITG | -46 | -60 | -12 | 12.24 | 8176 |
| R_aINS | 34 | 22 | -2 | 26.07 | 6176 |
| R_Crus1 | 32 | -62 | -30 | 7.46 | 5872 |
| L_aINS | -32 | 22 | 0 | 22.00 | 4432 |
| L_CAUD | -16 | 0 | 14 | 10.11 | 3128 |
| R_CAUD | 14 | 8 | 4 | 7.60 | 1608 |
| R_THAL | 10 | -12 | 4 | 6.10 | 1320 |
| L_lPFC | -38 | 52 | 12 | 9.40 | 1224 |
| L_Crus1 | -32 | -62 | -34 | 6.10 | 544 |
| L_Thal | -14 | -16 | 10 | 5.75 | 520 |
| L_aSMG | -56 | -26 | 38 | 5.99 | 432 |
| R_iLOC | 48 | -66 | 0 | 4.80 | 216 |
| R_IPL | 60 | -36 | 20 | 4.58 | 216 |
| R_sLOC | 36 | -84 | 16 | 3.99 | 64 |
| *Note.* | | | | | |

| **Supplementary Table 6B.** Peak Coordinates of the **CN5 - Cognitive Control Network** (with Subpeaks) | | | | |
| --- | --- | --- | --- | --- |
| Regions | MNI Coordinates | | | Peak intensity (Z-score) |
|  | x | y | z |  |
| R_aINS | 34 | 22 | -2 | 26.07 |
| aMCC/pre-SMA | 0 | 16 | 48 | 25.28 |
| L_dlPFC | -44 | 8 | 28 | 24.19 |
| L_aINS | -32 | 22 | 0 | 22.00 |
| R_dlPFC | 48 | 10 | 28 | 21.44 |
| L_FEF | -26 | -2 | 54 | 20.65 |
| R_pSMG | 42 | -44 | 46 | 19.49 |
| L_SPL | -36 | -48 | 44 | 18.45 |
| R_FEF | 30 | 0 | 56 | 16.76 |
| L_IPS | -26 | -62 | 48 | 14.91 |
| R_IPS | 34 | -58 | 48 | 14.84 |
| R_dlPFC | 44 | 38 | 24 | 13.64 |
| L_ITG | -46 | -60 | -12 | 12.24 |
| R_SPL | 18 | -64 | 56 | 10.95 |
| L_iLOC | -44 | -70 | -6 | 10.77 |
| L_CAUD | -16 | 0 | 14 | 10.11 |
| L_lPFC | -38 | 52 | 12 | 9.40 |
| L_SPL | -12 | -68 | 54 | 8.98 |
| R_CAUD | 14 | 8 | 4 | 7.60 |
| R_Crus1 | 32 | -62 | -30 | 7.46 |
| R_ITG | 50 | -56 | -12 | 7.03 |
| R_iLOC | 36 | -86 | 0 | 6.23 |
| R_THAL | 10 | -12 | 4 | 6.10 |
| L_Crus1 | -32 | -62 | -34 | 6.10 |
| L_aSMG | -56 | -26 | 38 | 5.99 |
| L_Thal | -14 | -16 | 10 | 5.75 |
| R_iLOC | 48 | -66 | 0 | 4.80 |
| R_IPL | 60 | -36 | 20 | 4.58 |
| VTA | 2 | -20 | -4 | 4.47 |
| R_sLOC | 36 | -84 | 16 | 3.99 |
| *Note.* | | | | |

| **Supplementary Table 7A.** Peak Coordinates of the **CN6 - Physiological Arousal Network** | | | | | |
| --- | --- | --- | --- | --- | --- |
| Regions | MNI Coordinates | | | Peak intensity  (Z-score) | Cluster size  (mm^3^) |
|  | x | y | z |  |  |
| R_aINS | 40 | 18 | 0 | 14.35 | 27392 |
| L_aINS | -34 | 16 | 2 | 12.76 | 21368 |
| dACC | 2 | 14 | 36 | 10.41 | 8656 |
| R_THAL | 12 | -16 | 6 | 12.91 | 8088 |
| R_lPFC | 38 | 48 | 14 | 6.75 | 3224 |
| R_PALL | 12 | 6 | -4 | 5.28 | 1408 |
| L_AMY | -22 | -4 | -14 | 5.49 | 680 |
| R_PUT | -20 | 6 | 4 | 5.20 | 640 |
| R_AMY | 24 | 0 | -12 | 4.71 | 568 |
| L_Crus1 | -36 | -62 | -32 | 4.61 | 384 |
| R_CerebVI | 26 | -64 | -24 | 4.61 | 304 |
| SMA | 8 | 10 | 62 | 5.38 | 280 |
| L_PreCG | -56 | 4 | 24 | 4.76 | 256 |
| L_CerebVI | -22 | -66 | -26 | 4.61 | 200 |
| R_CerebV | 2 | -64 | -14 | 4.44 | 144 |
| *Note.* | | | | | |

| **Supplementary Table 7B.** Peak Coordinates of the **CN6 - Physiological Arousal Network** (with Subpeaks) | | | | |
| --- | --- | --- | --- | --- |
| Regions | MNI Coordinates | | | Peak intensity (Z-score) |
|  | x | y | z |  |
| R_aINS | 40 | 18 | 0 | 14.35 |
| R_THAL | 12 | -16 | 6 | 12.91 |
| L_aINS | -34 | 16 | 2 | 12.76 |
| L_THAL | -12 | -16 | 6 | 11.99 |
| R_SII | 56 | -24 | 22 | 11.88 |
| L_SII | -56 | -24 | 18 | 10.78 |
| dACC | 2 | 14 | 36 | 10.41 |
| L_pdINS | -40 | 0 | 8 | 9.64 |
| R_PreCG | 54 | 4 | 16 | 9.13 |
| R_pdINS | 40 | -6 | 12 | 8.27 |
| R_CO | 42 | -16 | 16 | 8.27 |
| L_CO | -40 | -20 | 14 | 7.79 |
| L_PreCG | -54 | 4 | 6 | 7.68 |
| MCC | 0 | 2 | 46 | 7.14 |
| R_lPFC | 38 | 48 | 14 | 6.75 |
| R_MFG | 42 | 36 | 28 | 5.94 |
| L_AMY | -22 | -4 | -14 | 5.49 |
| SMA | 8 | 10 | 62 | 5.38 |
| R_PALL | 12 | 6 | -4 | 5.28 |
| R_PUT | -20 | 6 | 4 | 5.20 |
| R_pSMG | 50 | -38 | 44 | 4.88 |
| L_PreCG | -56 | 4 | 24 | 4.76 |
| R_AMY | 24 | 0 | -12 | 4.71 |
| L_CerebVI | -22 | -66 | -26 | 4.61 |
| L_Crus1 | -36 | -62 | -32 | 4.61 |
| R_CerebVI | 26 | -64 | -24 | 4.61 |
| R_CerebV | 2 | -64 | -14 | 4.44 |
| *Note.* | | | | |

| **Supplementary Table 8A.** Peak Coordinates of the **CN7 - Auditory Perception Network** | | | | | |
| --- | --- | --- | --- | --- | --- |
| Regions | MNI Coordinates | | | Peak intensity  (Z-score) | Cluster size (mm^3^) |
|  | x | y | z |  |  |
| L_Planum | -58 | -16 | 0 | 7.73 | 8272 |
| R_Planum | 60 | -18 | 2 | 7.81 | 3832 |
| R_TP | 52 | 10 | -18 | 5.04 | 1064 |
| R_PreCG | 54 | 0 | 46 | 5.57 | 648 |
| L_Crus1 | -34 | -56 | -34 | 5.59 | 376 |
| dACC | 6 | 12 | 38 | 5.77 | 296 |
| R_lPFC | 44 | 44 | 6 | 4.94 | 160 |
| R_PO | 56 | -26 | 22 | 4.33 | 128 |
| R_pSMG | 54 | -42 | 50 | 4.28 | 88 |
| R_THAL | 14 | -14 | 8 | 4.22 | 80 |
| *Note.* | | | | | |

| **Supplementary Table 8B.** Peak Coordinates of the **CN7 - Auditory Perception Network** (with Subpeaks) | | | | |
| --- | --- | --- | --- | --- |
| Regions | MNI Coordinates | | | Peak intensity (Z-score) |
|  | x | y | z |  |
| R_Planum | 60 | -18 | 2 | 7.81 |
| L_Planum | -58 | -16 | 0 | 7.73 |
| R_pSTG | 58 | -28 | 6 | 7.44 |
| L_pSMG | -54 | -50 | 12 | 5.86 |
| dACC | 6 | 12 | 38 | 5.77 |
| L_Crus1 | -34 | -56 | -34 | 5.59 |
| R_PreCG | 54 | 0 | 46 | 5.57 |
| L_SII | -58 | -24 | 20 | 5.31 |
| L_Heschl | -42 | -28 | 10 | 5.11 |
| R_TP | 52 | 10 | -18 | 5.04 |
| R_lPFC | 44 | 44 | 6 | 4.94 |
| L_PO | -50 | -36 | 28 | 4.44 |
| R_PO | 56 | -26 | 22 | 4.33 |
| R_pSMG | 54 | -42 | 50 | 4.28 |
| R_THAL | 14 | -14 | 8 | 4.22 |
| *Note.* | | | | |

| **Supplementary Table 9A.** Peak Coordinates of the **CN8 - Social Inference Network** | | | | | |
| --- | --- | --- | --- | --- | --- |
| Regions | MNI Coordinates | | | Peak intensity  (Z-score) | Cluster size (mm^3^) |
|  | x | y | z |  |  |
| mPFC | 2 | 58 | 14 | 12.25 | 17024 |
| PCC/PCUN | -2 | -56 | 30 | 18.14 | 10696 |
| L_pSTS | -50 | -60 | 24 | 18.31 | 7816 |
| R_pSTS | 54 | -56 | 24 | 14.37 | 7488 |
| L_aMTG | -58 | -10 | -16 | 11.40 | 6936 |
| R_aMTG | 56 | -4 | -20 | 9.68 | 6112 |
| L_lOFC | -48 | 28 | -8 | 8.99 | 3832 |
| L_AMY | -20 | -4 | -18 | 6.85 | 1072 |
| L_HIPP | -24 | -24 | -18 | 5.47 | 744 |
| L_dmPFC | -10 | 34 | 52 | 4.63 | 440 |
| L_MFG | -44 | 8 | 46 | 5.56 | 432 |
| L_IFG | -52 | 24 | 14 | 5.08 | 416 |
| R_lOFC | 52 | 30 | -6 | 4.61 | 320 |
| R_pPHG | 24 | -34 | -18 | 5.39 | 312 |
| pMCC | -2 | -28 | 34 | 4.75 | 184 |
| aMCC/preSMA | -2 | 20 | 48 | 4.70 | 160 |
| R_HIPP | 26 | -14 | -20 | 4.35 | 120 |
| L_AG | 58 | -52 | 40 | 4.57 | 120 |
| *Note.* | | | | | |

| **Supplementary Table 9B.** Peak Coordinates of the **CN8 - Social Inference Network** (with Subpeaks) | | | | |
| --- | --- | --- | --- | --- |
| Regions | MNI Coordinates | | | Peak intensity  (Z-score) |
|  | x | y | z |  |
| mPFC | 2 | 58 | 14 | 12.25 |
| PCC/PCUN | -2 | -56 | 30 | 18.14 |
| L_pSTS | -50 | -60 | 24 | 18.31 |
| R_pSTS | 54 | -56 | 24 | 14.37 |
| L_aMTG | -58 | -10 | -16 | 11.40 |
| vmPFC | -2 | 52 | -6 | 11.39 |
| dmPFC | 0 | 56 | 28 | 10.79 |
| R_aMTG | 56 | -4 | -20 | 9.68 |
| L_lOFC | -48 | 28 | -8 | 8.99 |
| R_iLOC | 50 | -68 | 10 | 7.10 |
| L_pMTG | -58 | -26 | -8 | 6.90 |
| L_AMY | -20 | -4 | -18 | 6.85 |
| L_MFG | -44 | 8 | 46 | 5.56 |
| L_HIPP | -24 | -24 | -18 | 5.47 |
| R_pPHG | 24 | -34 | -18 | 5.39 |
| L_IFG | -52 | 24 | 14 | 5.08 |
| pMCC | -2 | -28 | 34 | 4.75 |
| aMCC/preSMA | -2 | 20 | 48 | 4.70 |
| L_dmPFC | -10 | 34 | 52 | 4.63 |
| R_lOFC | 52 | 30 | -6 | 4.61 |
| L_AG | 58 | -52 | 40 | 4.57 |
| R_HIPP | 26 | -14 | -20 | 4.35 |
| *Note.* | | | | |

| **Supplementary Table 10.** Peak Coordinates of the **CN9 - Spatial Attention Network** | | | | | |
| --- | --- | --- | --- | --- | --- |
| Regions | MNI Coordinates | | | Peak intensity  (Z-score) | Cluster size (mm^3^) |
|  | x | y | z |  |  |
| R_sLOC | 34 | -72 | 34 | 5.73 | 840 |
| R_PCUN | 10 | -66 | 34 | 5.65 | 648 |
| R_SPL | 18 | -74 | 52 | 6.31 | 504 |
| R_iLOC | 48 | -64 | -8 | 5.01 | 488 |
| R_FEF | 28 | -2 | 50 | 5.07 | 392 |
| L_sLOC | -26 | -80 | 32 | 4.98 | 352 |
| R_AG | 48 | -56 | 40 | 4.75 | 304 |
| R_sLOC2 | 38 | -80 | 24 | 5.16 | 280 |
| R_PCUN2 | 6 | -54 | 42 | 4.61 | 192 |
| R_lPFC | 36 | 50 | 4 | 4.11 | 120 |
| R_LOC | 32 | -82 | 12 | 4.39 | 96 |
| pMCC | -4 | -26 | 30 | 4.23 | 88 |
| *Note.* | | | | | |

| **Supplementary Table 11A.** Peak Coordinates of the **CN10 - Action Network** | | | | | |
| --- | --- | --- | --- | --- | --- |
| Regions | MNI Coordinates | | | Peak intensity  (Z-score) | Cluster size (mm^3^) |
|  | x | y | z |  |  |
| SMA | -2 | 2 | 56 | 13.00 | 29712 |
| R_SPL | 12 | -64 | 56 | 6.60 | 5256 |
| R_FEF | 28 | 4 | 54 | 8.99 | 3960 |
| R_CerebVI | 18 | -54 | -22 | 11.05 | 3880 |
| L_dlPFC | -54 | 6 | 28 | 8.48 | 2880 |
| L_PUT | -26 | -6 | 0 | 8.40 | 2824 |
| R_dlPFC | 58 | 10 | 16 | 6.94 | 2632 |
| R_PUT | 24 | 8 | 6 | 6.60 | 2312 |
| L_THAL | -14 | -20 | 6 | 11.43 | 1880 |
| L_SII | -50 | -24 | 18 | 6.69 | 1336 |
| R_PreCG | 40 | -18 | 58 | 5.99 | 952 |
| R_aSMG | 56 | -24 | 40 | 5.15 | 824 |
| R_THAL | 14 | -18 | 8 | 6.21 | 752 |
| L_CerebVI | -22 | -56 | -24 | 5.54 | 720 |
| L_iLOC | -48 | -66 | -6 | 5.27 | 304 |
| R_PoCG | 60 | -16 | 28 | 5.00 | 240 |
| L_CO | -44 | 0 | 10 | 4.65 | 216 |
| Cereb_I_IV | 2 | -54 | -12 | 4.43 | 152 |
| L_MFG | -36 | 40 | 30 | 4.55 | 136 |
| *Note.* | | | | | |

| **Supplementary Table 11B.** Peak Coordinates of the **CN10 - Action Network** (with Subpeaks) | | | | |
| --- | --- | --- | --- | --- |
| Regions | MNI Coordinates | | | Peak intensity  (Z-score) |
|  | x | y | z |  |
| SMA | -2 | 2 | 56 | 13.00 |
| L_PreCG | -38 | -22 | 58 | 12.88 |
| L_FEF | -24 | -4 | 56 | 11.45 |
| L_THAL | -14 | -20 | 6 | 11.43 |
| R_CerebVI | 18 | -54 | -22 | 11.05 |
| R_FEF | 28 | 4 | 54 | 8.99 |
| L_dlPFC | -54 | 6 | 28 | 8.48 |
| L_PUT | -26 | -6 | 0 | 8.40 |
| L_IPS | -24 | -60 | 54 | 7.34 |
| R_dlPFC | 58 | 10 | 16 | 6.94 |
| L_SII | -50 | -24 | 18 | 6.69 |
| R_PUT | 24 | 8 | 6 | 6.60 |
| R_SPL | 12 | -64 | 56 | 6.60 |
| R_SPL | 40 | -42 | 52 | 6.37 |
| R_THAL | 14 | -18 | 8 | 6.21 |
| R_SPL | 32 | -52 | 48 | 6.05 |
| R_PreCG | 40 | -18 | 58 | 5.99 |
| L_CerebVI | -22 | -56 | -24 | 5.54 |
| L_PoCG | -36 | -36 | 56 | 5.34 |
| L_iLOC | -48 | -66 | -6 | 5.27 |
| R_aSMG | 56 | -24 | 40 | 5.15 |
| R_PoCG | 60 | -16 | 28 | 5.00 |
| L_CO | -44 | 0 | 10 | 4.65 |
| L_MFG | -36 | 40 | 30 | 4.55 |
| Cereb_I_IV | 2 | -54 | -12 | 4.43 |
| *Note.* | | | | |

| **Supplementary Table 12A.** Peak Coordinates of the **CN11 - Valuation Network** | | | | | |
| --- | --- | --- | --- | --- | --- |
| Regions | MNI Coordinates | | | Peak intensity  (Z-score) | Cluster size  (mm^3^) |
|  | x | y | z |  |  |
| mPFC/pgACC | 8 | 50 | 6 | 8.22 | 15736 |
| vPCC | -8 | -56 | 14 | 7.20 | 1080 |
| pMCC | -2 | -22 | 38 | 7.25 | 864 |
| dmPFC | 2 | 48 | 38 | 6.70 | 624 |
| R_AG | 58 | -46 | 32 | 5.18 | 280 |
| L_dlPFC | -38 | 36 | 34 | 4.72 | 192 |
| L_NACC | -10 | 8 | -6 | 4.67 | 152 |
| R_aSMG | 58 | -34 | 36 | 4.56 | 120 |
| R_PreCG | 44 | 6 | 30 | 4.30 | 120 |
| dmPFC | 6 | 56 | 24 | 4.24 | 120 |
| *Note.* | | | | | |

| **Supplementary Table 12B.** Peak Coordinates of the **CN11 - Valuation Network** (with Subpeaks) | | | | |
| --- | --- | --- | --- | --- |
| Regions | MNI Coordinates | | | Peak intensity  (Z-score) |
|  | x | y | z |  |
| mPFC/pgACC | 8 | 50 | 6 | 8.22 |
| vmPFC | -2 | 50 | -10 | 8.08 |
| pMCC | -2 | -22 | 38 | 7.25 |
| vPCC | -8 | -56 | 14 | 7.20 |
| dmPFC | 2 | 48 | 38 | 6.70 |
| pgACC | -2 | 40 | 2 | 6.24 |
| L_pgACC | -12 | 44 | 14 | 6.04 |
| dACC | 0 | 44 | 20 | 5.44 |
| R_AG | 58 | -46 | 32 | 5.18 |
| L_dlPFC | -38 | 36 | 34 | 4.72 |
| L_NACC | -10 | 8 | -6 | 4.67 |
| R_aSMG | 58 | -34 | 36 | 4.56 |
| R_PreCG | 44 | 6 | 30 | 4.30 |
| dmPFC2 | 6 | 56 | 24 | 4.24 |
| *Note.* | | | | |

| **Supplementary Table 13A.** Peak Coordinates of the **CN12 - Motivation Network** | | | | | |
| --- | --- | --- | --- | --- | --- |
| Regions | MNI Coordinates | | | Peak intensity  (Z-score) | Cluster size (mm^3^) |
|  | x | y | z |  |  |
| R_NACC | 12 | 10 | -6 | 20.52 | 63144 |
| vmPFC | 0 | 46 | -10 | 9.77 | 21704 |
| pMCC | -2 | -32 | 30 | 9.57 | 1656 |
| PCC | -2 | -50 | 24 | 4.68 | 344 |
| R_dlPFC | 54 | 14 | 18 | 4.95 | 200 |
| L_iLOC | -34 | -84 | -8 | 5.24 | 200 |
| L_MFG | -24 | 32 | 48 | 4.89 | 168 |
| L_dlPFC | -46 | 6 | 30 | 4.55 | 168 |
| L_mINS | -40 | -2 | 0 | 4.39 | 152 |
| *Note.* | | | | | |

| **Supplementary Table 13B.** Peak Coordinates of the **CN12 - Motivation Network** (with Subpeaks) | | | | |
| --- | --- | --- | --- | --- |
| Regions | MNI Coordinates | | | Peak intensity  (Z-score) |
|  | x | y | z |  |
| R_NACC | 12 | 10 | -6 | 20.52 |
| L_NACC | -10 | 8 | -6 | 20.47 |
| R_AMY | 24 | -2 | -16 | 15.18 |
| R_aINS | 36 | 22 | -6 | 15.10 |
| L_AMY | -20 | -4 | -16 | 13.13 |
| L_aINS | -32 | 22 | -4 | 13.05 |
| vmPFC | 0 | 46 | -10 | 9.77 |
| pMCC | -2 | -32 | 30 | 9.57 |
| aMCC/pre-SMA | 2 | 22 | 40 | 9.53 |
| R_THAL | 2 | -14 | 8 | 9.09 |
| L_VTA | -4 | -20 | -12 | 9.01 |
| MCC/SMA | 2 | 10 | 48 | 8.85 |
| pgACC | 0 | 40 | 10 | 8.10 |
| R_VTA | 6 | -18 | -10 | 7.76 |
| vmPFC | -2 | 56 | -6 | 7.48 |
| L_iLOC | -34 | -84 | -8 | 5.24 |
| R_dlPFC | 54 | 14 | 18 | 4.95 |
| L_MFG | -24 | 32 | 48 | 4.89 |
| PCC | -2 | -50 | 24 | 4.68 |
| L_dlPFC | -46 | 6 | 30 | 4.55 |
| L_mINS | -40 | -2 | 0 | 4.39 |
| *Note.* | | | | |

| **Supplementary Table 14A.** Peak Coordinates of the **CN13 - Social Representation Network** | | | | | |
| --- | --- | --- | --- | --- | --- |
| Regions | MNI Coordinates | | | Peak intensity  (Z-score) | Cluster size  (mm^3^) |
|  | x | y | z |  |  |
| L_pMTG | -58 | -12 | -14 | 11.68 | 8200 |
| R_pMTG | 60 | -8 | -16 | 8.78 | 7184 |
| L_lOFC | -46 | 28 | -10 | 12.01 | 6640 |
| dmPFC | 4 | 58 | 24 | 8.25 | 4464 |
| R_IFG | 56 | 28 | 6 | 10.81 | 4184 |
| R_AG | 54 | -50 | 22 | 9.15 | 3384 |
| vmPFC | 0 | 46 | -16 | 7.35 | 2176 |
| L_AG | -52 | -58 | 22 | 9.35 | 2064 |
| PCC/PCUN | 0 | -54 | 34 | 8.77 | 1880 |
| preSMA | -4 | 18 | 54 | 7.73 | 1744 |
| L_AMY | -22 | -6 | -18 | 8.06 | 1448 |
| L_PreCG | -42 | 0 | 54 | 6.17 | 648 |
| R_lOFC | 34 | 22 | -14 | 5.57 | 624 |
| R_pMTG | 62 | -24 | -10 | 5.42 | 256 |
| L_THAL | -6 | -12 | 2 | 4.48 | 240 |
| L_CrusII | -24 | -78 | -36 | 5.11 | 200 |
| R_AMY | 24 | -6 | -18 | 4.18 | 176 |
| R_CrusI | 28 | -78 | -34 | 4.91 | 176 |
| L_FFG | -42 | -52 | -16 | 4.33 | 176 |
| L_PMC | -38 | 4 | 36 | 4.89 | 160 |
| R_iLOC | 52 | -72 | 8 | 4.79 | 152 |
| *Note.* | | | | | |

| **Supplementary Table 14B.** Peak Coordinates of the **CN13 - Social Representation Network** (with Subpeaks) | | | | |
| --- | --- | --- | --- | --- |
| Regions | MNI Coordinates | | | Peak intensity (Z-score) |
|  | x | y | z |  |
| L_lOFC | -46 | 28 | -10 | 12.01 |
| L_pMTG | -58 | -12 | -14 | 11.68 |
| R_IFG | 56 | 28 | 6 | 10.81 |
| L_pMTG | -56 | -42 | 4 | 9.94 |
| L_IFG | -50 | 26 | 6 | 9.44 |
| L_AG | -52 | -58 | 22 | 9.35 |
| R_AG | 54 | -50 | 22 | 9.15 |
| L_TP | -52 | 4 | -28 | 8.89 |
| R_pMTG | 60 | -8 | -16 | 8.78 |
| PCC/PCUN | 0 | -54 | 34 | 8.77 |
| R_pMTG | 52 | -36 | 1 | 8.51 |
| dmPFC | 4 | 58 | 24 | 8.25 |
| R_TP | 50 | 10 | -28 | 8.25 |
| dmPFC | 4 | 58 | 24 | 8.25 |
| dmPFC | -6 | 54 | 34 | 8.23 |
| L_IFG | -50 | 20 | 18 | 8.17 |
| L_AMY | -22 | -6 | -18 | 8.06 |
| R_aMTG | 54 | 0 | -22 | 7.93 |
| preSMA | -4 | 18 | 54 | 7.73 |
| vmPFC | 0 | 46 | -16 | 7.35 |
| R_IFG | 54 | 24 | 24 | 7.09 |
| R_IPL | 54 | -64 | 24 | 6.25 |
| L_PreCG | -42 | 0 | 54 | 6.17 |
| R_lOFC | 34 | 22 | -14 | 5.57 |
| R_pMTG | 62 | -24 | -10 | 5.42 |
| L_mPFC | -8 | 52 | 0 | 5.31 |
| L_CrusII | -24 | -78 | -36 | 5.11 |
| R_CrusI | 28 | -78 | -34 | 4.91 |
| L_PMC | -38 | 4 | 36 | 4.89 |
| R_iLOC | 52 | -72 | 8 | 4.79 |
| L_THAL | -6 | -12 | 2 | 4.48 |
| L_FFG | -42 | -52 | -16 | 4.33 |
| R_AMY | 24 | -6 | -18 | 4.18 |
| *Note.* | | | | |
